# Human cancer genomes harbor the mutational signature of tobacco-specific nitrosamines NNN and NNK

**DOI:** 10.1101/2024.06.28.600253

**Authors:** Michael Korenjak, N. Alpay Temiz, Stéphane Keita, Bérénice Chavanel, Claire Renard, Cécilia Sirand, Vincent Cahais, Tanguy Mayel, Karin R. Vevang, Foster C. Jacobs, Jiehong Guo, William E. Smith, Marissa K. Oram, Flaviu A. Tăbăran, Ozan Ahlat, Ingrid Cornax, M. Gerard O’Sullivan, Samrat Das, Shuvro P. Nandi, Yuhe Cheng, Ludmil B. Alexandrov, Silvia Balbo, Stephen S Hecht, Sergey Senkin, Francois Virard, Lisa A. Peterson, Jiri Zavadil

## Abstract

Tobacco usage is linked to multiple cancer types and accounts for a quarter of all cancer-related deaths. Tobacco smoke contains various carcinogenic compounds, including polycyclic aromatic hydrocarbons (PAH), though the mutagenic potential of many tobacco-related chemicals remains largely unexplored. In particular, the highly carcinogenic tobacco-specific nitrosamines NNN and NNK form pre-mutagenic pyridyloxobutyl (POB) DNA adducts. In the study presented here, we identified genome-scale POB-induced mutational signatures in cell lines and rat tumors, while also investigating their role in human cancer. These signatures are characterized by T>N and C>T mutations forming from specific POB adducts damaging dT and dC residues. Analysis of 2,780 cancer genomes uncovered POB signatures in ∼180 tumors; from cancer types distinct from the ones linked to smoking-related signatures SBS4 and SBS92. This suggests that, unlike PAH compounds, the POB pathway may contribute uniquely to the mutational landscapes of certain hematological malignancies and cancers of the kidney, breast, prostate and pancreas.

## INTRODUCTION

Tobacco use, linked to 25% of cancer deaths, is associated with multiple cancer types, including cancers of the lung, oral cavity, pharynx, larynx, esophagus, pancreas, urinary bladder, kidney and liver^1^. Smokeless tobacco contains >4,000 chemicals, including at least 30 compounds linked to cancer^2^. As a product of combustion, tobacco smoke is a mixture of >7,000 compounds^3^, of which 80 cause cancer in laboratory animal models^4^. This chemical diversity helps explain tobacco’s multiple-organ tumorigenic effects. However, the causal roles of most of the specific chemical components present in tobacco in human cancer development are not clear.

*N’*-nitrosonornicotine (NNN) and 4-(methylnitrosamino)-1-(3-pyridyl)-1-butanone (NNK), two tobacco-specific carcinogens, likely contribute to the mutagenic effects of tobacco and tobacco smoke in humans. Formed from tobacco alkaloids during the curing process of tobacco, both are present in smokeless tobacco and tobacco smoke^5^. NNN is the most abundant esophageal carcinogen in tobacco products^5^. In rats, it induces squamous cell esophageal, nasal cavity and oral tumors^5–7^. In smokers, urinary levels of NNN and its glucuronide are strongly associated with esophageal cancer risk^8,9^. NNK is a potent lung carcinogen, which also induces liver and nasal tumors in animals^10–12^. It induces lung adenocarcinomas in rodents at doses comparable to those experienced by smokers^5^.

Together with the well-established metabolic activation of NNN and NNK, resulting in the formation of DNA adducts, and their detection in tobacco users, these observations provided key mechanistic information for the IARC classification of these two tobacco-specific nitrosamines as Group 1 carcinogens^8,13,14^. The NNK metabolite 4-(methylnitrosamino)-1-(3-pyridyl)-1-butanol (NNAL) is detected in the urine of all smokers, and its levels are significantly related to increased lung cancer risk, indicating a specific role for NNK^15–17^. Similarly, the relationship of NNN to esophageal cancer in smokers has been demonstrated epidemiologically^8^. Specific patterns of molecular damage in human tumors, such as mutational signatures linked to these compounds, have not been identified.

Somatic mutations are important contributors to carcinogenesis, and characteristic mutation patterns attributed to individual mutagenic sources have been mathematically extracted as mutational signatures from many tumor samples^18^. Reference sets of mutational signatures have been compiled in the Catalogue of Somatic Mutations in Cancer (COSMIC)^19^ and the Signal database^20^. They are based on single-base substitutions (SBS), doublet-base substitutions (DBS), small insertions and deletions (indels)^21^, and large-scale genomic structural alterations such as CNV and SV^22,23^. These patterns result from either endogenous processes or environmental exposures or their combined effects^21,24,25^. As numerous mutational signatures are associated with specific mutagenic processes, their presence provides evidence that a given exposure or endogenous activity plays a role in the mutagenic and, possibly, carcinogenic process. In this context, experimentally generated signatures play an important role in revealing the specific underlying causal factors^26,27^.

There are mutational signatures that are associated with tobacco use: SBS4 (smoking)^28^, SB29 (smokeless tobacco use)^29^, and SBS92 (smoking)^20,30,31^. Multiple tobacco smoke chemicals likely contribute to the main features of SBS4, for which benzo[*a*]pyrene (BaP) and other polycyclic aromatic hydrocarbons (PAH) have been cited as likely causes of mutations at guanine residues^32–35^ while SBS4 also contains T>A transversions, with the underlying adenine damage previously linked to the effects of other PAH or to glycidamide, a genotoxic metabolite of the tobacco-smoke chemical acrylamide^32,35^. Recently, additional mutational signatures of yet unknown etiologies with predominant T>A alterations have been linked to tobacco smoking^36,37^. SBS4 further associates with signature ID3, characterized by single base deletions of predominantly C, with signature DBS2, characterized predominantly by CC>AA doublets, and with DBS6, harboring mostly TG>AT and TG>CT. SBS29 is similar to but discernible from SBS4, possibly as PAHs like BaP are less abundant in smokeless tobacco. The individual chemical(s) or processes responsible for SBS29 and SBS92 remain unknown^30,31^.

Here we tested whether NNN and NNK are possible contributors to the tobacco-associated mutational signatures. Both compounds are converted to reactive metabolites that chemically modify DNA and form pyridyloxobutyl (POB) DNA adducts^38^. NNK also forms methyl, pyridylhydroxybutyl and formaldehyde DNA adducts^38^. Due to the complexity of NNK- and NNN-derived DNA adducts, model compounds have been used to characterize the mutagenic properties of the various metabolic pathways. 4-(acetoxymethylnitrosamino)-1-(3-pyridyl)-1-butanone (NNKOAc) models the DNA pyridyloxobutylation pathway of NNK and NNN in cell lines and in laboratory animal models^38^. NNKOAc is converted by esterases to the same α-hydroxymethyl-NNK metabolite that is formed upon cytochrome P450 catalyzed hydroxylation of NNK’s methyl group. It generates the same POB DNA adducts caused by NNK and NNN in rodents and is mutagenic, causing a variety of DNA adducts and discernible mutation patterns ^38^ ^39^.

To test the hypothesis that POB DNA adducts contribute to the carcinogenic properties of NNK and NNN we devised a multi-system, multi-species approach to comprehensively investigate the toxicogenomic effects of these compounds (see overall design in Supplementary Figure 1). The mutational signatures of these chemicals were determined using NNKOAc exposure in human A459 lung cancer cells, normal human oral keratinocytes (NOK) and the Hupki murine embryonic fibroblasts (MEF) as well as in rat esophageal tumors arising due to NNN exposure. Levels of POB DNA adducts were measured in the model systems to identify the DNA adducts likely responsible for the observed signatures. Computational screening of public human pan cancer genome data for the experimentally derived POB signature was then performed to assess its contribution to mutational landscapes of human cancer.

## RESULTS

### NNKOAc induces a characteristic mutational signature in cell-based exposure models

The human cell lines, A549 and NOK, were selected based on the relevance of tobacco-associated tumorigenesis in the source tissue types. MEFs were included as a well-established experimental model system for mutational signatures analysis^33–35,40,41^. To establish how exposure levels affect cell fitness, we monitored cell viability and cell growth rates (Supplementary Figure 2). All cell types exhibited an NNKOAc exposure dose-dependent decrease in viability. The concentration ranges affecting cell viability (or growth) were quite similar for A549 cells and MEFs, while NOK were more sensitive to NNKOAc exposure. NOK and A549 cells were repeatedly exposed to NNKOAc for five weeks and then single cell subcloned for conventional NGS (see Methods). Clones from NNKOAc-exposed primary MEF cells applicable to NGS were generated using a one-time exposure, followed by senescence bypass and immortalization (see Methods).

Whole-exome sequencing (WES) of NNKOAc-exposed NOK, A549 and MEF clones (6, 6 and 4 clones respectively), alongside untreated controls for each cell type, revealed per-sample mutation numbers ranging from the low hundreds to thousands for the exposed clones, while the control clones exhibited fewer single base substitutions (SBS) (Supplementary Table 1). Average mutation numbers among exposed clones were the highest in A549 cells and the lowest in NOK, in line with the lesser sensitivity of A549 cells to NNKOAc treatment, which had prompted us to use higher exposure concentrations in A549 cells as compared to NOK (75 vs 6.25 µM).

From the three cell types, we extracted exposure-specific mutational signatures, supported by the majority of the observed SBSs, likely caused by NNKOAc-induced POB DNA adduct formation (Supplementary Figure 3). Individual extraction of the treatment-specific signatures from the different experimental models shows resemblance between the signatures, which are all characterized by prominent T>A, T>C, T>G and C>T alterations, with transcriptional strand bias for these mutation types on the non-transcribed/coding strand of genes (Figure 1A-C), consistent with the principal mutagenic damage on thymidine and cytidine. Despite their similarity, model system-specific signature differences with respect to the main mutation types and strand-biased mutation classes are discernible in the signature profiles and in the corresponding cosine similarity analysis (Figure 1A-C, I) and may represent differences in adduct formation and/or repair.

**Figure 1.**
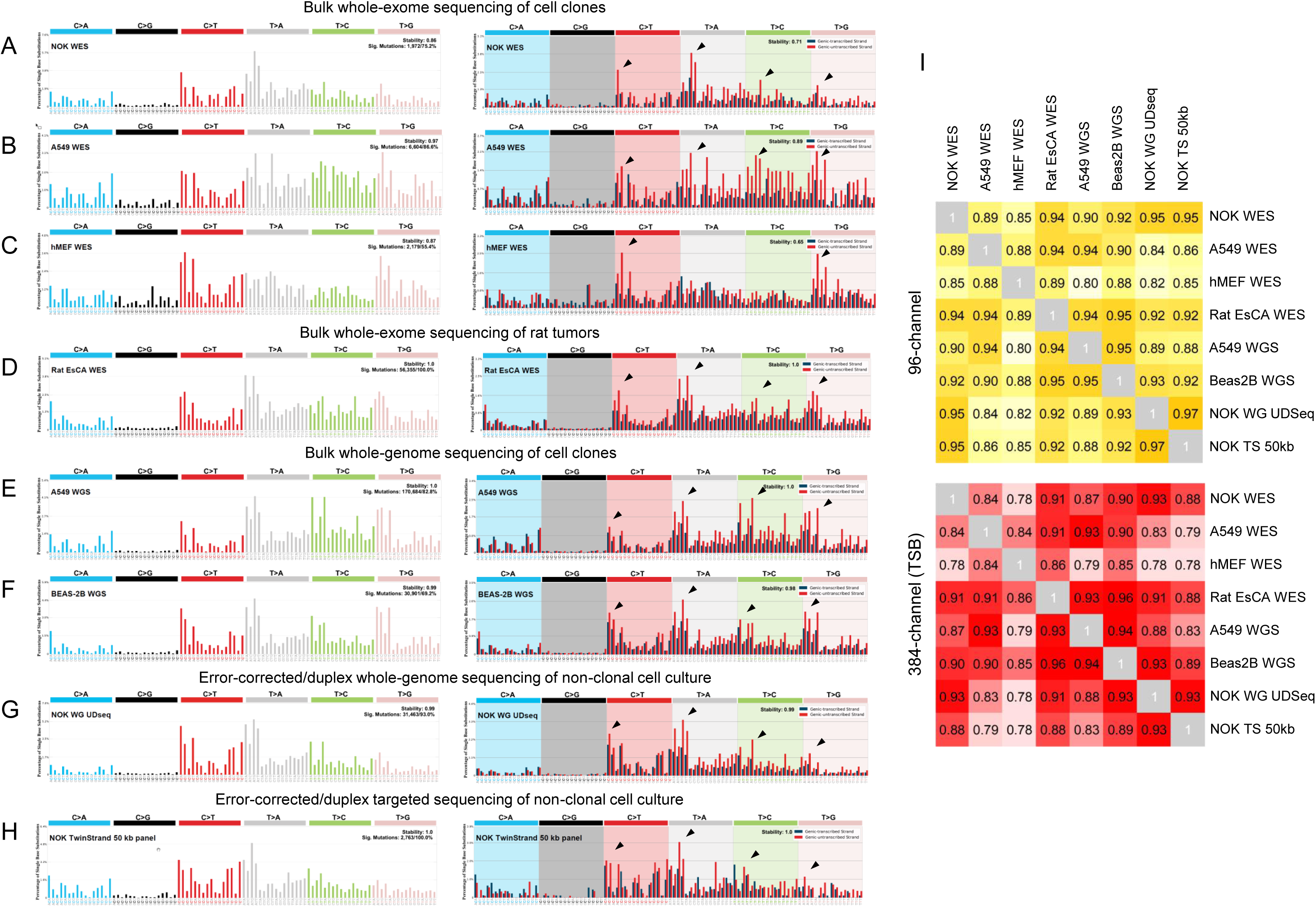
Experimental modeling of the mutational signatures of NNKOAc and NNN. 96-channel signatures (left panels) and transcription strand-based 192-channel signatures (right panels) are shown. **(A-D)** Exome sequencing-based characterization of signatures of NNKOAc exposure in three cell models (NOK, A549, hMEF) and NNN-induced rat esophageal tumors (Rat EsCa). **(E,F)** Conventional whole-genome sequencing-based NNKOAc signature extracted from A549 and BEAS-2B cells. Single base substitution calls from NNKOAc-treated BEAS-2B cells for signature extraction were taken from^39^. **(G)** Whole genome Universal-Duplex sequencing-based NNKOAc signature extracted from NOK cells. **(H)** Targeted Duplex sequencing-based NNKOAc signature extracted from NOK cells. **(I)** Cosine similarity analysis among all experimentally derived mutational signatures (top panel: 96-channel analysis; bottom panel: 384-channel analysis).

We next generated a genome-scale version of the signature from clonally expanded A549 cells using bulk whole genome sequencing (WGS) (Figure 1E). We also reanalyzed the recently reported mutation pattern observed in NNKOAc-exposed human BEAS-2B cells, to obtain its transcription strand bias (TSB) version (Figure 1F). The BEAS-2B data further strengthens the multisystem findings reported in this study. In addition, we applied error-corrected/duplex whole genome and targeted sequencing to NNKOAc-exposed heterogeneous NOK cultures to demonstrate the similarity of POB signatures extracted from cells that have not undergone clonal expansion before sequencing (Figure 1G,H). All cell-based experimental signatures show remarkable similarity, including their TSB.

### The mutational signature of NNN in rat tumors corresponds to the experimental NNKOAc signature

The DNA damage generated by NNKOAc mimics that of the POB pathway upon metabolization of both NNN and NNK. We thus analyzed rat tumors induced by exposure to NNN, to examine the POB signature *in vivo*. We focused on esophageal cancer, which was the most common tumor type induced by NNN, followed by cancers of the pharynx, tongue and oral cavity (Supplementary Table 2). Signature extraction from WES of twelve NNN-induced tumors revealed a signature that is highly concordant with the cell-based signatures (Figure 1D, 2I), similarly characterized by T>N and C>T alterations with transcription strand bias. Overall, the rat tumors show high SBS counts (∼2,000 to >7,500, Supplementary Table 1), indicating strong mutagenicity of NNN, which is further supported by the penetrating effects of the signature in the individual tumors (Figure 2).The putative POB signature (SBS_POB) derived from rat esophageal tumors exhibited high similarity with most other experimental signatures, and the genome-scale NOK and A549 SBS_POB signatures showed the highest cosine similarity with the exome-based signatures generated in the same cell line (Figure 2I). Moreover, higher similarity was observed between signatures derived from the same cell type (A549 and BEAS-2B lung cells *vs* NOK), while the signature from mouse fibroblasts was less similar to the other signature patterns.

**Figure 2.**
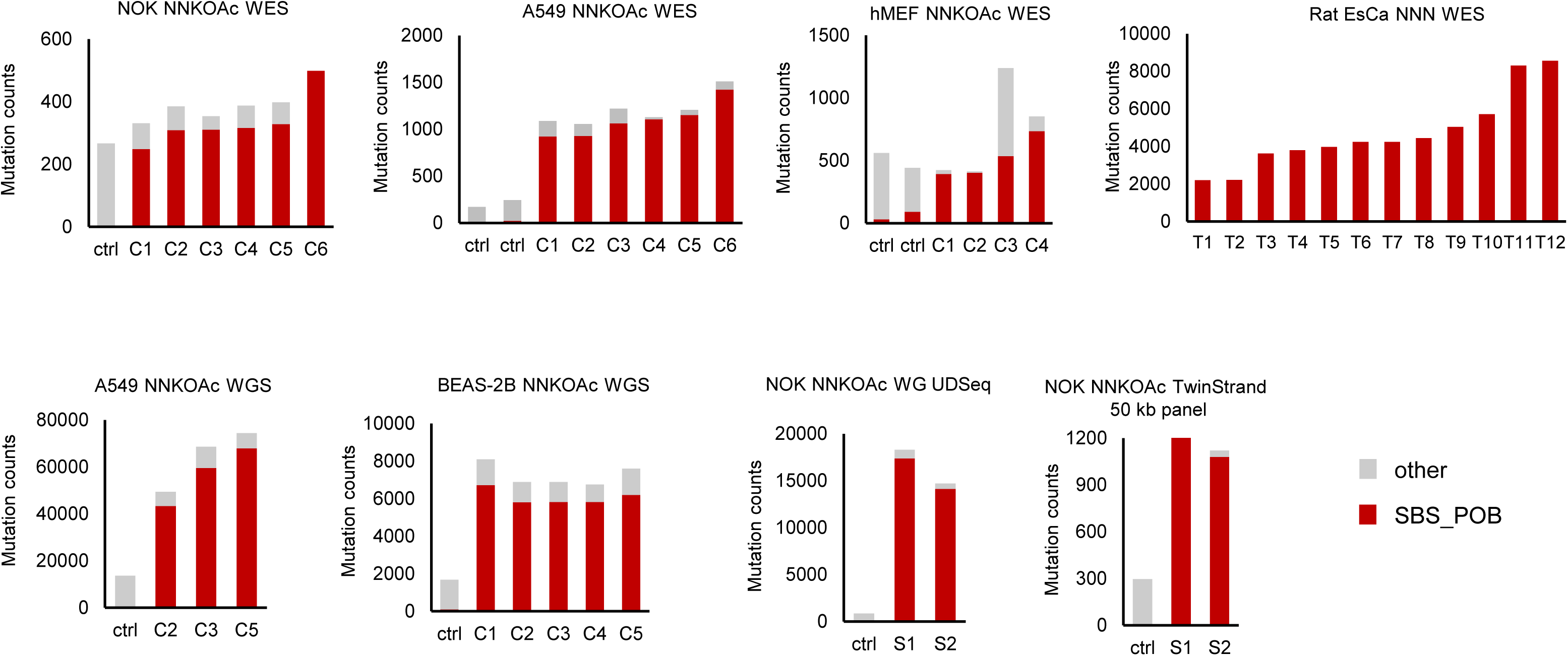
Total mutation count attributed to the extracted experimental SBS_POB and other SBS mutational signatures in cells and rat tumor samples. WES – whole exome sequencing; WG UDSeq – whole-genome Universal Duplex sequencing; WGS – conventional whole-genome sequencing; ctrl – untreated control; C – clone from exposed cells; S – bulk cell sample; T – tumor sample

Mutations attributed to the experimentally derived SBS_POB signature accounted for the vast majority of mutations in the exposed samples (Figure 2). Importantly, SBS_POB was not detected in the untreated control cultures. The findings confirm the clear separation of the exposure-specific mutation patterns from any background biological or cell culture-derived processes present in the untreated control cells. Contribution of such background processes are moderately observed in the context of MEFs (Supplementary Figure 4A), for which characteristic, intrinsic mutation patterns defined by C>G alterations in a 5’-G<u>C</u>C-3’ sequence context (the mouse signature mSBS_N3)^42^ and matching COSMIC SBS17 have been described^33,43^. In addition, A549 cells harbor a background mutational signature linked to oxidative stress induced DNA damage with similarity to SBS36 (Supplementary Figure 4B).

Next, we carried out targeted DNA adduct analyses of A549 and BEAS-2B cells and rat esophageal tissue using mass spectrometry (Figure 3, Supplementary Tables 3 and 4) to confirm the formation of POB DNA adducts under the exposure conditions used in the sequencing and mutational signature analysis. O^2^-POBdT and N^7^-POBG were readily detected in NNKOAc-treated A549 and BEAS-2B cells, and in NNN-treated rat tissues, with adduct levels increasing in an exposure concentration-dependent manner (Figure 3B,C). The adduct species were not detected in untreated control cells and animals. The O^6^-POBdG adduct was detected at considerably lower levels in the cell and rat tissue samples, suggesting that it was repaired more efficiently. The NNN dose-dependency of POB DNA adduct formation in esophageal tissue is further reflected in the concentration-dependent increase in mutation loads observed in tumors (Figure 3D). The presence of the O^2^-POBdT DNA adduct reflects the predominant T>N mutations of the experimentally derived POB mutational signature and suggests a convergent (pre)mutagenic mechanism attributed to both NNN and NNK.

**Figure 3.**
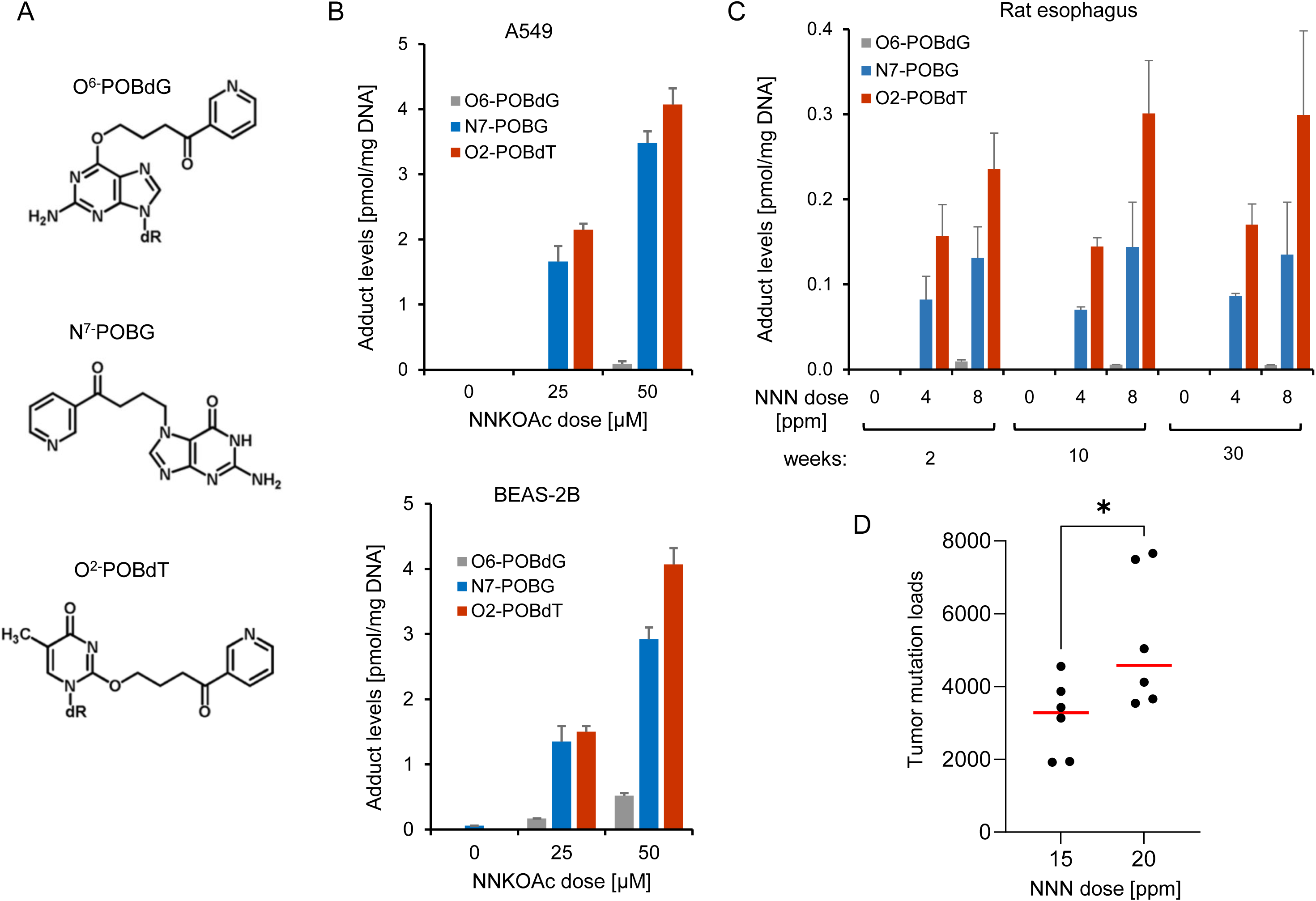
POB DNA adducts form in cells and rat tissue. (A) Structures of the analyzed pyridyloxobutyl (POB) DNA adducts. **(B)** Levels of POB DNA adducts in NNKOAc-exposed A549 and BEAS-2B cells. Cells were treated with the indicated concentrations of NNKOAc for 24 hrs. **(C)** Levels of POB DNA adducts in rat esophageal tissue from NNN-treated animals, collected after 2, 10 and 30 weeks of exposure to the indicated concentrations of NNN. **(D)** Mutation loads of NNN-induced rat esophageal tumors. Mutation numbers are based on WES. The mutation load comparison between tumors induced by different NNN exposure concentrations (15 *vs.* 20 ppm) is shown. Significance: *=p<0.05, Mann-Whitney U test.

### SBS_POB mutational signature is observed in human pan-cancer genome data

Next, we investigated the ICGC’s Pan-Cancer Analysis of Whole Genomes Consortium (PCAWG) collection of 2,780 human cancer genomes for the presence of the SBS_POB signature. For this, we assembled a curated set of 54 strand-biased signatures from the reference set of COSMIC mutational signatures (see Methods). The whole genome SBS_POB signature derived from A549 cells, used for screening, did not exhibit substantial cosine similarity with any of the 54 COSMIC mutational signatures (max. similarity 0.71) (Supplementary Table 5).

Using optimized signature refitting analysis within the Mutational Signature Attribution (MSA) tool^44^, which uses the non-negative least square (NNLS) approach, we determined the presence of the 54 COSMIC mutational signatures and either of the A549-derived SBS_POB signature or the negative control signatures (one with scrambled mutation types, another with inverted TSB, see details in Methods and Supplementary Figure 5), in the collection of 2,780 individual ICGC PCAWG tumors. The PCAWG set comprised of 37 tumor types of which 25 have been epidemiologically associated with tobacco use (Supplementary Table 6). The screen of the target PCAWG data was performed at the TSB signature level. As TSB is a prominent feature of the SBS_POB signature, this step ensured high attribution efficiency and specificity.

Upon correction for the negative control signature presence, the SBS_POB signature was observed at an attribution level >=3% in 180 PCAWG samples from 22 tumor types. 165 of the 180 (90.5%) SBS_POB-positive samples were from 16 cohorts of tumor types associated with tobacco smoking (Figure 4 and Supplementary Table 7). The relative proportion of the SBS_POB signature among other signatures was 6.6% on average (median 5.8%, range 3-22.3%). The confidence of the attribution process was established by a simulation using 2,780 synthetic genomes mimicking the cohort and signature compositions observed in the real PCAWG data, with the controlled admixture of low-level presence (average 5.7%, median 3.3%, range 2.3-24.3%) of the SBS_POB signature. The false discovery rate (FDR) of 0.0128 determined by the simulation indicated high confidence of the optimized signature attribution process (see Methods, Supplementary Tables 8 and 9).

**Figure 4.**
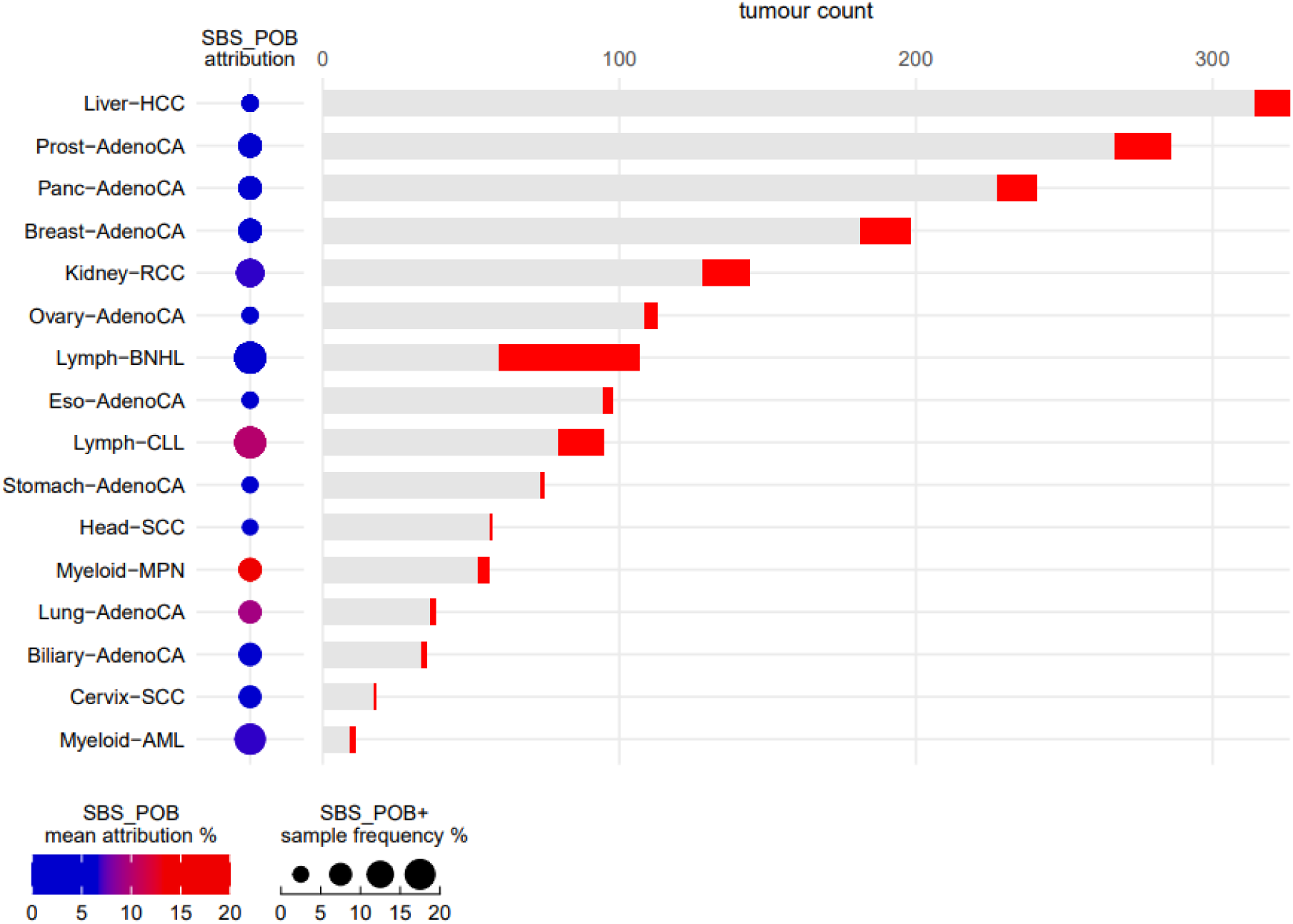
Presence of the SBS_POB signature in human pan-cancer genome data. The SBS_POB-positive samples (red) among the ICGC PCAWG Cohorts are shown as bar graphs. The signature was attributed by the optimized NNLS algorithm within the Mutational Signature Attribution (MSA) tool. The SBS_POB mean attribution percentage and the proportion of positive samples for each cohort are indicated on the left. ICGC PCAWG – International Cancer Genome Consortium Pan Cancer Analysis of Whole Genomes, NNLS – Non-negative least squares.

The proportion of SBS_POB positive samples ranged between 2 and 45%, depending on the tumor (sub)type. The SBS_POB signature was most commonly detected in tobacco-associated hematological disorders (Myeloid-AML, Myeloid-MPN, Lymph-BNHL, Lymph-CLL), renal cell carcinoma, prostate, pancreas and breast cancer. Cancers harboring previously characterized tobacco signatures (SBS4, SBS92), on the other hand, exhibited lower relative SBS_POB presence (lung, head and neck, esophagus, liver, bladder).

### Cancer site-specific distribution of tobacco signatures SBS_POB, SBS4 and SBS92

To help understand the relationship between SBS_POB, SBS4 and SBS92, we next investigated the overall presence of the tobacco smoking-related signatures in the SBS_POB-positive samples. Only 3 out of 165 (2%) SBS_POB-positive samples in tobacco-linked cancers overlapped with 228 SBS4-positive PCAWG tumors, and 10 (6%) SBS_POB-positive tumors overlapped with the set of 187 SBS92-positive tumors. The cancer types that were proportionately most enriched for SBS_POB (above 5% samples per cohort, Supplementary Table 10) and devoid of SBS4 and SBS92, were B-cell non-Hodgkin lymphoma (48 of 107 samples, 45%), acute myeloid leukemia (AML) (2 of 11 samples, 18%), chronic lymphocytic leukemia (CLL) (16 of 95 samples, 17%), clear cell renal cell carcinoma (ccRCC) (13 of 144 samples, 9%), breast adenocarcinoma (17 of 198, 9%), prostate adenocarcinoma (18 of 286 samples, 6%), pancreatic adenocarcinoma (14 of 241, 6%), myeloid myeloproliferative neoplasms (3 of 56 samples, 5.4%), lung adenocarcinoma (2 of 38 samples, 5.4%). In addition, the SBS_POB signature was observed at lower per-sample proportions, in 4 of 113 (3.5%) cases with ovarian adenocarcinoma, 3 of 98 (3%) cases with esophageal adenocarcinoma, 1 of 35 (3%) biliary adenocarcinoma, 2 of 75 (3%) stomach adenocarcinoma, 7 of 326 (2%) hepatocellular carcinoma (HCC) and 1 of 57 (2%) head and neck SCCs.

In contrast and as expected, the tumor types most enriched for SBS4 only were liver HCC (81 of 326, 25%), lung adenocarcinoma (7 of 38, 18%), biliary adenocarcinoma (2 of 35, 6%) and ccRCC (7 of 144, 5%). SBS92 only was observed mainly in bladder urothelial carcinoma (9 of 23, 39%), liver HCC (42 of 326, 13%), esophageal adenocarcinoma (8 of 98, 8%), head and neck SCC (3 of 57, 11%) (Supplementary Table 10).

Based on the presence of SBS_POB, SBS4 and SBS92-positive samples, or lack thereof, three groups of cancer types could be distinguished. Cancers positive for SBS_POB but mostly lacking SBS4 and SBS92 included B-cell non-Hodgkin lymphoma, CLL, AML, myeloid myeloproliferative neoplasms and pancreatic adenocarcinoma (Figure 5A, Supplementary Figure 6A). A second group of cancers was positive for SBS_POB in conjunction with SBS4 or SBS92, or both (ccRCC, prostate, breast, ovary and esophagus adenocarcinoma), although the number of samples harboring more than one signature was limited (Figure 5B, Supplementary Figure 6B). The general attribution levels for the three mutational signatures was very similar for the cancer types in these two groups (Figure 5A,B). A third group of cancers was negative for SBS_POB but positive for SBS4 and/or SBS92 (lung squamous cell and adenocarcinoma, HCC, head and neck SCC and bladder urothelial carcinoma) (Figure 5C, Supplementary Figure 6C). Signature attribution levels were generally higher in these cancer types except for SBS4 in bladder cancer and HCC, and SBS92 in lung adenocarcinoma (Figure 5C). In this group, there was a considerable set of samples (*n*=102) that harbored both SBS4 and SBS92. Among these, lung SCC predominated (44 of 48 samples, 92%), followed by lung adenocarcinoma (15 of 38, 39.5%), liver HCC (37 of 326, 11%), and head and neck SCC (4 of 57, 7%) (Supplementary Figure 6C). Collectively, these results suggest that there is a remarkable tissue-specificity for the putative mutagenic effects of POB, involving tissues and cell types in which SBS4 and SBS92 (both presumed to originate from the PAH compounds’ effects) are generally not operative.

**Figure 5.**
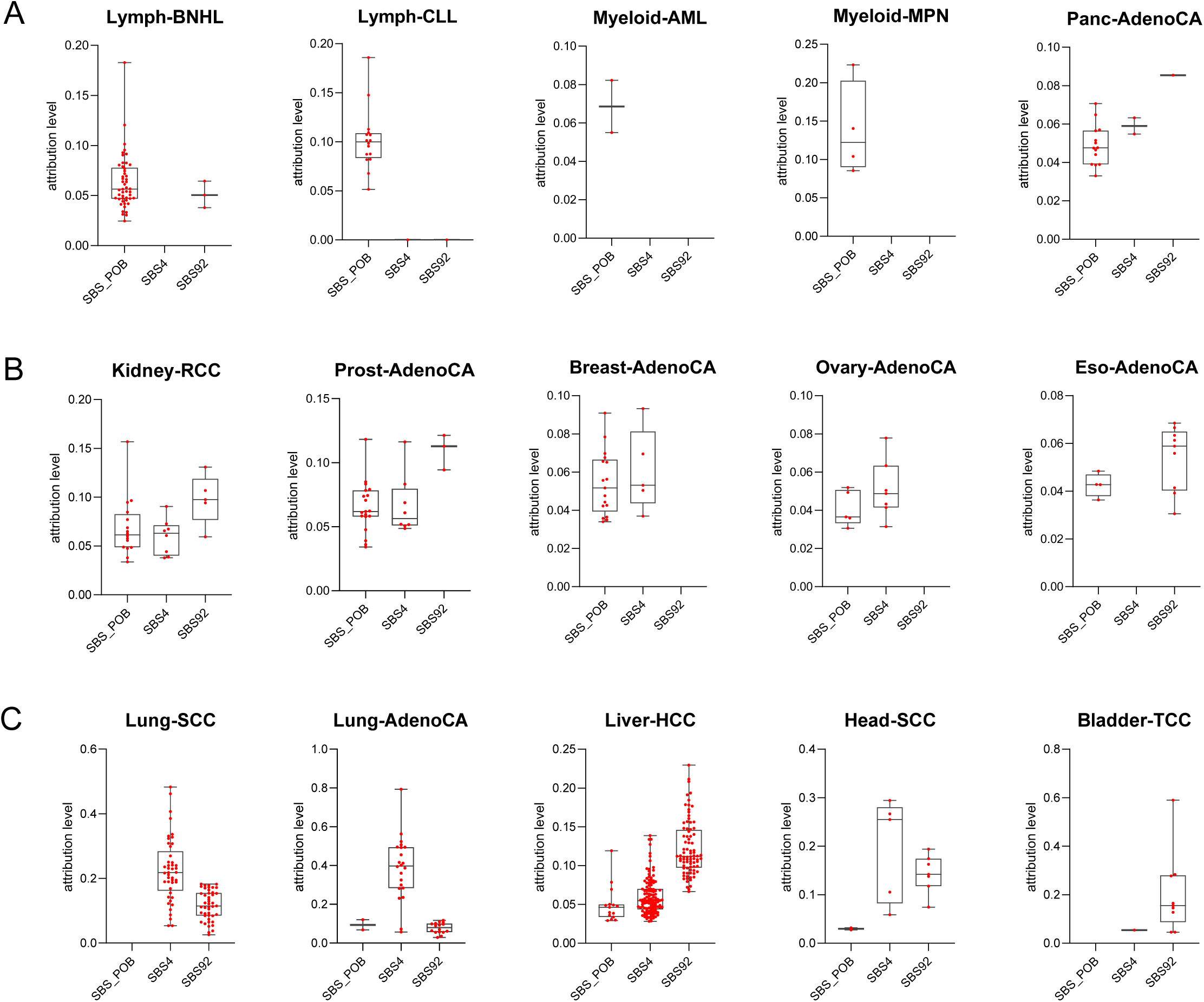
SBS_POB, SBS4 and SBS92 presence in tobacco-linked ICGC PCAWG cohorts. The signatures were attributed by the optimized NNLS algorithm within the Mutational Signature Attribution (MSA) tool. For each cohort and mutational signature, the attribution level of positive samples is shown as bar graphs. **(A)** Cohorts with SBS_POB presence, lacking SBS4 and SBS92. **(B)** Cohorts with presence of SBS_POB, SBS4 and SBS92. **(C)** Cohorts with presence of SBS4 and SBS92, lacking SBS_POB. ICGC PCAWG – International Cancer Genome Consortium Pan Cancer Analysis of Whole Genomes, NNLS – Non-negative least squares.

## DISCUSSION

The experimentally modelled mutational signatures presented in this study show similar SBS profiles, trinucleotide contexts and transcription strand bias between all model systems and are likely the result of POB DNA adduct formation. The transcription strand bias of the signature, which had not been described in a previous study^39^, is very pronounced and suggests DNA damage on thymidines and cytidines as the cause of the signature. NNKOAc and NNN both result in the formation of POB DNA adducts^38^, which were readily detected in the studied samples. O^2^-POBdT has been shown to primarily cause T>A mutations in human cells, as well as a smaller number of T>G and T>C alterations^45^. The O^2^-POBdT adducts result in blocked replication forks that are resolved by error-prone translesion DNA polymerases, and knockdown of which considerably limits the number of T>A changes^45,46^. These observations are in line with O^2^-POBdT being the predominant POB DNA adduct detected in our samples as well as with the major contribution of T>N alterations to the identified mutational signature, suggesting that O^2^-POBdT is the major pre-mutagenic POB DNA adduct in the cell-based and animal models. We speculate that the prominent C>T component of the signature may be the result of O^2^-POBdC adducts, which have been detected in *in vitro* NNKOAc-treated DNA^47^. The cytidine adducts explain the transcription strand bias of the signature, but they are thermally less stable than the thymidine adducts of POB^47^.

Decomposition of the strand-biased versions of the experimentally derived SBS_POB patterns into known COSMIC content could not explain the observed SBS_POB signature. Moreover, none of the experimentally characterized mutational signatures to date, including those of well-established tobacco smoke carcinogens, show high similarity with the SBS_POB signature^32^. This strongly suggests that the signature is unique and has not been previously identified by *de novo* signature extraction from human cancer genome sequencing data.

*In silico* signature attribution revealed the presence of the comprehensively characterized experimental SBS_POB signature, and established tobacco signatures, in human cancer genomes. Among the established tobacco smoking-associated signatures, SBS4 was mostly detected in lung and liver cancer, while SBS92 was predominantly present in bladder cancer. Moreover, our data analysis demonstrated that the majority of PCAWG lung cancer samples harbored both SBS4 and SBS92. This is the first time this extensive co-occurrence of the signatures has been shown. The attribution results for SBS4 and SBS92 corroborate the previously reported presence of the signatures in these cancer types^20,21,28,30,31^, underscoring the confidence in the attribution approach. The general absence of the SBS_POB signature in samples enriched for SBS4 and SBS92 is surprising, but it may indicate that the PAH-related mutation types underlying SBS4 and SBS92 in certain cancer types out compete SBS_POB, or that there is another unknown mechanism that restricts the manifestation of POB adduct-mediated mutagenesis to smoking-related cancer types lacking SBS4 and SBS92. Instead, SBS_POB is enriched in hematological malignancies and pancreatic adenocarcinoma that do not harbor SBS4 and SBS92, raising the possibility of a remarkable tissue specificity for the putative mutagenic effects of POB. Although no data are available for POB DNA adducts in blood, both the NNK metabolite NNAL and NNN have been detected in plasma of tobacco users^15,48^. A detailed history of smoking was available from the ICGC PCAWG records for only 10% (18 of 180) of SBS_POB-positive cases limiting the possibility to link the signature with smoking on a case-by-case basis and highlighting the need to improve the tobacco exposure information in future sample collections.

It is conceivable that certain COSMIC signatures, present in the cancer genome sequencing data, interfere with the attribution of the SBS_POB signature. In our analysis, B-cell non-Hodgkin lymphoma (BNHL) and CLL are the cancer types with the largest contribution of the SBS_POB signature. These two cancer types have the largest proportion of samples harboring SBS9^21^, one of the COSMIC signatures showing a more pronounced cosine similarity with SBS_POB (Supplementary Table 5). SBS9 and the related SBS85 have been linked to somatic hypermutation in lymphoid cells and activation-induced cytidine deaminase (AID) activity, and they are also characterized by T>N alterations with strand bias on the untranscribed strand. Hence, we carried out correlation analysis of the SBS_POB signature with all COSMIC signatures in the studied cohorts. An increased correlation between the presence of SBS_POB and SBS9 in BNHL or CLL was not observed (Supplementary Figure 7), suggesting that the mutational events associated with the SBS_POB signature can be differentiated from those corresponding to SBS9. In contrast to acute myeloid leukemia, for which smoking is an established risk factor, the epidemiological association of BNHL and CLL with tobacco smoking is less clear. Interestingly, an elevated odds ratio for follicular NHL, which makes up around 20% of BNHL cases, was observed in smokers compared to never smokers and it is associated with smoking intensity^49,50^.

Signature attribution provides a powerful complementary strategy to established, de novo signature extraction approaches. Depending on the characteristics of the studied signature, such as trinucleotide mutation contexts and overall signature presence, however, interference of similar signatures that may result in misattributions need to be carefully considered when interpreting findings from signature attribution.

The identification of the SBS_POB signature in the cancer genome data using an attribution strategy is analogous to what we previously reported for the signature of the acrylamide metabolite glycidamide^35^, suggesting that the signatures’ dispersed mutation patterns may have escaped identification in standard mutational signature extraction analyses. It is also possible that, in humans, other POB DNA adducts contribute to NNN and NNK’s mutational profile due to differences in DNA damage repair, or other pathways of DNA adduct formation may contribute more to the human mutation patterns caused by tobacco-specific nitrosamines (TSNA) exposure. For example, NNK forms methyl and pyridylhydroxylbutyl (PHB) DNA adducts, in addition to the POB DNA adducts^5,38^. In addition, it generates formaldehyde and 4-oxo-4-(3-pyridyl)-1-butanone (OPB), which can also modify DNA^38^. NNN also generates adducts resulting from 5′-hydroxylation^51–53^. In contrast to experimental studies, evidence for the presence of NNN- and NNK-derived POB DNA adduct formation in human lung tissue, and its association with smoking, is limited and based on a small number of studies, which assessed 4-hydroxy-(3-pyridyl)-l-butanone (HPB)-releasing DNA adducts that can result from tobacco-specific nitrosamines^5,54,55^. While levels of HPB-releasing DNA adducts were higher in smokers with lung cancer than smokers without lung cancer^56^, analysis of HPB release in non-cancerous human lung tissue from smokers and non-smokers resulted in inconsistent findings, and a clear association of tobacco-specific nitrosamine DNA adducts with cigarette smoking could not be established. This is also reflected in the IARC Monographs evaluation of NNN and NNK, both of which were classified as Group 1 carcinogens based on strong evidence in animals and mechanistic studies, while evidence for their carcinogenicity in humans was inadequate, as most data were associative, and it was not possible to exclude the potential confounding effect of other tobacco smoke chemicals^14,57^.

The SBS_POB signature adds to a growing number of mutational signatures introduced by compounds present in tobacco products and extracted based on controlled exposure studies and next-generation sequencing-based mutation analysis^32–35,39,40,58,59^. The findings provide valuable insight into the contribution of individual chemicals to the mutation landscapes associated with tobacco consumption, including the presence of the SBS_POB signature in human pan-cancer genome data. The detection of the signature in hematological cancers and tumors of the kidney, breast, prostate and pancreas raises important questions regarding the potential role of certain tobacco smoke chemicals in tobacco-linked cancer types lacking SBS4 and SBS92, while the lower presence of some experimentally derived signatures, such as the SBS_POB signature, in human cancer genome sequencing data highlights possible limitations associated with current signature extraction and attribution strategies. The study provides the first report of the presence of a TSNA mutational signature in human cancers, and it emphasizes the need for additional cancer genome sequencing studies, the analysis of organ- and cell type-specific variations of the SBS_POB signature and prospective sample collections with comprehensive exposure information for a wide range of tobacco products.

## MATERIALS AND METHODS

### Cell culture

The A549 lung cell line was purchased from American Type Culture Collection (ATCC). hTERT-immortalized normal oral keratinocytes (NOK) were kindly provided by Dr. Paul Lambert (University of Wisconsin, Madison). <u>Hu</u>man <u>p</u>53 <u>k</u>nock-in (Hupki) primary mouse embryonic fibroblasts (MEFs) were a gift from Dr. Monica Hollstein (University of Leeds, UK). All cells were grown in standard conditions at 37°C and with 5% CO2. A549 cells were grown in MEM (Gibco, Thermo Fisher Scientific), supplemented with 5% fetal calf serum (FCS), 1% glutamine, 1% penicillin/streptomycin. NOK were grown in Keratinocyte Growth Medium 2 (Promocell) with 1% penicillin/streptomycin. MEF were maintained in Advanced DMEM (Gibco, Thermo Fisher Scientific) with 15% FCS, 2% sodium pyruvate and 1% penicillin/streptomycin.

### Chemicals for cell treatments

Acetoxymethyl-NNK (NNKOAc) was purchased from Santa Cruz Biotechnology (sc-206764). Ten mg were dissolved in dimethyl sulfoxide (DMSO) to obtain a 100 mM stock solution. 10 µL and 20 µL aliquots of this stock solution were prepared and stored at -80°C. This stock solution was then diluted in cell culture medium to obtain the desired exposure concentration.

### MTS assay

The experiment was performed in 96-well plates. A549 cells were seeded at a concentration of 3,000 cells per well, NOKs at 5,000 cells per well, and MEFs were seeded at 10,000 cells per well. In addition to the NNKOAc exposure conditions, two control conditions were included: untreated cells and DMSO treated cells. The DMSO concentration used corresponded to the amount of DMSO in the highest tested NNKOAc concentration. The exposures for the MTS assay were carried out for 24 hours or 48 hours, depending on the cell type. At the end of the exposure, cells were washed with PBS and incubated in regular medium for 48 hours before the absorbance reading. This setup was chosen to account for delayed exposure effects on cell fitness. The MTS assay (CellTiter 96 AQueous One solution cell proliferation assay, Promega -Ref G3580) was performed by a medium change adding 100 μL of cell culture medium containing 10% MTS solution to the wells, incubating for 3 hours in the incubator and then recording the absorbance at 490 nm with a 96-well plate reader (Apollo 11 - LB913, Berthold Technologies). Each condition was run in triplicate and the absorbance relative to untreated control cells was analyzed.

### Exposure of human cell lines

Exposure of human cell lines was performed in 6-well plates. A549 cells and NOK were seeded at a concentration of 200,000 cells per well. NNKOAc was added to the cells while untreated or mock treated cells were kept as controls. Five exposure rounds were carried out for A549 and NOK. A round of exposure was performed every week. Cells were seeded on day 1, the exposure started on day 2 for 24 (A549) or 48 hours (NOK). During 48 hour exposures, NNKOAc was refreshed after 24 hours. After ending the exposure, cells were subcultured on day 4 or 5 of the experimental cycle depending on their density. Based on the MTS results, A549 cells were exposed to 75 µM NNKOAC during the first exposure round, while NOK were treated with 6.25 µM NNKOAc, inducing 30-40% cytotoxicity. The exposure concentrations were adjusted for the subsequent exposures based on visual inspection of the cells after each round (A549 exposure range: 25 to 75µM; NOK exposure range: 6.25 to 12.5µM). NNKOAc exposure resulted in considerably reduced cell growth compared to controls in both A549 cells and NOK (Supplementary Figure 2D,E), probably as a result of reduced cell viability. However, the exposed cells continued to grow throughout the experiment, which is a prerequisite for fixing NNKOAc-induced DNA damage into mutations. At the end of the exposure regimen, cultures were single-cell subcloned.

### Single-cell subcloning

Single-cell subcloning was carried out in a 96-well plate, seeding “0.5 cells per well”, to obtain a theoretical seeding of one cell in half the wells of the plate. A549 cells were cultured in regular growth medium during early stages of single-cell subcloning, while NOK were grown in base medium supplemented with 1% serum. All single-cell subcloning experiments were carried out in an incubator with 5% O2, to reduce the effect of oxidative stress on cell survival. After four to five days of incubation, the plates were screened for wells with individual clones under a microscope. Once dense, the cells were transferred to larger well plates, and later to flasks at which point they were frozen.

### MEF exposure and immortalization

MEF exposure and immortalization were carried out according to an established protocol^35^. Primary Hupki MEFs were subjected to a one-time acute treatment with NNKOAc for 48 hours, using concentrations that had been determined with the help of the MTS assay. Independent primary MEF cultures (untreated or exposed) were grown to senescence and maintained until they immortalized (in a clonal manner), as evident in the characteristic growth curves of MEF immortalization (Supplementary Figure 2F). Two independent clones were generated for each condition (DMSO control, NNKOAc low – 75µM, NNKOAc high – 100µM).

### DNA extraction, NGS library preparation and sequencing of cell lines

DNA was extracted from 1×10^6^ cells using a standard protocol (Macherey-Nagel Nucleospin). Sequencing library preparation was performed using the KAPA HyperPlus Library Preparation kit and associated protocol. Optimal library yields and fragment size distribution were confirmed using a Qubit Fluorometer and an Agilent Tapestation 4200, respectively, at various stages of the protocol. For whole-exome sequencing, target regions were selected by hybridization-based target enrichment (Roche, SeqCap EZ) after library preparation. Sequencing was performed on the Illumina NextSeq 500 platform in paired-end 75-bp runs, to achieve the target ∼50 million paired reads per sample. DNA from six NNKOAc-exposed A549 clones were sequenced alongside two untreated control clones and the parental cells. For NOK, exome sequencing of six NNKOAc-treated clones, one untreated control clone and the parental cells was carried out. Four immortalized, exposed MEF clones and two untreated, spontaneously immortalized clones, together with parental (primary) MEFs were sequenced. Whole-genome sequencing was performed on the Illumina Novaseq 6000 platform (S4 flow cell) in paired-end 150-bp runs, loading six samples per lane to achieve the target ∼400 million paired reads per sample. Three NNKOAc-treated A549 clones, one untreated clone and the parental culture were processed for whole genome sequencing. All exome- and genome-sequenced samples from this study are summarized in Supplementary Table 11.

### Cell culture for DNA adducts exposure

A549 cells were grown in Minimum Essential Media Eagle medium (Gibco, Waltham, MA) with 5% fetal bovine serum, 1% penicillin/streptomycin, and 1% L-glutamine. BEAS-2B cells were grown in BEGM complete media (Lonza, Basel, Switzerland) in vessels coated with a mixture of fibronectin, bovine serum albumin, and collagen. Cells were grown in a humidified atmosphere with 5% CO2 at 37 °C. Cells were subcultured prior to confluence and maintained at no more than 20 passages. Cells were split using 0.5% trypsin with 0.2% EDTA (A549; Sigma Aldrich, Burlington, MA) or 0.25% trypsin with 0.5% added polyvinylchloride (BEAS-2B; ATCC, Manassas, VA) and were routinely tested for mycoplasma using a commercially available surveillance kit (Mycoscope, Genlantis, San Diego, CA). Triplicate T75 flasks were seeded with 1.5 × 10^6^ A549 cells A549 or 2.25 × 10^6^ BEAS-2B 24 h prior to exposure. Cells were treated with 0, 25 or 50 µM NNKOAc for 24 h. Cell monolayers were washed with PBS, trypsinized and harvested by centrifugation and then repelleted for DNA isolation. Pellets were stored at -80°C until analysis.

### Animal treatments

This study was approved by the University of Minnesota Animal Care and Use Committee. NNN was purchased from Toronto Research Chemicals (North York, ON) and purity was confirmed by NMR analysis. Male F344 rats (6 weeks old) were purchased from Charles River Laboratories (Kingston, NY) and housed 2 per cage and allowed to acclimate for 1 week prior to the start of the experiment. There were two experiments. In the first experiment, groups of 24 rats were treated with 0, 4 or 8 ppm NNN in the drinking water. Three rats per group were euthanized by CO2 overdose at 2, 10 or 30 weeks. Tissues were harvested and stored at -80 C for DNA isolation. The remaining rats were treated for 90 weeks or until they started losing weight due to esophageal tumors or had other health issues. In the second experiment, groups of 30 rats were treated with 0, 8, 15 or 20 ppm of NNN in the drinking water for up to 90 weeks or until they lost weight or had other health issues. In both experiments, fresh NNN solutions were prepared each week with the water bottles weighed and refilled twice a week. The NNN concentrations in the stock solutions were analyzed by HPLC as previously described^7^. The average concentrations for the solutions over the course of the experiments were 3.8 ± 0.2 and 7.7 ± 0.3 ppm for the first experiment and 7.9 ± 0.2, 14.9 ± 0.4 and 20.0 ± 0.6 ppm for the second experiment. The animals were euthanized with CO2 and necropsy was performed immediately with all rats evaluated for the number of tumors. Major organs and gross lesions were fixed in 10% formalin for histopathological analysis. In the second experiment, only gross lesions were counted. In addition, select tumors and surrounding normal tissue were removed and flash frozen for DNA isolation for whole-exome sequence analysis. Samples were prepared for histopathological analysis by using a commercially available decalcifying solution containing ethylenediaminetetraacetic acid and dilute HCl (Newcomer Supply, Middleton, WI) to decalcify the head, tongue and larynx. After processing fixed tissue specimens into paraffin blocks, they were sectioned to 4 µM thickness and stained with hematoxylin and eosin. Blinded to treatment group, the evaluation of histological slides was performed by two A.C.V.P-certified pathologists using light microscopy to verify diagnoses. A Nikon SMZ 1000 stereomicroscope was used to carefully evaluate the esophagus as well as the tongue, larynx, pharynx, oral mucosa, and soft and hard palate for tumor ≥ 0.5 mm.

### DNA isolation for DNA adduct measurements

DNA was isolated from esophagi or pelleted cells using Qiagen Puregene isolation reagents (Germantown, MD) using a modification of the Puregene Handbook protocol for cell DNA isolation. To each cell pellet, 1 mL of cell lysis solution and 6 µL of proteinase K was added. After overnight shaking, 6 µL RNAse A was added. After 2 hours, 300 µL protein precipitation solution was added. Following 5 minutes on ice, the mixture was centrifuged at 3800 x g at 4°C for 10 minutes. The supernatant was poured into 1.3 mL cold 100% isopropanol and inverted to precipitate DNA. The DNA was pelleted by centrifugation, then rinsed with 70% cold isopropanol and 100% cold isopropanol, re-pelleting after each step. The DNA was dried and stored at -80°C until analysis.

### Mass Spectral analysis for POB DNA Adducts

Levels of 7-POBG, *O*^6^-POBdG, and *O*^2^-POBdT were measured in enzyme hydrolysates of DNA (18-50 µg) by published LC-MS/MS methods and expressed relative to the 2’-deoxyguanosine concentrations in the samples^60,61^.

### Exome sequencing of rat tumors

DNA was isolated from tumors and the surrounding tissue by DNeasy blood and tissue kit. Whole-exome libraries were captured with Agilent QXT Seq-Cap rat capture. Whole-exome sequencing was performed on the Illumina NovaSeq 6000 sequencer (S4 flow cell) in paired-end 150-bp runs to achieve 57x coverage.

### Standard sequencing data analysis

The FASTQ files were pre-processed by FastQC and Trim Galore and mapped on the human genome reference using the BWA-MEM aligner. Duplicate reads were flagged by the Picard tool and processed for base quality score recalibration and indel realignment with the corresponding tools from the GATK suite. Two somatic variant callers (MuTect2 and Strelka2) were applied to detect single base substitutions (SBS) and indels in exposed cell clones, using the parental cells as baseline reference. Consensus mutation data were processed with established in-house pipelines (https://github.com/IARCbioinfo), for variant allele frequency filtering, annotation with ANNOVAR, variant filtering to remove dbSNP contents, segmental duplicates, repeats, and tandem repeat regions. VCF files were used to plot sample mutation spectra using the SigProfilerExtractor tool^21^ (Supplementary Figure 8). Mutational signatures were extracted by the non-negative matrix factorization module within SigProfilerExtractor. The number of signatures to be extracted was estimated using SigProfilerExtractor, based on the stability and mean sample cosine distance for different signature numbers. Different numbers of signatures were extracted from each sample set and the optimal number, avoiding the splitting of unique signatures, was selected empirically. SigProfilerExtractor was further used to decompose the de novo signatures into established signature sets such as COSMIC content. Similarity among the extracted experimental signatures and with COSMIC signatures was evaluated using cosine similarity (lsa package, version 0.73.3).

### Whole Genome Universal-Duplex sequencing (UDSeq) and data analysis

Modified single molecule library preparation protocol was used to perform the library and sequencing for genome wide detection of mutations. Briefly: (i) To avoid damaged induced by mechanical fragmentation and also to capture whole genomic regions, enzymatic (NEBNext^®^ dsDNA Fragmentase) fragmentation approach was used to fragment the dsDNA for 20 min with 100 ng DNA as input; (ii) following fragmentation and AMPure beads purification, 10 ng DNA was carried out as input for library preparation by using IDT’s xGen™ cfDNA & FFPE DNA Library Prep Kit; (iii) after quantification of adapter ligated DNA molecules with qPCR, 0.2 fmol diluted DNA libraries were taken forward for unique dual indexing and Illumina 150PE sequencing. Sequencing reads were mapped on the human genome reference using the BWA-MEM aligner, as described above. An in-house developed error corrected mutation calling method DupCaller (https://github.com/AlexandrovLab/DupCaller) was used for somatic mutation calling using unexposed samples as matched normal. Mutational profile and signature analysis were performed with SigProfiler bioinformatic tools developed within the Alexandrov lab^31,62^ at UC San Diego. Mutation spectra of all analyzed samples are shown in Supplementary Figure 8.

### TwinStrand Duplex sequencing and data analysis

DNA extracted from NOK cells was processed for library preparation and sequencing according to the instructions provided by the manufacturer, using the TwinStrand Enzymatic Fragmentation Module and Duplex Sequencing Mutagenesis Kit (Human-50) (TwinStrand Biosciences, Seattle, WA, USA). Briefly, 750ng of DNA were enzymatically fragmented, followed by end repair, A-tailing and Duplex Sequencing Adapter ligation. After a PCR amplification step, the target DNA regions were captured by tandem hybridization steps using the provided biotinylated probes. Libraries were sequenced on the Illumina NovaSeq 6000 platform in PE150 configuration, aiming for ∼225 million reads per sample to achieve a Mean On-Target Duplex Depth of ∼25,000.

Demultiplexed FASTQ sequencing files were analyzed using the TwinStrand Biosciences DuplexSeq Mutagenesis App™ (version 4.2.0) on the DNAnexus platform. The methods included in the App have been previously described^58^. Raw sequencing reads were aligned to the human reference genome (hg38) using BWA, and read pairs were identified and grouped based on unique molecular identifiers. For downstream analysis, low quality bases were masked, followed by the generation of duplex consensus reads. To reduce biases or artifacts during variant calling, balanced overlap hard clipping and 5’ end trimming were applied. The Kraken taxonomic classifier was used to keep only duplex consensus reads that aligned to the human genome. Variant calling was performed using VarDictJava, and mutations present in more than one molecule of the sample were only counted once (as they were regarded results of clonal expansion). The generated VCF files were used to plot sample mutation spectra (Supplementary Figure 8) and to extract mutational signatures using the SigProfilerExtractor tool^21^.

### Mutational signature attribution to cancer genome data

*PCAWG and other published data*. Tumor data from PCAWG7 was extracted from Synapse^63^ and used for the signature attribution analysis. This dataset includes 37 different cancer sites and 2780 tumors (see Suppl Table 6). The transcription strand-biased (192-channel) matrix of the PCAWG data was extracted and used for attributions by 192-channel signature versions (see below). The raw BEAS-2B cell line data were obtained from Mingard et al^39^ and were reanalyzed to extract the TSB signature content as described above.

#### Screening for experimental mutational signatures

A set of 54 COSMIC signatures, compiled from from versions 3.3 and 3.2, was assembled by selecting all strand-biased signatures and removing signatures flagged as artifacts. The manually curated signatures were obtained from the official COSMIC website and were formatted into the 192-channel versions (see matrices in Supplementary Table 12 and spectra plots in Supplementary Figure 9). The PCAWG data described above were screened for the 54 COSMIC signatures alongside the 192-channel whole genome sequencing-derived SBS_POB signature obtained from A549 cells (see matrix in Supplementary Table 13), using Mutational Signature Attribution (MSA)^44^. Attribution levels for the SBS_POB signature (Supplementary Figure 5A), alongside the 54 COSMIC signatures in the PCAWG data are shown in Supplementary Table 14. The same process was repeated for the modelled, negative control-like signatures SBS_POB_flip (SBS_POB mutations flipped with the strand bias conserved; see Supplementary Figure 5B for signature plot) and SBS_POB_revTSB (SBS_POB strand bias reversed; see Supplementary Figure 5C for signature plot) (Supplementary Tables 15 and 16). MSA was conducted with the “removal” NNLS strategy, and the removal threshold was automatically optimized through genome modelling and customized bootstrapping, which also allows to calculate confidence intervals around attribution values. The analysis was prioritized for the experimental signature(s), and specificity was chosen as the target metric. In order to include the strand bias in the analysis, MSA was always used with 192 channels signatures and genomes. The interferences between signatures during the attribution process were assessed using Kendall correlation tests. A small number of SBS_POB positive samples was removed due to attribution of the negative control signatures SBS_POB_flip and/or SBS_POB_revTSB in the same samples.

#### Synthetic genome analysis

Based on the attribution results for the 54 COSMIC signatures (see above), 2780 synthetic tumors stochastically mimicking the COSMIC signature content in PCAWG data, with and without the experimentally observed SBS_POB signature were generated by using the simulation tool in MSA^44^ (https://gitlab.com/s.senkin/MSA/-/blob/master/bin/simulate_data.py). In each synthetic cohort we spiked ∼50% of samples with a low presence of SBS_POB (mean 5.7%, median 3.3%, range 2.3-24.3%), representing the ground truth scenario. MSA performed on the synthetic genomes allowed determining the false (FPR) and true positive rates (TPR) of signature attributions. Training of the MSA NNLS algorithm on this set established the FPRs, TPRs, and false discovery rates (FDR) of the attributions, as described previously^35^. The FDRs were calculated per cancer type and for the whole PCAWG 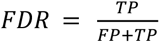

## Acknowledgments

This work was supported in part by the Cancer Research UK Grand Challenge 2016 Award “Mutographs of Cancer” C98/A24032 (L.B.A., J.Z.), National Cancer Institute grants R01CA184987 (L.A.P.), P30CA077598 (N.A.T. and L.A.P.) and NCI-CA220376 (S.B.), and research funds from the Masonic Cancer Center, University of Minnesota (N.A.T. and L.A.P.). This work was also supported by the US National Institute of Health grants R01ES032547-01, R01CA269919-01, and 1U01CA290479-01 to LBA as well as by LBA’s Packard Fellowship for Science and Engineering. The Hupki MEF exposure system had been established with the support from INCa-INSERM Plan Cancer 2015 (J.Z.) and NIH/NIEHS (1R03ES025023-01A1). The Analytical Biochemistry and the Comparative Pathology Shared Resources in the Masonic Cancer Center were supported in part by the National Cancer Institute grant P30CA077598. We would like to thank Marie-Pierre Cros for expert assistance with the chronic exposure and sequencing studies in cell lines, and TwinStrand Biosciences for expert assistance and guidance with the DuplexSeq^TM^ Mutagenesis Assay and data processing. We also thank Donna Seabloom, Monica Flavin, Beverly Wuertz, Atlanta Bidinger, Anna Haynes, Elizabeth Bonilla, Tia Eskridge, Ebisie Deressa, Makenzie Cavil, Benjamin Ransom, Adam Zarth and Erik Carlson for their excellent assistance with the rat studies.

## Competing interests

LBA is a co-founder, CSO, scientific advisory member, and consultant for io9, has equity and receives income. The terms of this arrangement have been reviewed and approved by the University of California, San Diego in accordance with its conflict of interest policies. LBA is a compensated member of the scientific advisory board of Inocras. LBA’s spouse is an employee of Biotheranostics, Inc. LBA declares U.S. provisional applications with serial numbers: 63/289,601; 63/269,033; 63/366,392; 63/412,835 as well as international patent application PCT/US2023/010679. LBA is also an inventor of a US Patent 10,776,718 for source identification by non-negative matrix factorization. All other authors declare that they have no competing interests.

## Contributions

The study was conceived and designed and supervised by M.K., J.Z., L.A.P., N.A.T and S.S.H., and it was supervised by M.K., J.Z., L.A.P. and N.A.T. Data generation was performed by B.C., T.M., M.K., C.S., K.R.V., J.G., W.E.S., M.K.O., F.A.T., O.A., I.C., M.G.O, S.D. and S.B. Analysis of data was carried out by S.K., F.V., C.R., V.C., N.A.T., F.C.J., S.N., Y.C., L.B.A. and S.S. The manuscript was written by M.K., J.Z., L.A.P., and N.A.T., with contributions from all authors.

## Disclaimer

Where authors are identified as personnel of the International Agency for Research on Cancer/World Health Organization, the authors alone are responsible for the views expressed in this article and they do not necessarily represent the decisions, policy or views of the International Agency for Research on Cancer /World Health Organization.

## REFERENCES

1. Freedman, N. D. & Thun, M. J. in World Cancer Report: Cancer Research for Cancer Prevention (eds C. P. Wild, E. Weiderpass, & B. W. Stewart) 50-60 (International Agency for Research on Cancer, 2020).

2 National Cancer Institute & Centers for Disease Control and Prevention. Smokeless Tobacco and Public Health: A Global Perspective.. (Bethesda, MD, 2014).

3. Rodgman, A. & Perfetti, T. A. The chemical components of tobacco and tobacco smoke. 2nd Edition edn, (CRC Press, 2013).

4 Li, Y. & Hecht, S. S. Carcinogenic components of tobacco and tobacco smoke: A 2022 update. Food Chem Toxicol 165, 113179, doi:10.1016/j.fct.2022.113179 (2022).

5 Hecht, S. S. Biochemistry, biology, and carcinogenicity of tobacco-specific *N*-nitrosamines. Chem. Res. Toxicol.. 11, 559–603 (1998).

6 Castonguay, A. et al. Effects of chronic ethanol consumption on the metabolism and carcinogenicity of N’-nitrosonornicotine in F344 rats. Cancer Research 44, 2285–2290 (1984).

7 Balbo, S. et al. (*S*)-*N*’-Nitrosonornicotine, a constituent of smokeless tobacco, is a powerful oral cavity carcinogen in rats. Carcinogenesis 34, 2178–2183 (2013).

8 Yuan, J. M. et al. Urinary levels of the tobacco-specific carcinogen *N*’-nitrosonornicotine and its glucuronide are strongly associated with esophageal cancer risk in smokers. Carcinogenesis 32, 1366–1371 (2011).

9 Stepanov, I. et al. Tobacco-specific *N*-nitrosamine exposures and cancer risk in the Shanghai cohort study: Remarkable coherence with rat tumor sites. Int. J. Cancer 134, 2278–2283 (2014).

10 Hecht, S. S., Chen, C. B., Ohmori, T. & Hoffmann, D. Comparative carcinogenicity in F344 rats of the tobacco specific nitrosamines, *N*’-nitrosonornicotine and 4-(*N*-methyl-*N*-nitrosamino)-1-(3-pyridyl)-1-butanone. Cancer Research 40, 298–302 (1980).

11 Rivenson, A., Hoffmann, D., Prokopczyk, B., Amin, S. & Hecht, S. S. Induction of lung and exocrine pancreas tumors in F344 rats by tobacco-specific and areca-derived *N*-nitrosamines. Cancer Res. 48, 6912–6917 (1988).

12 Hecht, S. S. et al. Rapid single dose model for lung tumor induction in A/J mice by 4-(methylnitrosamino)-1-(3-pyridyl)-1-butanone and the effect of diet. Carcinogenesis 10, 1901–1904 (1989).

13 Yuan, J. M. et al. Urinary levels of cigarette smoke constituent metabolites are prospectively associated with lung cancer development in smokers. Cancer Res 71, 6749–6757, doi:10.1158/0008-5472.CAN-11-0209 (2011).

14. International Agency for Research on Cancer. Personal Habits and Indoor Combustions. Vol. 100E (IARC, 2012).

15 Church, T. R. et al. A prospectively measured serum biomarker for a tobacco-specific carcinogen and lung cancer in smokers. Cancer Epidemiol Biomarkers Prev 18, 260–266, doi:10.1158/1055-9965.EPI-08-0718 (2009).

16 Yuan, J. M., Butler, L. M., Stepanov, I. & Hecht, S. S. Urinary tobacco smoke-constituent biomarkers for assessing risk of lung cancer. Cancer Res 74, 401–411, doi:10.1158/0008-5472.CAN-13-3178 (2014).

17 Murphy, S. E. et al. Association of urinary biomarkers of tobacco exposure with lung cancer risk in African American and White cigarette smokers in the Southern Community Cohort Study. Cancer Epidemiol Biomarkers Prev, doi:10.1158/1055-9965.EPI-23-1362 (2024).

18 Alexandrov, L. B. et al. Signatures of mutational processes in human cancer. Nature 500, 415–421, doi:10.1038/nature12477 (2013).

19. Wellcome Sanger Institute. <https://cancer.sanger.ac.uk/cosmic/signatures> (

20 Degasperi, A. et al. Substitution mutational signatures in whole-genome-sequenced cancers in the UK population. Science 376, doi:10.1126/science.abl9283 (2022).

21 Alexandrov, L. B. et al. The repertoire of mutational signatures in human cancer. Nature 578, 94–101, doi:10.1038/s41586-020-1943-3 (2020).

22 Steele, C. D. et al. Signatures of copy number alterations in human cancer. Nature 606, 984–991, doi:10.1038/s41586-022-04738-6 (2022).

23 Everall, A. et al. Comprehensive repertoire of the chromosomal alteration and mutational signatures across 16 cancer types from 10,983 cancer patients. medRxiv, 2023.2006.2007.23290970, doi:10.1101/2023.06.07.23290970 (2023).

24 Hollstein, M., Alexandrov, L. B., Wild, C. P., Ardin, M. & Zavadil, J. Base changes in tumour DNA have the power to reveal the causes and evolution of cancer. Oncogene 36, 158–167, doi:10.1038/onc.2016.192 (2017).

25 Volkova, N. V. et al. Mutational signatures are jointly shaped by DNA damage and repair. Nat Commun 11, 2169, doi:10.1038/s41467-020-15912-7 (2020).

26 Ivanov, D., Hwang, T., Sitko, L. K., Lee, S. & Gartner, A. Experimental systems for the analysis of mutational signatures: no ’one-size-fits-all’ solution. Biochem Soc Trans 51, 1307–1317, doi:10.1042/BST20221482 (2023).

27 Melki, P. N., Korenjak, M. & Zavadil, J. Experimental investigations of carcinogen-induced mutation spectra: Innovation, challenges and future directions. Mutat Res 853, 503195, doi:10.1016/j.mrgentox.2020.503195 (2020).

28 Alexandrov, L. B. et al. Mutational signatures associated with tobacco smoking in human cancer. Science 354, 618–622, doi:10.1126/science.aag0299 (2016).

29. India Project Team of the International Cancer Genome, C. Mutational landscape of gingivo-buccal oral squamous cell carcinoma reveals new recurrently-mutated genes and molecular subgroups. Nat Commun 4, 2873, doi:10.1038/ncomms3873 (2013).

30 Lawson, A. R. J. et al. Extensive heterogeneity in somatic mutation and selection in the human bladder. Science 370, 75–82, doi:10.1126/science.aba8347 (2020).

31 Islam, S. M. A. et al. Uncovering novel mutational signatures by de novo extraction with SigProfilerExtractor. Cell Genom 2, None, doi:10.1016/j.xgen.2022.100179 (2022).

32 Kucab, J. E. et al. A compendium of mutational signatures of environmental agents. Cell 177, 821–836 e816, doi:10.1016/j.cell.2019.03.001 (2019).

33 Nik-Zainal, S. et al. The genome as a record of environmental exposure. Mutagenesis 30, 763–770, doi:10.1093/mutage/gev073 (2015).

34 Olivier, M. et al. Modelling mutational landscapes of human cancers *in vitro*. Sci. Rep. 4, 4482, doi:10.1038/srep04482 (2014).

35 Zhivagui, M. et al. Experimental and pan-cancer genome analyses reveal widespread contribution of acrylamide exposure to carcinogenesis in humans. Genome Res. 29, 521–531, doi:10.1101/gr.242453.118 (2019).

36 Yoshida, K. et al. Tobacco smoking and somatic mutations in human bronchial epithelium. Nature 578, 266–272, doi:10.1038/s41586-020-1961-1 (2020).

37 Torrens, L. et al. The Complexity of Tobacco Smoke-Induced Mutagenesis in Head and Neck Cancer. medRxiv, doi:10.1101/2024.04.15.24305006 (2024).

38 Peterson, L. A. Context matters: Contribution of specific DNA adducts to the genotoxic properties of the tobacco-specific nitrosamine NNK. Chem Res Toxicol 30, 420–433, doi:10.1021/acs.chemrestox.6b00386 (2017).

39 Mingard, C. et al. Dissection of Cancer Mutational Signatures with Individual Components of Cigarette Smoking. Chem Res Toxicol 36, 714–723, doi:10.1021/acs.chemrestox.3c00021 (2023).

40 Holzl-Armstrong, L. et al. Mutagenicity of acrylamide and glycidamide in human TP53 knock-in (Hupki) mouse embryo fibroblasts. Arch Toxicol 94, 4173–4196, doi:10.1007/s00204-020-02878-0 (2020).

41 Holzl-Armstrong, L. et al. Mutagenicity of 2-hydroxyamino-1-methyl-6-phenylimidazo[4,5-b]pyridine (N-OH-PhIP) in human TP53 knock-in (Hupki) mouse embryo fibroblasts. Food Chem Toxicol 147, 111855, doi:10.1016/j.fct.2020.111855 (2021).

42 Riva, L. et al. The mutational signature profile of known and suspected human carcinogens in mice. Nat Genet 52, 1189–1197, doi:10.1038/s41588-020-0692-4 (2020).

43 Behjati, S. et al. Genome sequencing of normal cells reveals developmental lineages and mutational processes. Nature 513, 422–425, doi:10.1038/nature13448 (2014).

44 Senkin, S. MSA: reproducible mutational signature attribution with confidence based on simulations. BMC Bioinformatics 22, 540, doi:10.1186/s12859-021-04450-8 (2021).

45 Weerasooriya, S., Jasti, V. P., Bose, A., Spratt, T. E. & Basu, A. K. Roles of translesion synthesis DNA polymerases in the potent mutagenicity of tobacco-specific nitrosamine-derived O2-alkylthymidines in human cells. DNA Repair (Amst) 35, 63–70 (2015).

46 Jasti, V. P., Spratt, T. E. & Basu, A. K. Tobacco-specific nitrosamine-derived *O*^2^-alkylthymidines are potent mutagenic lesions in SOS-induced Escherichia coli. Chem. Res. Toxicol. 24, 1833–1835 (2011).

47 Hecht, S. S. et al. Identification of *O*^2^-substituted pyrimidine adducts formed in reactions of 4-(acetoxymethylnitrosamino)-1-(3-pyridyl)-1-butanone and 4-(acetoxymethylnitrosamino)-1-(3-pyridyl)-1-butanol with DNA. Chem. Res. Toxicol. 17, 588–597 (2004).

48 Pluym, N. et al. Assessment of the Exposure to NNN in the Plasma of Smokeless Tobacco Users. Chem Res Toxicol 35, 663–669, doi:10.1021/acs.chemrestox.1c00431 (2022).

49 Morton, L. M. et al. Cigarette smoking and risk of non-Hodgkin lymphoma: a pooled analysis from the International Lymphoma Epidemiology Consortium (interlymph). Cancer Epidemiol Biomarkers Prev 14, 925–933, doi:10.1158/1055-9965.EPI-04-0693 (2005).

50 Taborelli, M. et al. The dose-response relationship between tobacco smoking and the risk of lymphomas: a case-control study. BMC Cancer 17, 421, doi:10.1186/s12885-017-3414-2 (2017).

51 Zarth, A. T., Upadhyaya, P., Yang, J. & Hecht, S. S. DNA Adduct formation from metabolic 5′-hydroxylation of the tobacco-specific carcinogen N′-nitrosonornicotine in human enzyme systems and in rats. Chem. Res. Toxicol. 29, 380–389, doi:10.1021/acs.chemrestox.5b00520 (2016).

52 Li, Y. & Hecht, S. S. Identification of an N’-Nitrosonornicotine-Specific Deoxyadenosine Adduct in Rat Liver and Lung DNA. Chem Res Toxicol 34, 992–1003, doi:10.1021/acs.chemrestox.1c00013 (2021).

53 Li, Y., Carlson, E. S., Zarth, A. T., Upadhyaya, P. & Hecht, S. S. Investigation of 2’-Deoxyadenosine-Derived Adducts Specifically Formed in Rat Liver and Lung DNA by N’-Nitrosonornicotine Metabolism. Chem Res Toxicol 34, 1004–1015, doi:10.1021/acs.chemrestox.1c00012 (2021).

54 Foiles, P. G. et al. Mass spectrometric analysis of tobacco-specific nitrosamine DNA adducts in smokers and non-smokers. Chem. Res. Toxicol.. 4, 364–368 (1991).

55 Khariwala, S. S. et al. High Level of Tobacco Carcinogen-Derived DNA Damage in Oral Cells Is an Independent Predictor of Oral/Head and Neck Cancer Risk in Smokers. Cancer Prev Res (Phila) 10, 507–513, doi:10.1158/1940-6207.CAPR-17-0140 (2017).

56 Holzle, D., Schlobe, D., Tricker, A. R. & Richter, E. Mass spectrometric analysis of 4-hydroxy-1-(3-pyridyl)-1-butanone-releasing DNA adducts in human lung. Toxicology 232, 277–285 (2007).

57 Humans, I. W. G. o. t. E. o. C. R. t. Smokeless tobacco and some tobacco-specific N-nitrosamines. IARC Monogr Eval Carcinog Risks Hum 89, 1–592 (2007).

58 Valentine, C. C., 3rd et al. Direct quantification of in vivo mutagenesis and carcinogenesis using duplex sequencing. Proc Natl Acad Sci U S A 117, 33414–33425, doi:10.1073/pnas.2013724117 (2020).

59 LeBlanc, D. P. M. et al. Duplex sequencing identifies genomic features that determine susceptibility to benzo(a)pyrene-induced in vivo mutations. BMC Genomics 23, 542, doi:10.1186/s12864-022-08752-w (2022).

60 Lao, Y., Villalta, P. W., Sturla, S. J., Wang, M. & Hecht, S. S. Quantitation of pyridyloxobutyl DNA adducts of tobacco-specific nitrosamines in rat tissue DNA by high-performance liquid chromatography-electrospray ionization-tandem mass spectrometry. Chem. Res. Toxicol 19, 674–682 (2006).

61 Thomson, N. M. et al. Development of a quantitative liquid chromatography/electrospray mass spectrometric assay for a mutagenic tobacco-specific nitrosamine-derived DNA adduct, *O*^6^-[4-oxo-4-(3-pyridyl)butyl]-2’-deoxyguanosine. Chemical Research in Toxicology 17, 1600–1606 (2004).

62 Bergstrom, E. N. et al. SigProfilerMatrixGenerator: a tool for visualizing and exploring patterns of small mutational events. BMC Genomics 20, 685, doi:10.1186/s12864-019-6041-2 (2019).

63. ICGC Pan Cancer Analysis Mutational Signatures Working Group. PCAWG7-public-sup-info, <https://www.synapse.org/#!Synapse:syn11726601/wiki/513478> (2018).

