## Supplementary Figures for "Human cancer genomes harbor the mutational signature of tobacco-specific nitrosamines NNN and NNK"

Supplementary Figure 1

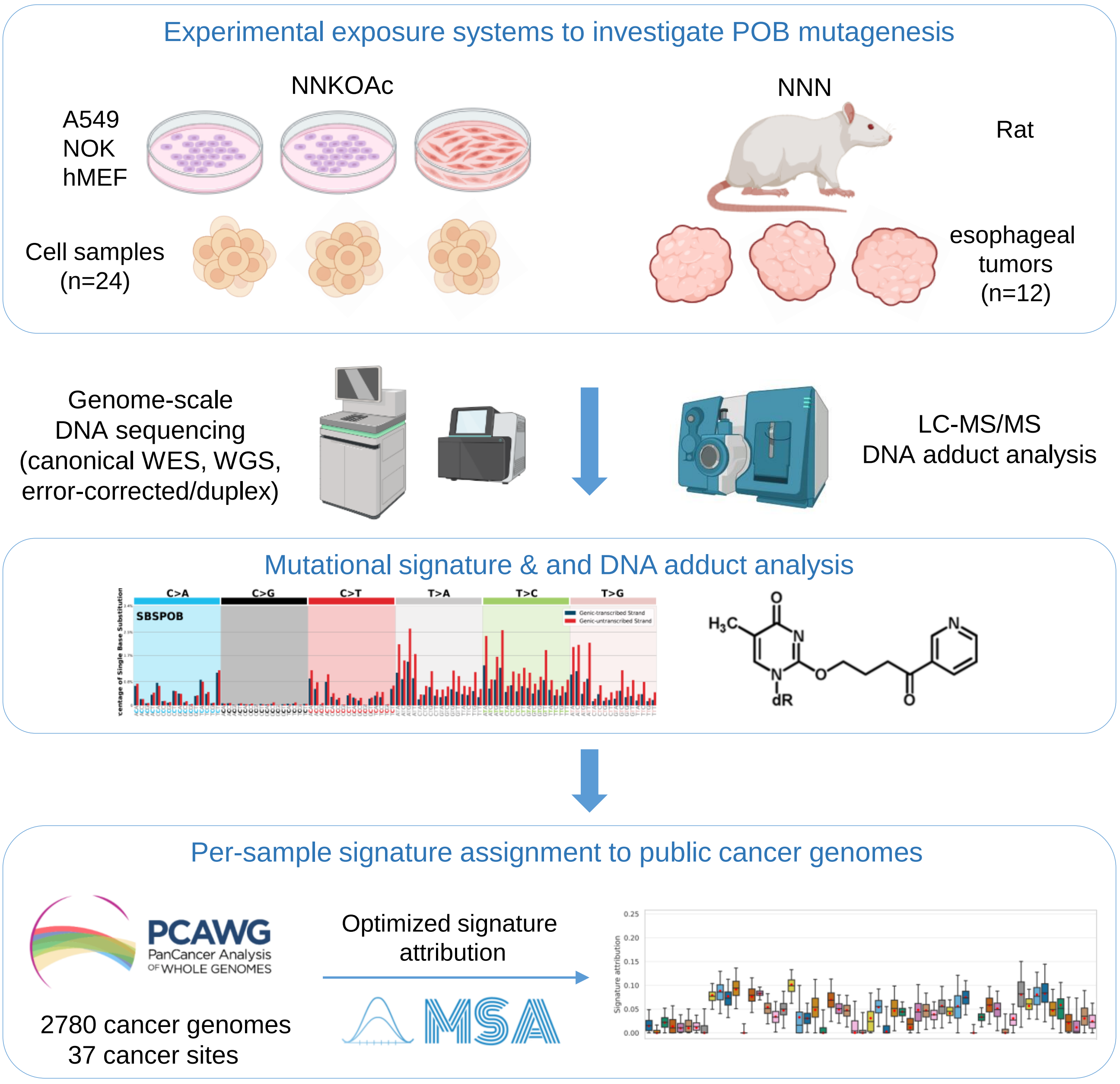

Supplementary Figure 2

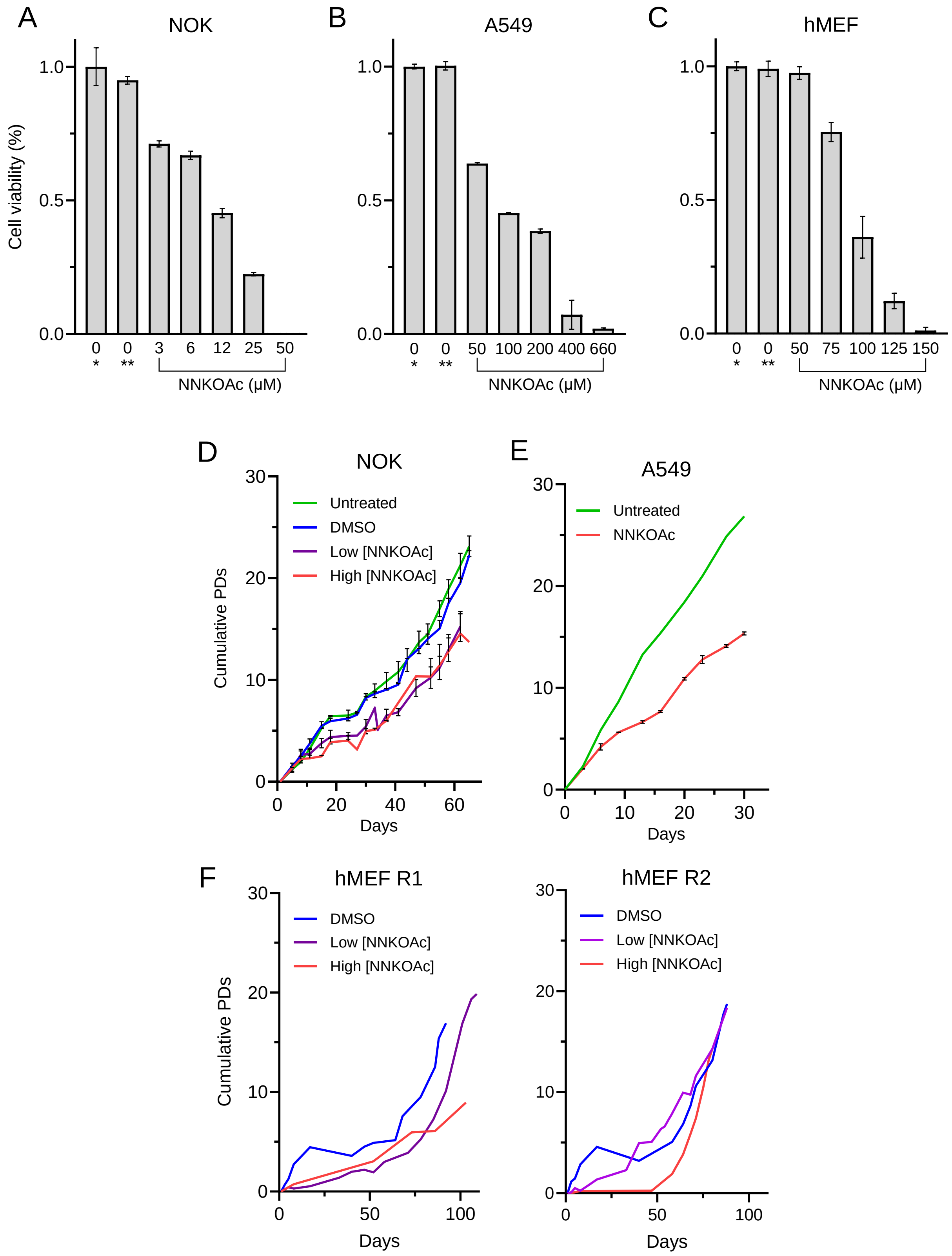

Supplementary Figure 3

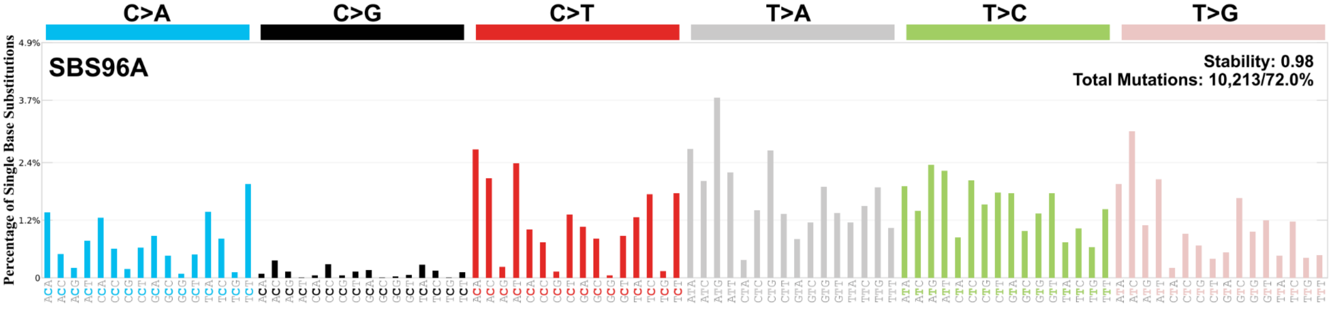

Supplementary Figure 4

A

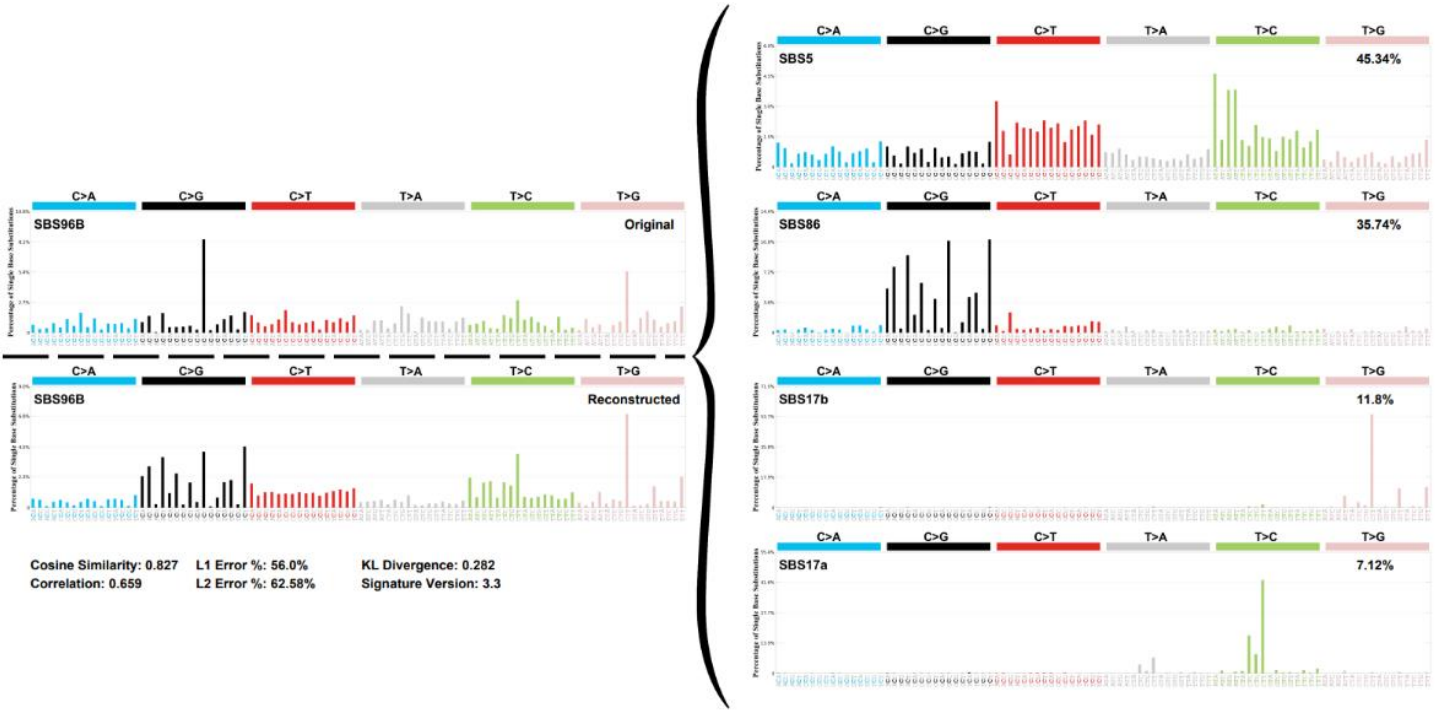


B

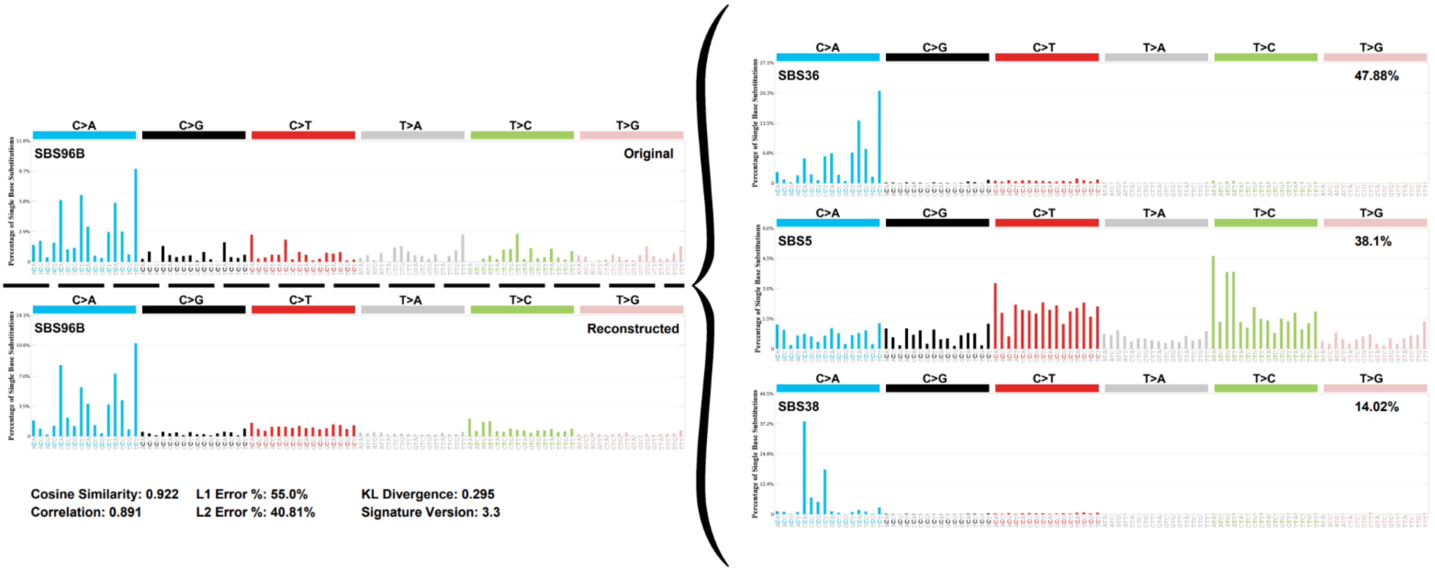

### Supplementary Figure 5

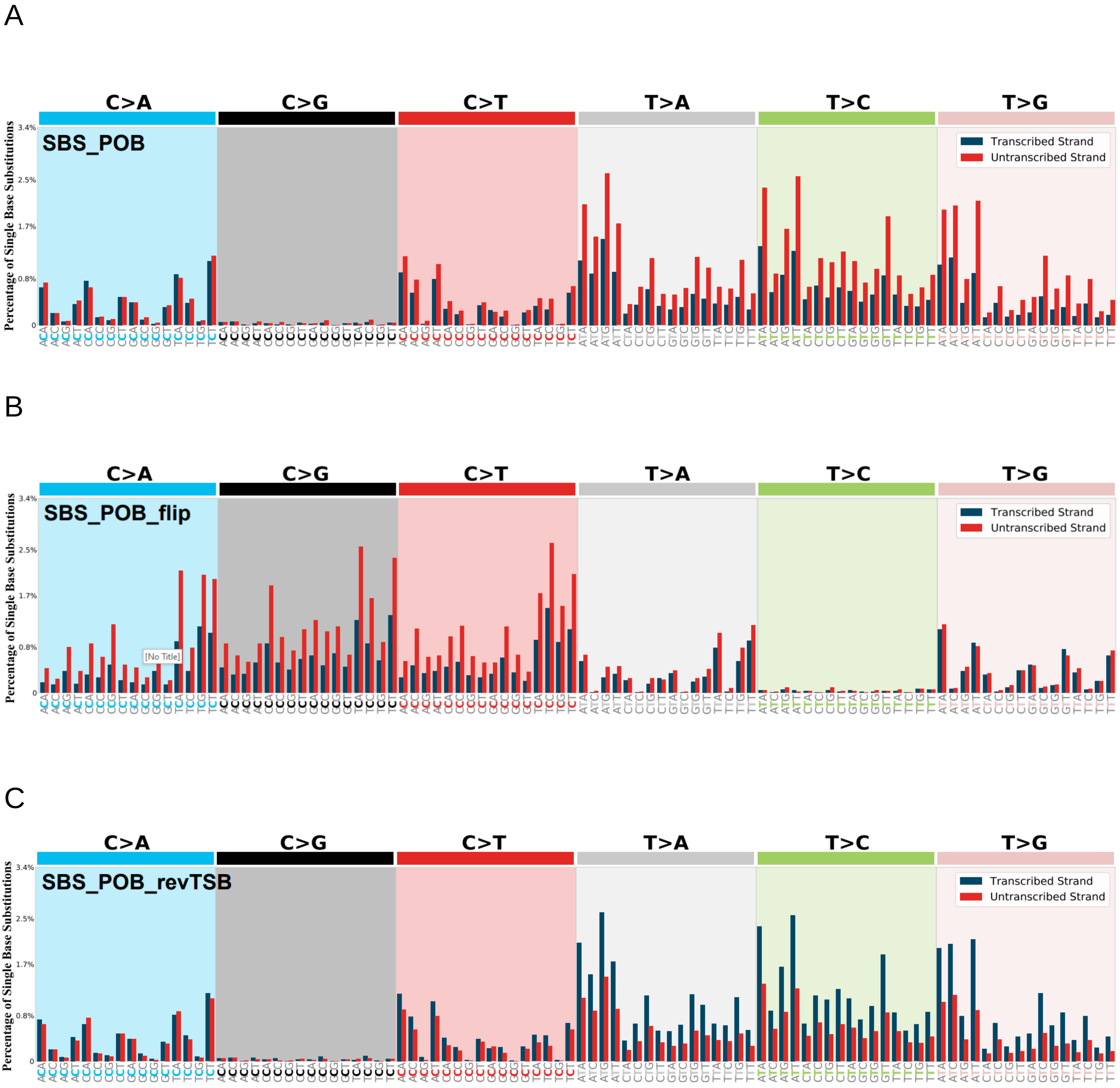

Supplementary Figure 6

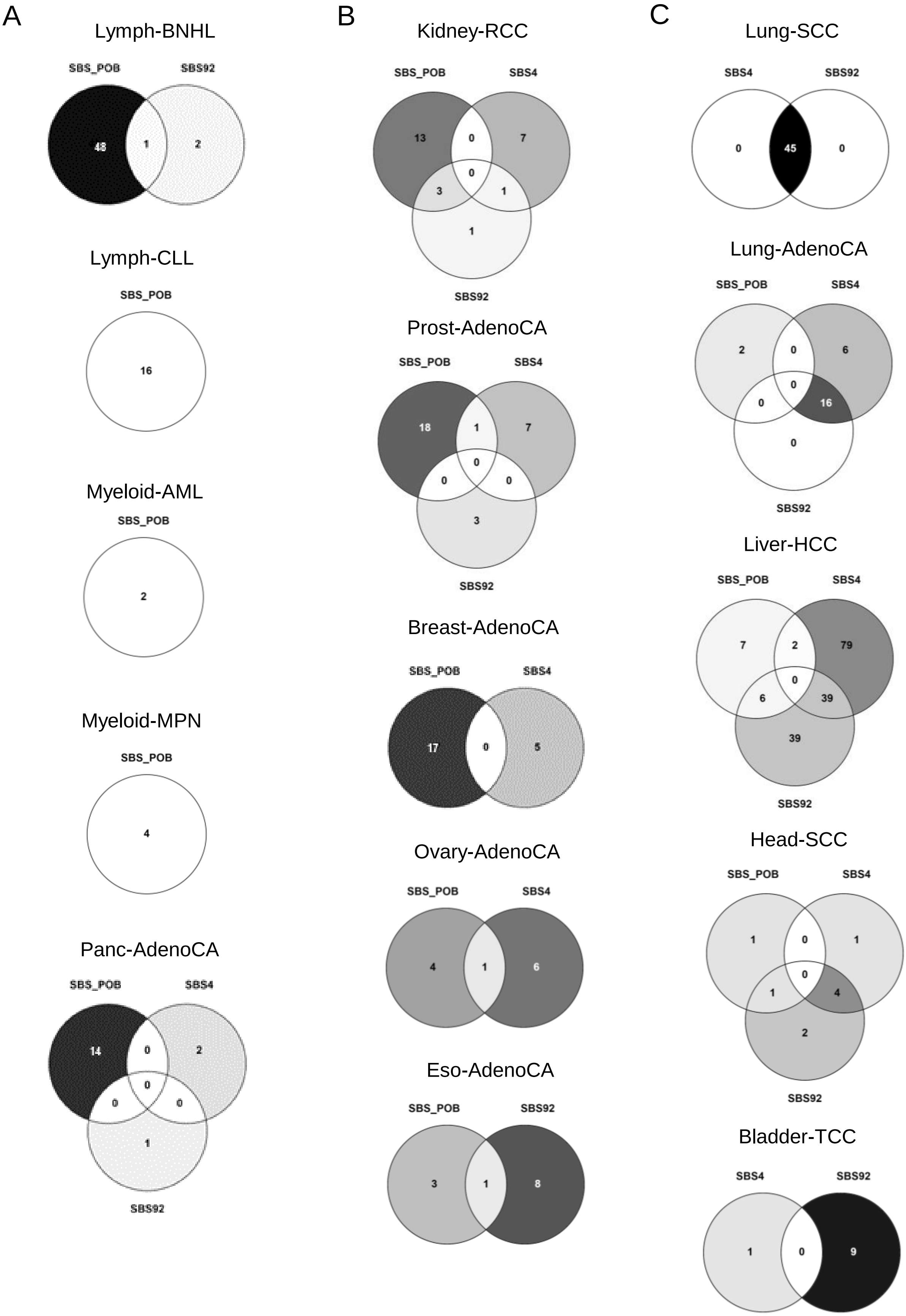

### Supplementary Figure 7

A

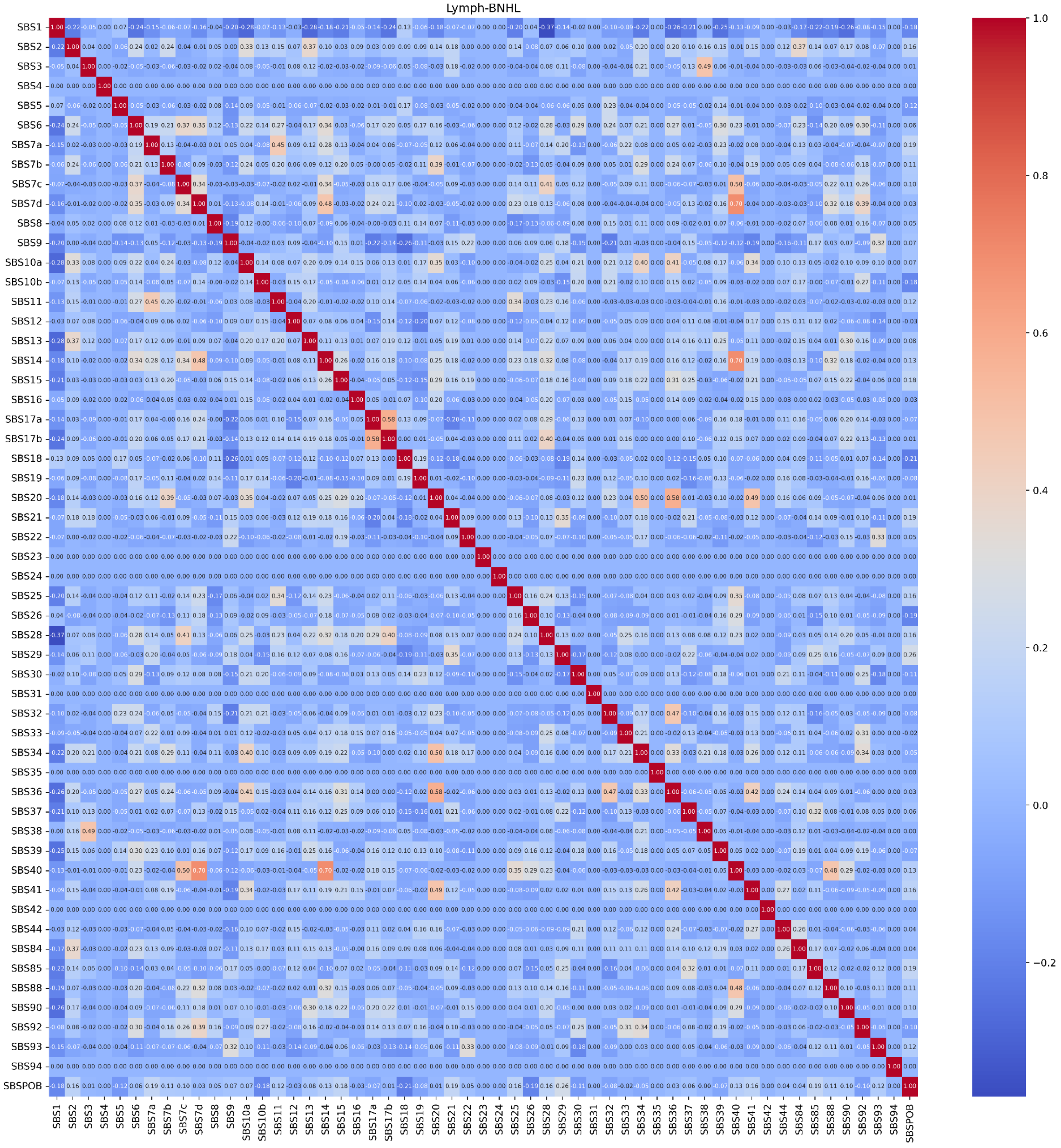

B

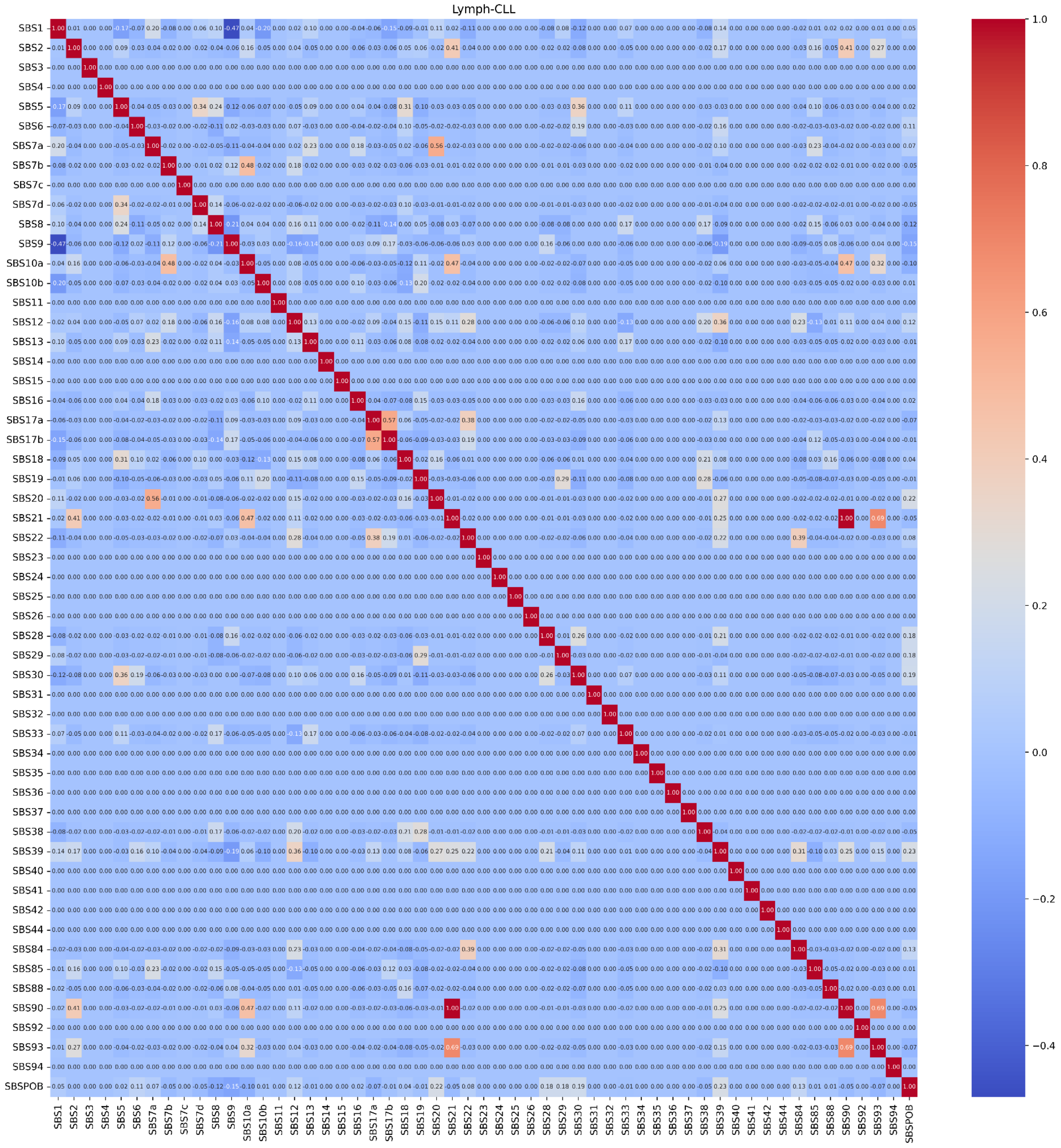
Lymph-CLL
SBS1 SBS2 SBS3 SBS4 SBS5 SBS6 SBS7a SBS7b SBS7c SBS7d SBS8 SBS9 SBS10a SBS10b SBS11 SBS12 SBS13 SBS14 SBS15 SBS16 SBS17a SBS17b SBS18 SBS19 SBS20 SBS21 SBS22 SBS23 SBS24 SBS25 SBS26 SBS28 SBS29 SBS30 SBS31 SBS32 SBS33 SBS34 SBS35 SBS36 SBS37 SBS38 SBS39 SBS40 SBS41 SBS42 SBS44 SBS84 SBS85 SBS88 SBS90 SBS92 SBS93 SBS94 SBSPOB
1.0
0.8
0.6
0.4
0.2
0.0
−0.2
−0.4

Supplementary Figure 8

A

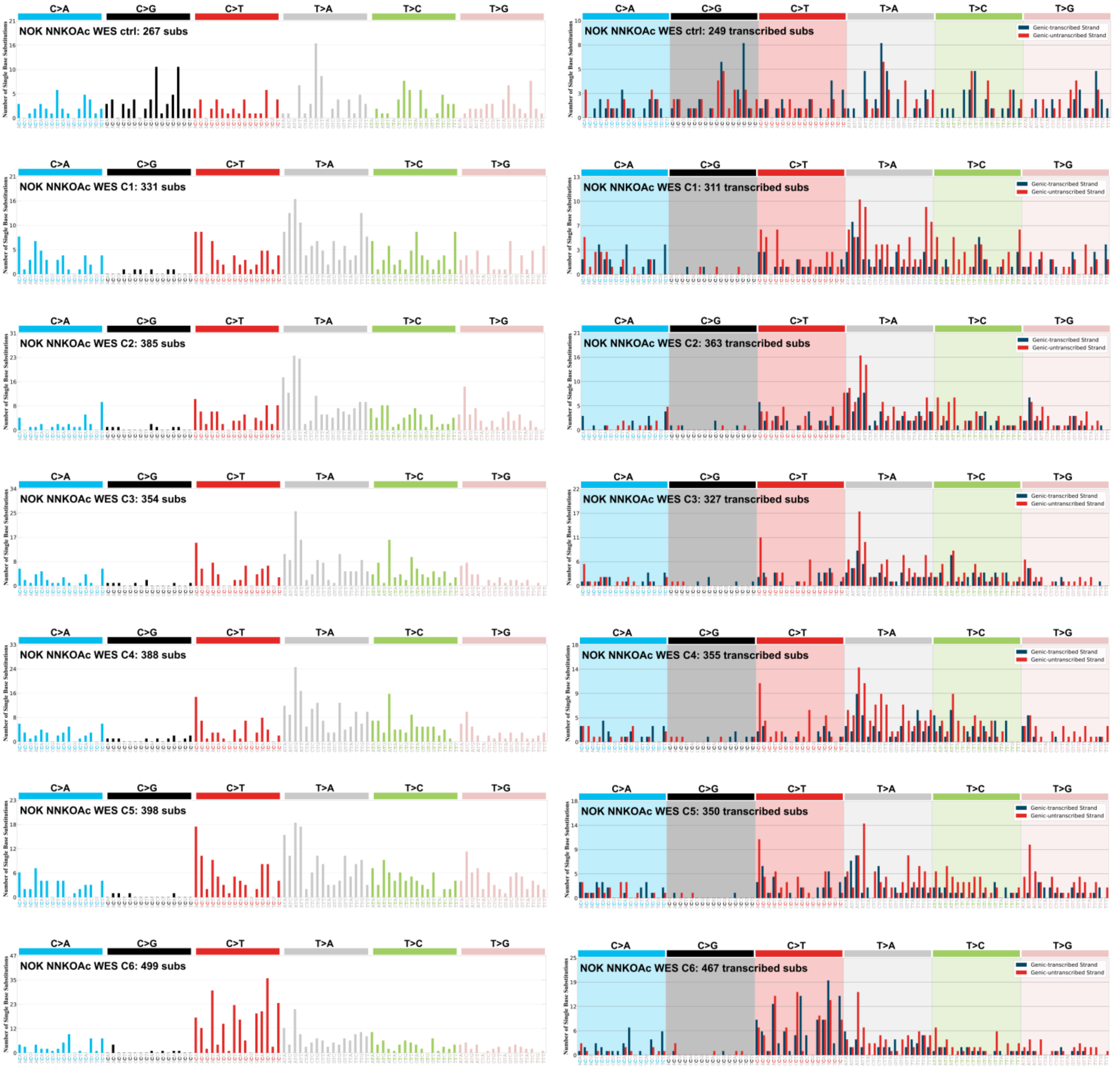

B

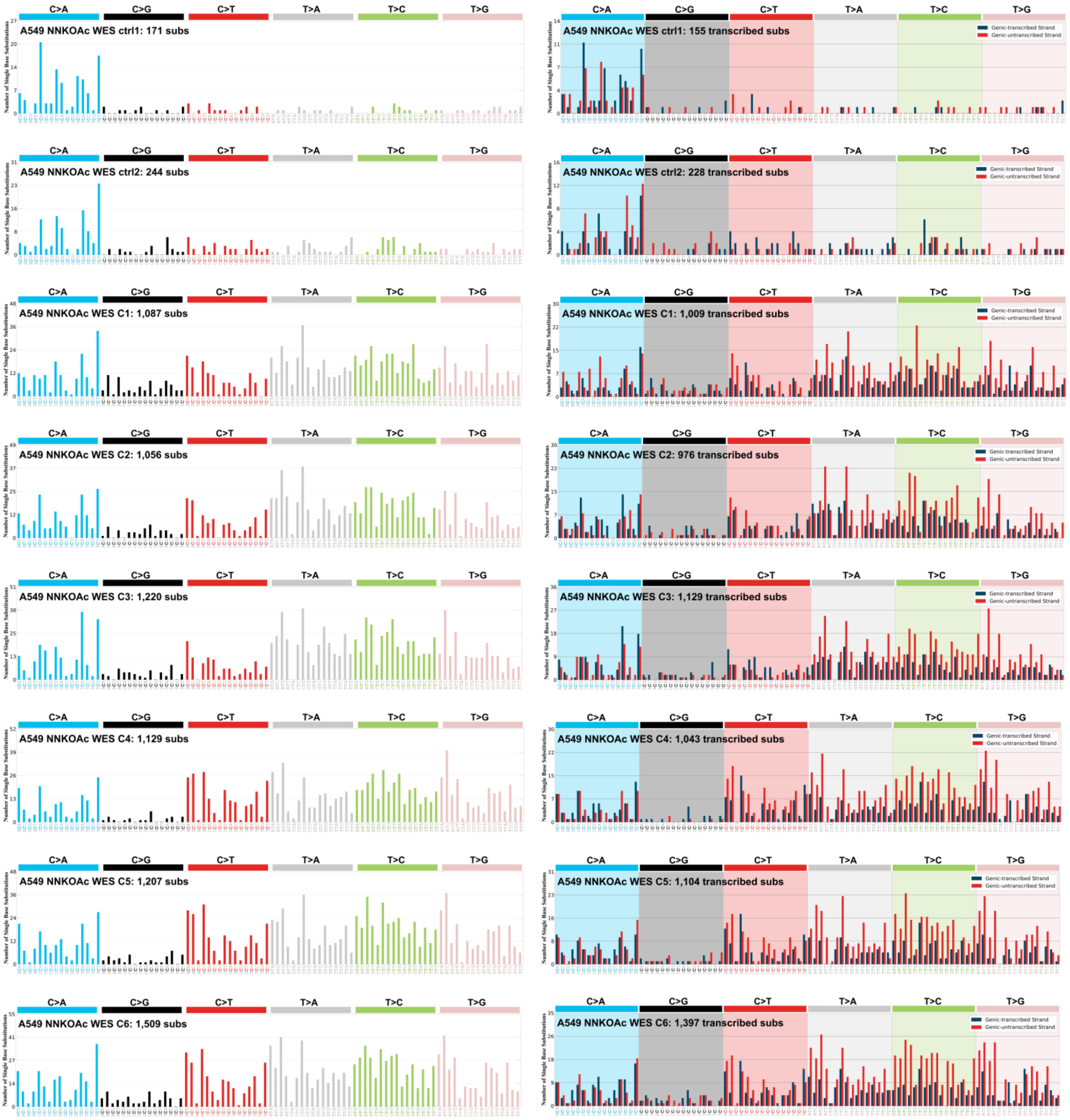
C>A
C>G
C>T
T>A
T>C
T>G
Number of Single Base Substitutions
A549 NNKOAc WES ctrl1: 171 subs
A549 NNKOAc WES ctrl1: 155 transcribed subs
A549 NNKOAc WES ctrl2: 244 subs
A549 NNKOAc WES ctrl2: 228 transcribed subs
A549 NNKOAc WES C1: 1,087 subs
A549 NNKOAc WES C1: 1,009 transcribed subs
A549 NNKOAc WES C2: 1,056 subs
A549 NNKOAc WES C2: 976 transcribed subs
A549 NNKOAc WES C3: 1,220 subs
A549 NNKOAc WES C3: 1,129 transcribed subs
A549 NNKOAc WES C4: 1,129 subs
A549 NNKOAc WES C4: 1,043 transcribed subs
A549 NNKOAc WES C5: 1,207 subs
A549 NNKOAc WES C5: 1,104 transcribed subs
A549 NNKOAc WES C6: 1,509 subs
A549 NNKOAc WES C6: 1,397 transcribed subs
Genic-transcribed Strand
Genic-untranscribed Strand

C

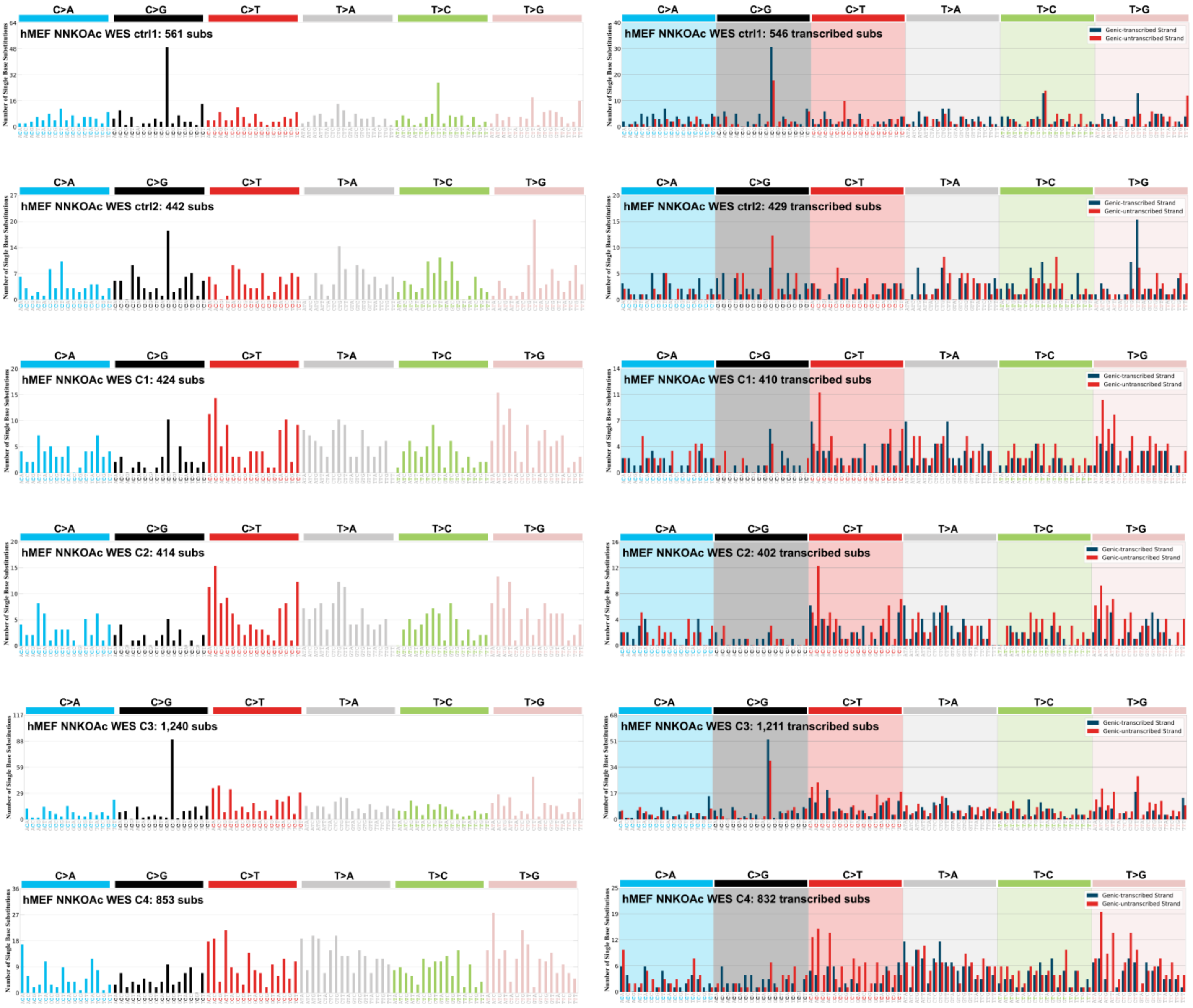

C>A
C>G
C>T
T>A
T>C
T>G
Number of Single Base Substitutions
hMEF NNKOAc WES ctrl1: 561 subs
hMEF NNKOAc WES ctrl1: 546 transcribed subs
hMEF NNKOAc WES ctrl2: 442 subs
hMEF NNKOAc WES ctrl2: 429 transcribed subs
hMEF NNKOAc WES C1: 424 subs
hMEF NNKOAc WES C1: 410 transcribed subs
hMEF NNKOAc WES C2: 414 subs
hMEF NNKOAc WES C2: 402 transcribed subs
hMEF NNKOAc WES C3: 1,240 subs
hMEF NNKOAc WES C3: 1,211 transcribed subs
hMEF NNKOAc WES C4: 853 subs
hMEF NNKOAc WES C4: 832 transcribed subs
Genic-transcribed Strand
Genic-untranscribed Strand

D

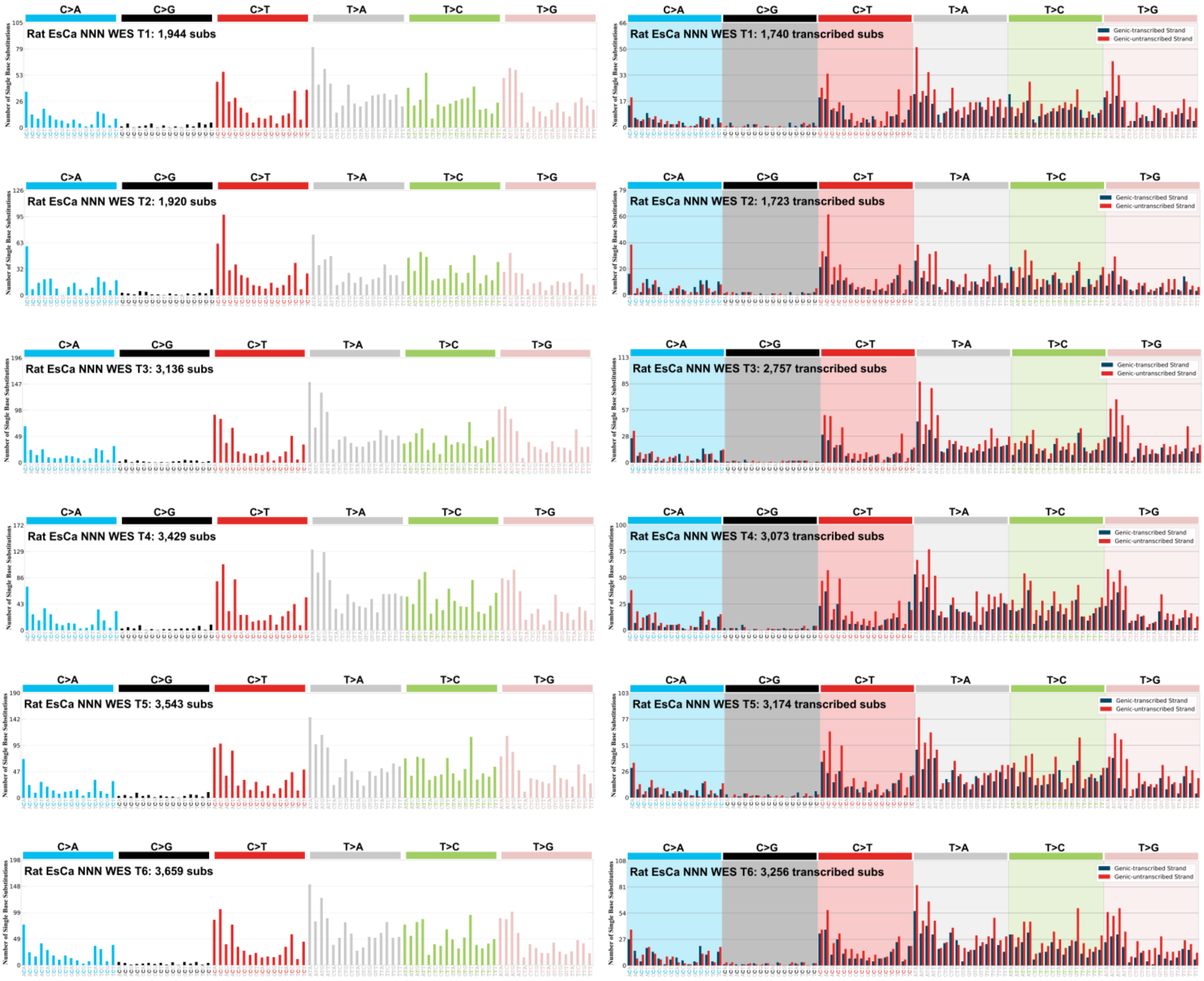

C>A
C>G
C>T
T>A
T>C
T>G
Rat EsCa NNN WES T1: 1,944 subs
Rat EsCa NNN WES T1: 1,740 transcribed subs
Rat EsCa NNN WES T2: 1,920 subs
Rat EsCa NNN WES T2: 1,723 transcribed subs
Rat EsCa NNN WES T3: 3,136 subs
Rat EsCa NNN WES T3: 2,757 transcribed subs
Rat EsCa NNN WES T4: 3,429 subs
Rat EsCa NNN WES T4: 3,073 transcribed subs
Rat EsCa NNN WES T5: 3,543 subs
Rat EsCa NNN WES T5: 3,174 transcribed subs
Rat EsCa NNN WES T6: 3,659 subs
Rat EsCa NNN WES T6: 3,256 transcribed subs
Genic-transcribed Strand
Genic-untranscribed Strand
Number of Single Base Substitutions

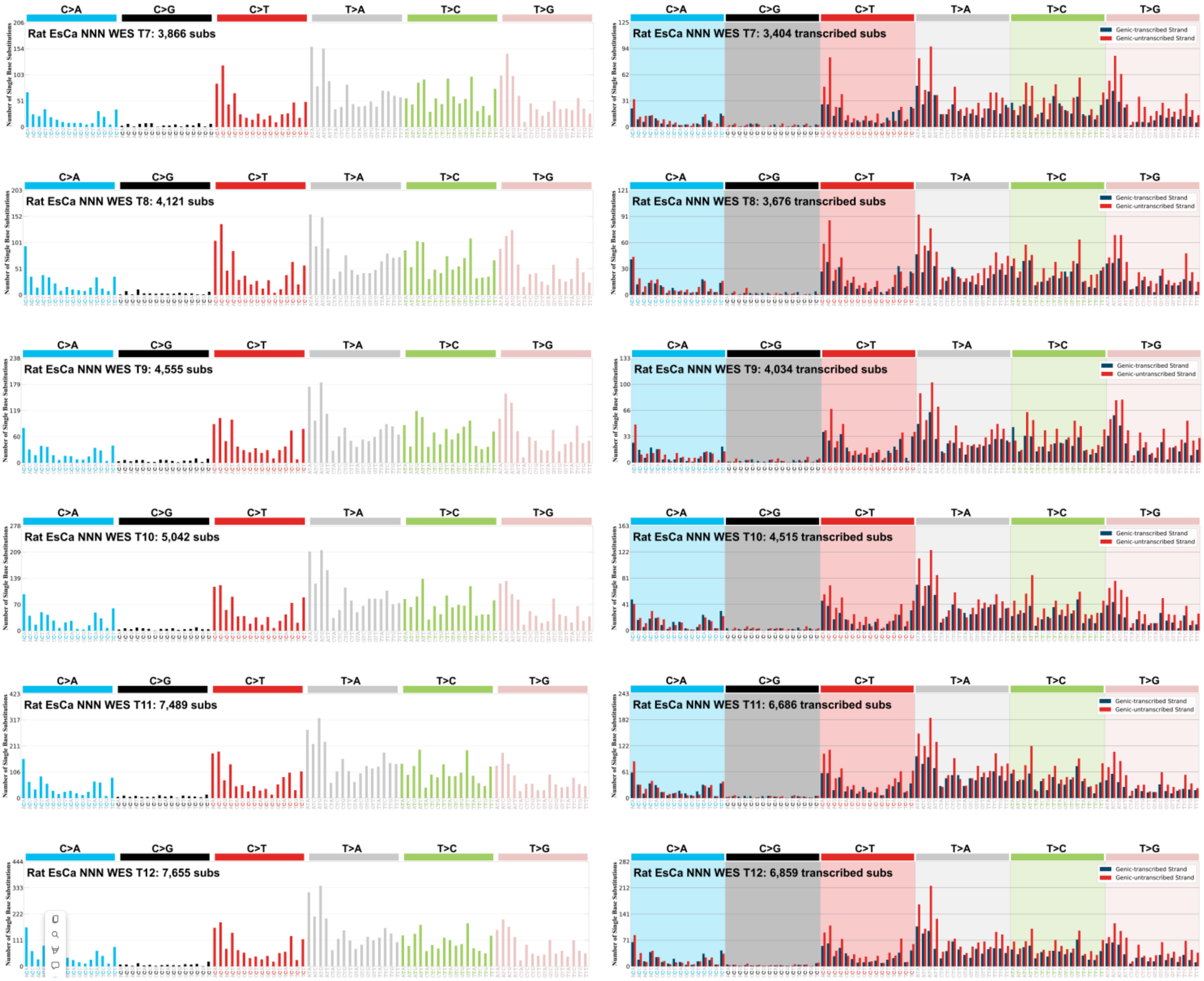
Rat EsCa NNN WES T7: 3,866 subs
Rat EsCa NNN WES T7: 3,404 transcribed subs
Rat EsCa NNN WES T8: 4,121 subs
Rat EsCa NNN WES T8: 3,676 transcribed subs
Rat EsCa NNN WES T9: 4,555 subs
Rat EsCa NNN WES T9: 4,034 transcribed subs
Rat EsCa NNN WES T10: 5,042 subs
Rat EsCa NNN WES T10: 4,515 transcribed subs
Rat EsCa NNN WES T11: 7,489 subs
Rat EsCa NNN WES T11: 6,686 transcribed subs
Rat EsCa NNN WES T12: 7,655 subs
Rat EsCa NNN WES T12: 6,859 transcribed subs
C>A
C>G
C>T
T>A
T>C
T>G
Genic-transcribed Strand
Genic-untranscribed Strand
Number of Single Base Substitutions

E

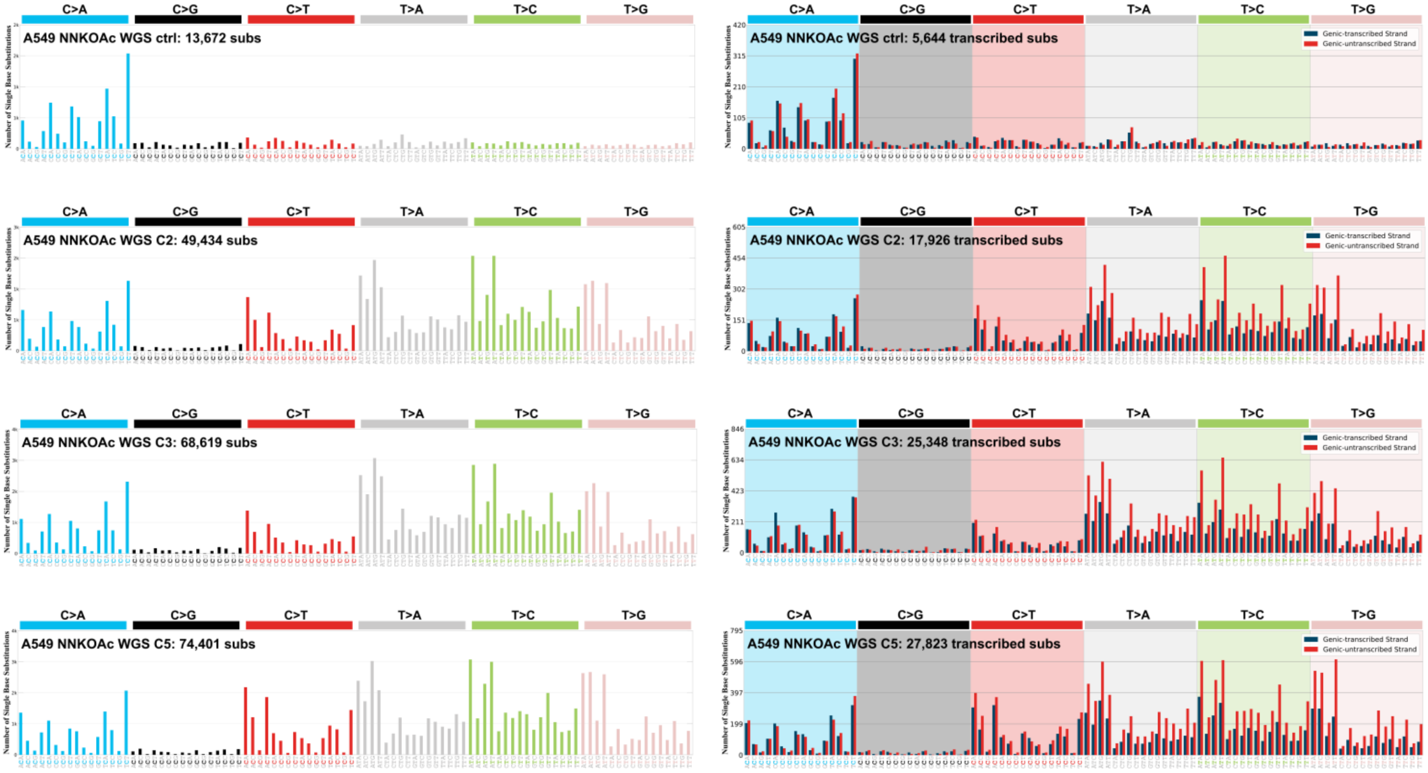
C>A
C>G
C>T
T>A
T>C
T>G
A549 NNKOAc WGS ctrl: 13,672 subs
A549 NNKOAc WGS ctrl: 5,644 transcribed subs
A549 NNKOAc WGS C2: 49,434 subs
A549 NNKOAc WGS C2: 17,926 transcribed subs
A549 NNKOAc WGS C3: 68,619 subs
A549 NNKOAc WGS C3: 25,348 transcribed subs
A549 NNKOAc WGS C5: 74,401 subs
A549 NNKOAc WGS C5: 27,823 transcribed subs
Number of Single Base Substitutions
Genic-transcribed Strand
Genic-untranscribed Strand

F

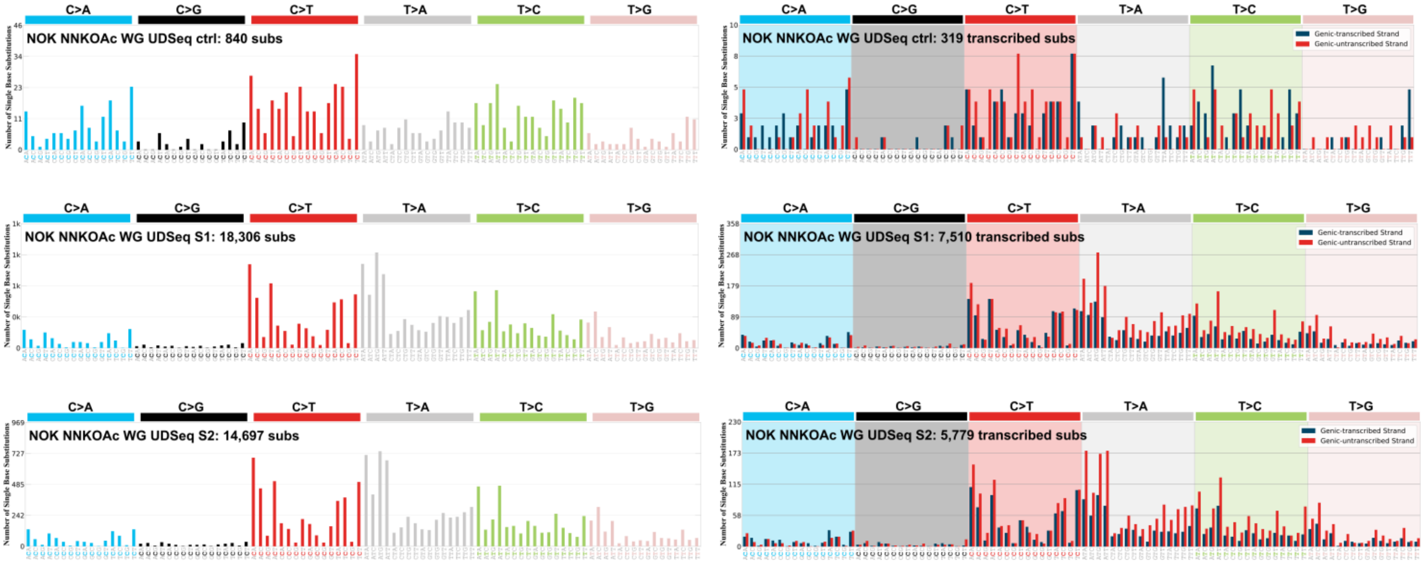

C>A
C>G
C>T
T>A
T>C
T>G
NOK NNKOAc WG UDSeq ctrl: 840 subs
NOK NNKOAc WG UDSeq ctrl: 319 transcribed subs
Genic-transcribed Strand
Genic-untranscribed Strand
NOK NNKOAc WG UDSeq S1: 18,306 subs
NOK NNKOAc WG UDSeq S1: 7,510 transcribed subs
NOK NNKOAc WG UDSeq S2: 14,697 subs
NOK NNKOAc WG UDSeq S2: 5,779 transcribed subs
Number of Single Base Substitutions

G

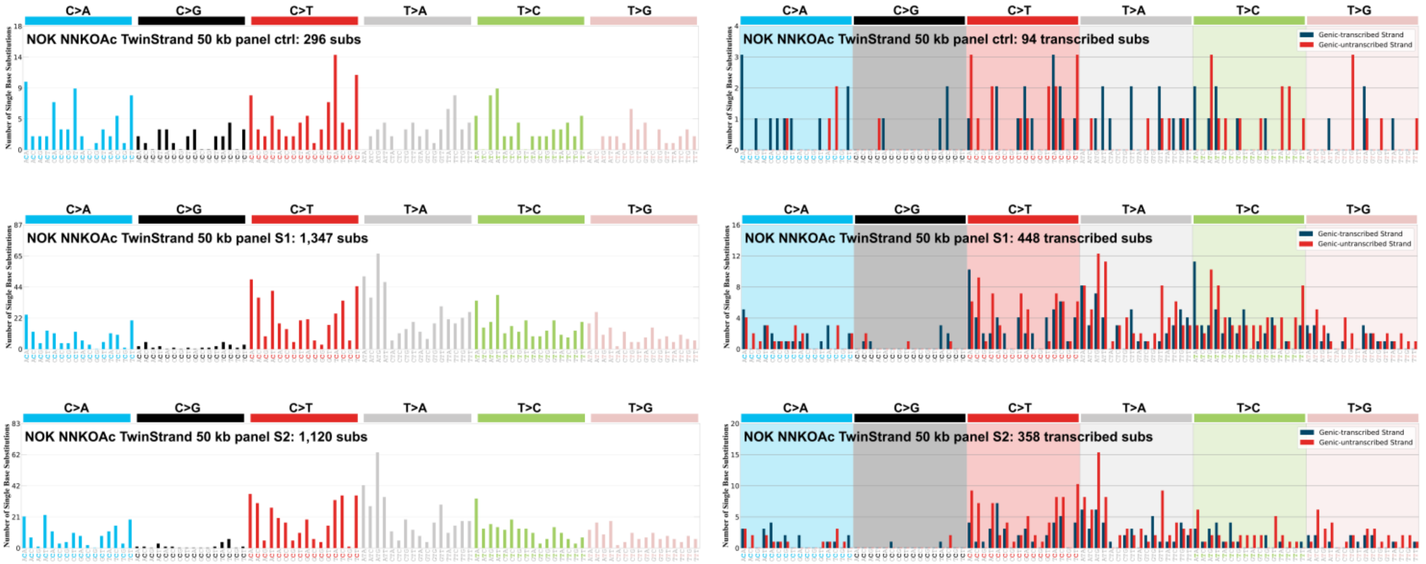

C>A
C>G
C>T
T>A
T>C
T>G
NOK NNKOAc TwinStrand 50 kb panel ctrl: 296 subs
NOK NNKOAc TwinStrand 50 kb panel ctrl: 94 transcribed subs
Genic-transcribed Strand
Genic-untranscribed Strand
NOK NNKOAc TwinStrand 50 kb panel S1: 1,347 subs
NOK NNKOAc TwinStrand 50 kb panel S1: 448 transcribed subs
NOK NNKOAc TwinStrand 50 kb panel S2: 1,120 subs
NOK NNKOAc TwinStrand 50 kb panel S2: 358 transcribed subs

Supplementary Figure 9

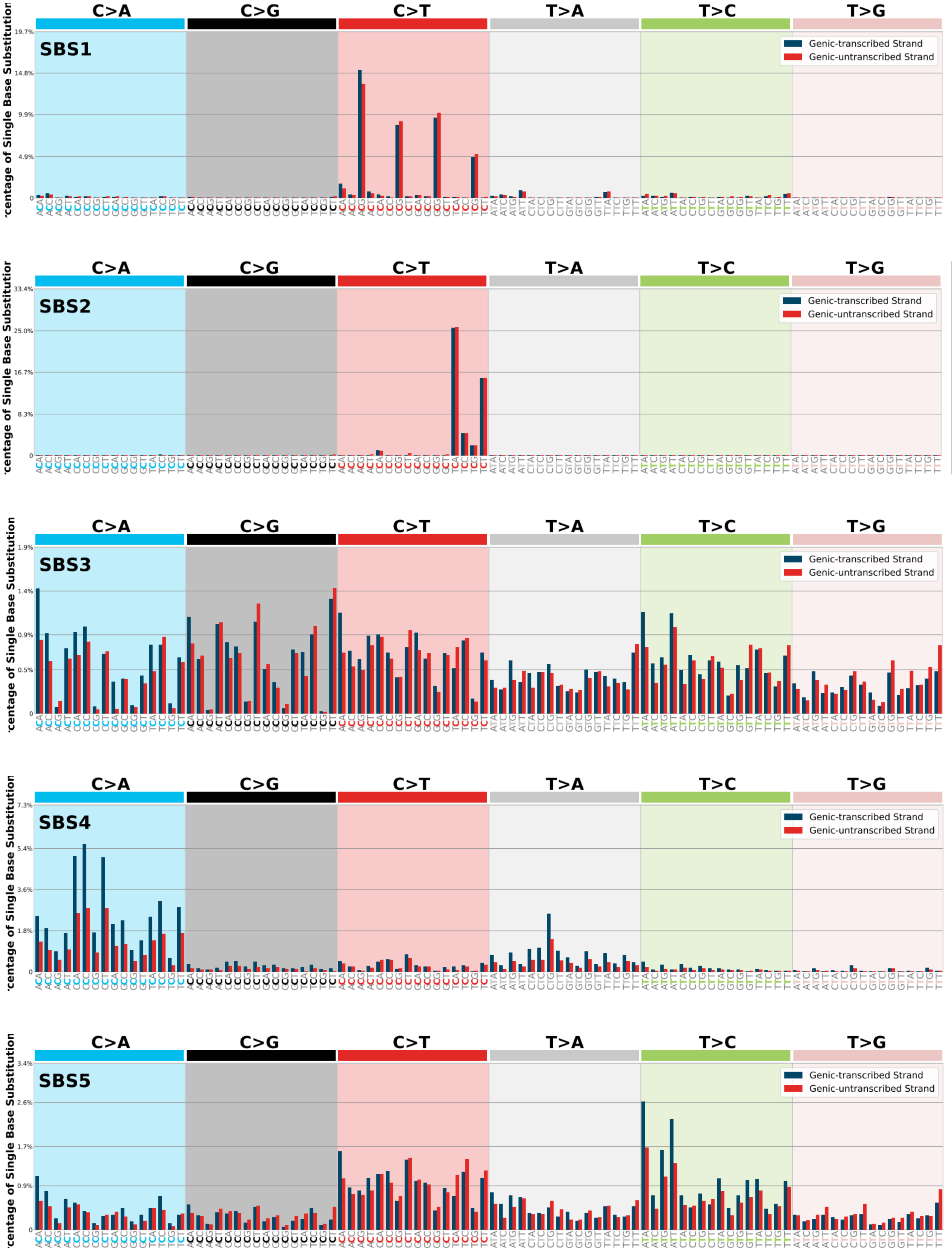

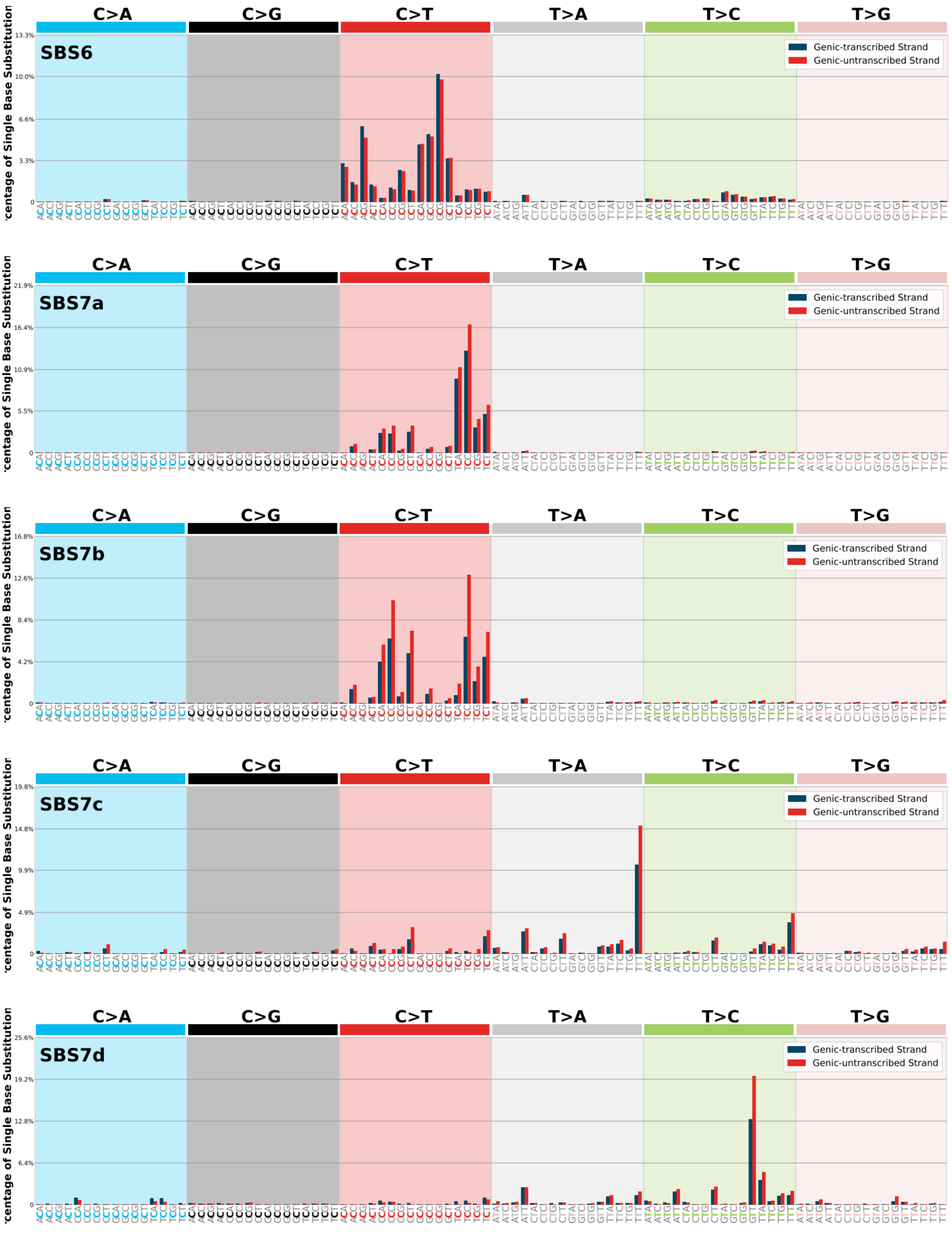
C>A
C>G
C>T
T>A
T>C
T>G
SBS6
Genic-transcribed Strand
Genic-untranscribed Strand
SBS7a
SBS7b
SBS7c
SBS7d

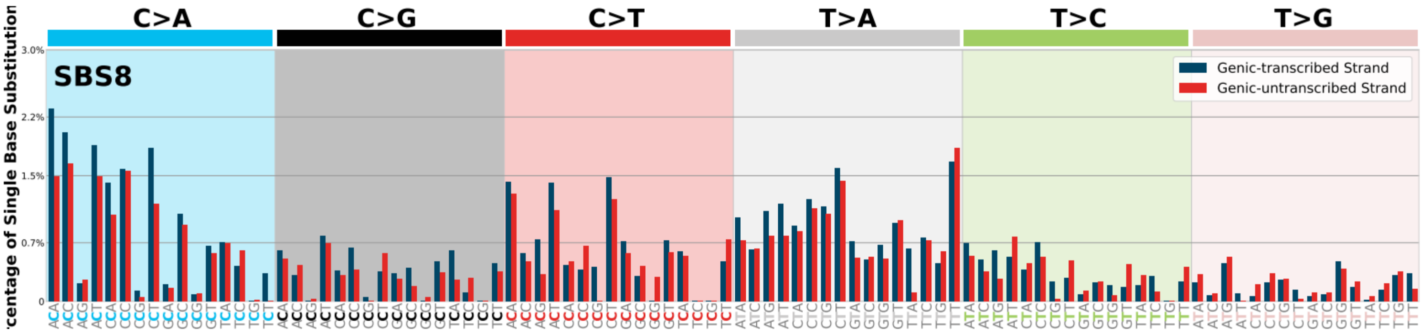
SBS8
C>A
C>G
C>T
T>A
T>C
T>G
Genic-transcribed Strand
Genic-untranscribed Strand
centage of Single Base Substitution
3.0%
2.2%
1.5%
0.7%
0

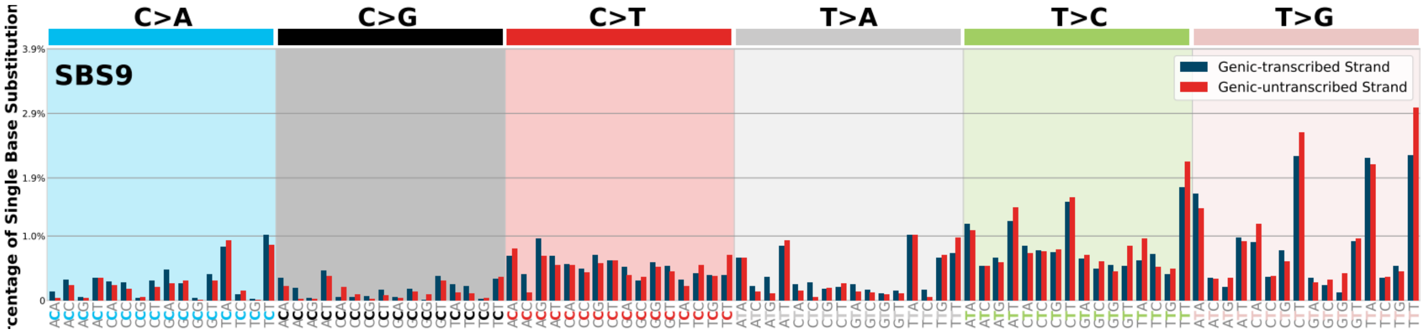
SBS9
C>A
C>G
C>T
T>A
T>C
T>G
Genic-transcribed Strand
Genic-untranscribed Strand
centage of Single Base Substitution
3.9%
2.9%
1.9%
1.0%
0

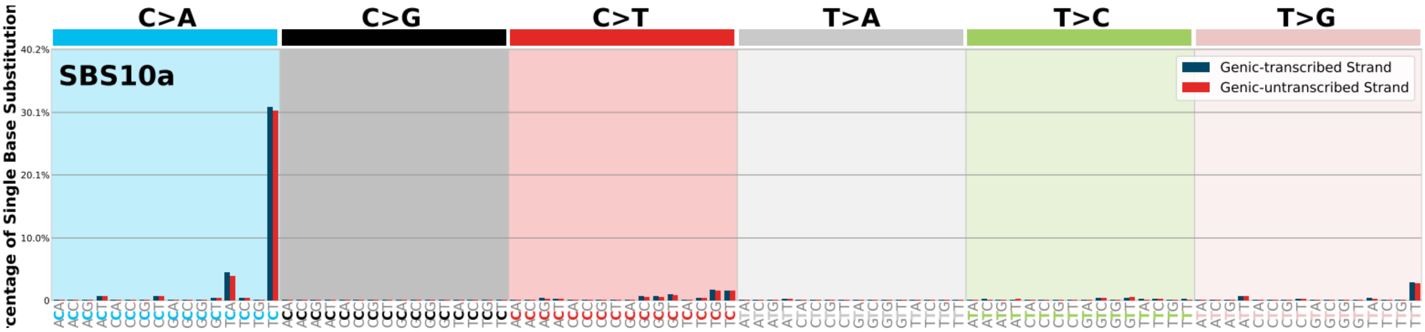
SBS10a
C>A
C>G
C>T
T>A
T>C
T>G
Genic-transcribed Strand
Genic-untranscribed Strand
centage of Single Base Substitution
40.2%
30.1%
20.1%
10.0%
0

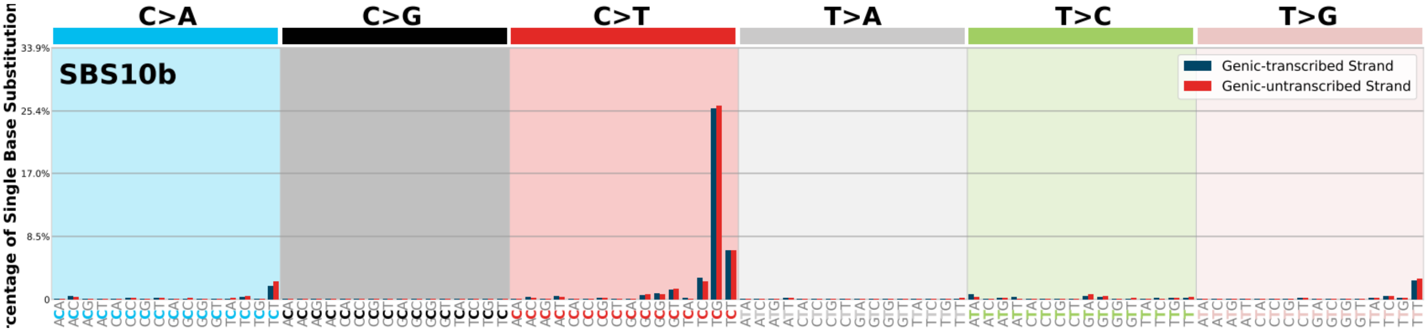
SBS10b
C>A
C>G
C>T
T>A
T>C
T>G
Genic-transcribed Strand
Genic-untranscribed Strand
centage of Single Base Substitution
33.9%
25.4%
17.0%
8.5%
0

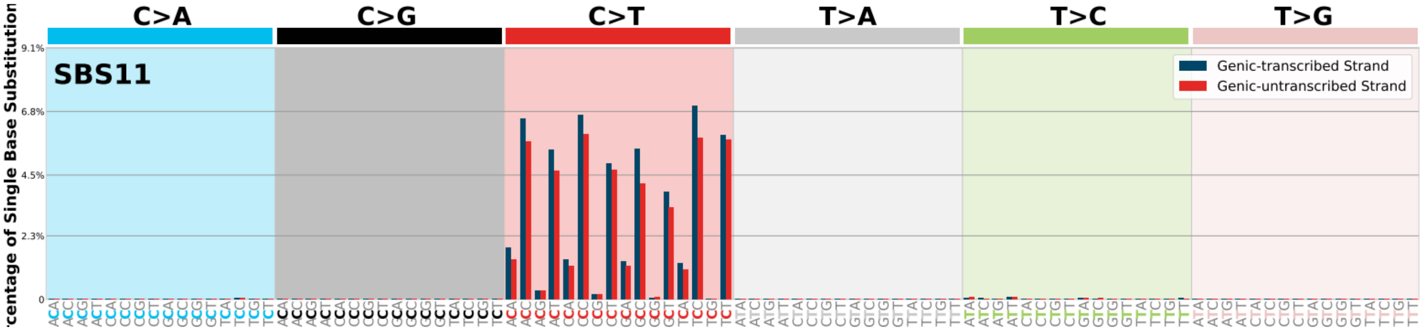
SBS11
C>A
C>G
C>T
T>A
T>C
T>G
Genic-transcribed Strand
Genic-untranscribed Strand
centage of Single Base Substitution
9.1%
6.8%
4.5%
2.3%
0

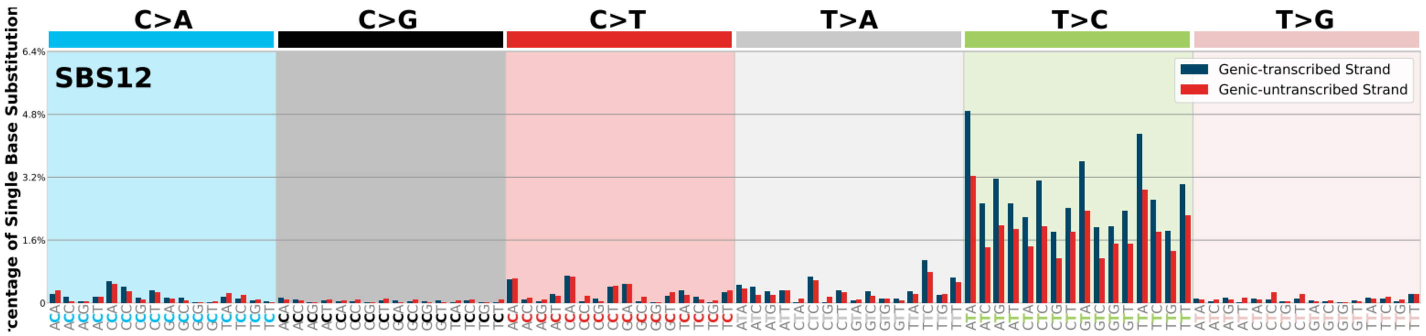

SBS12
C>A
C>G
C>T
T>A
T>C
T>G
Genic-transcribed Strand
Genic-untranscribed Strand
centage of Single Base Substitution
6.4%
4.8%
3.2%
1.6%
0


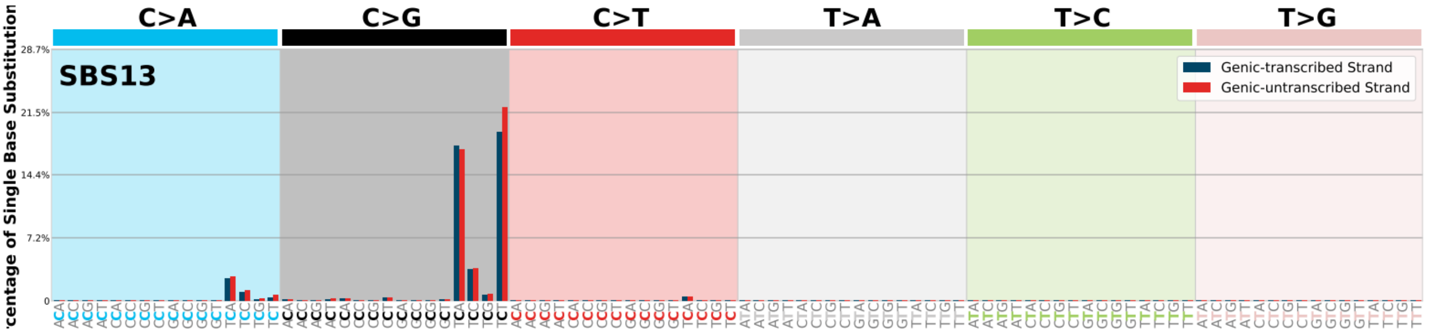

SBS13
C>A
C>G
C>T
T>A
T>C
T>G
Genic-transcribed Strand
Genic-untranscribed Strand
centage of Single Base Substitution
28.7%
21.5%
14.4%
7.2%
0


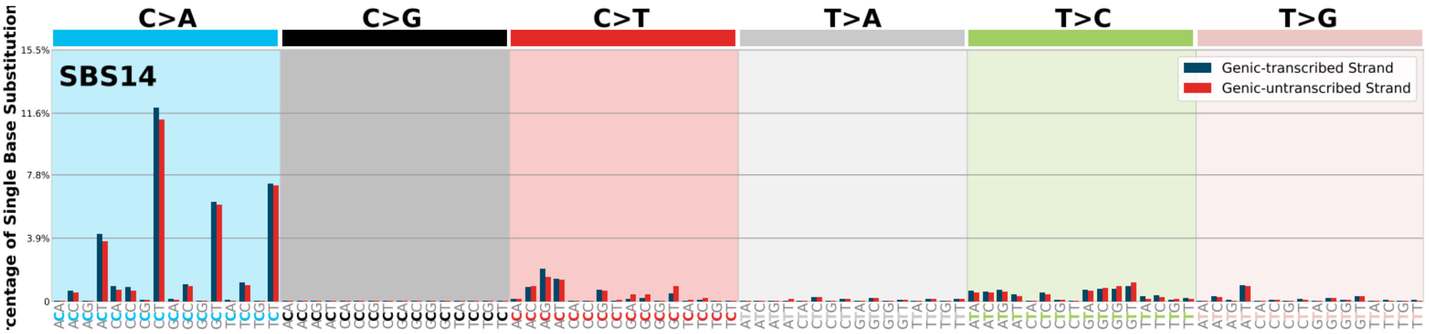

SBS14
C>A
C>G
C>T
T>A
T>C
T>G
Genic-transcribed Strand
Genic-untranscribed Strand
centage of Single Base Substitution
15.5%
11.6%
7.8%
3.9%
0


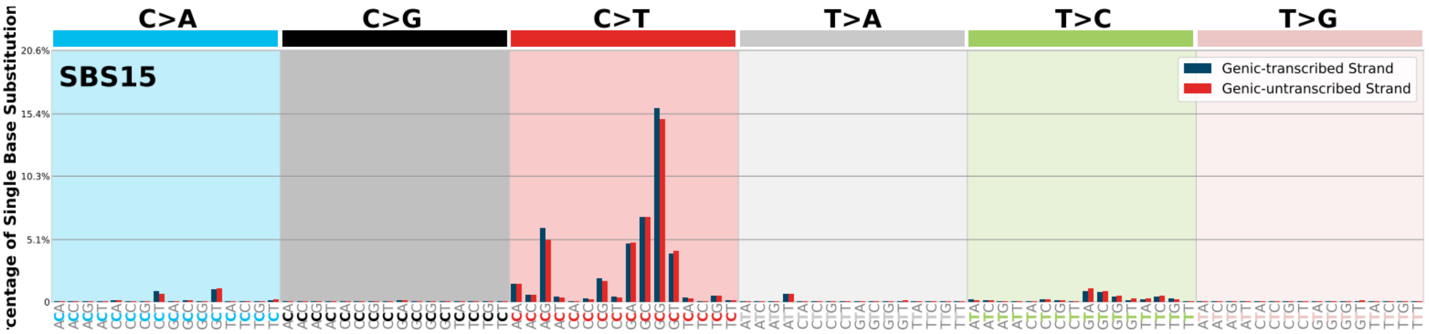

SBS15
C>A
C>G
C>T
T>A
T>C
T>G
Genic-transcribed Strand
Genic-untranscribed Strand
centage of Single Base Substitution
20.6%
15.4%
10.3%
5.1%
0


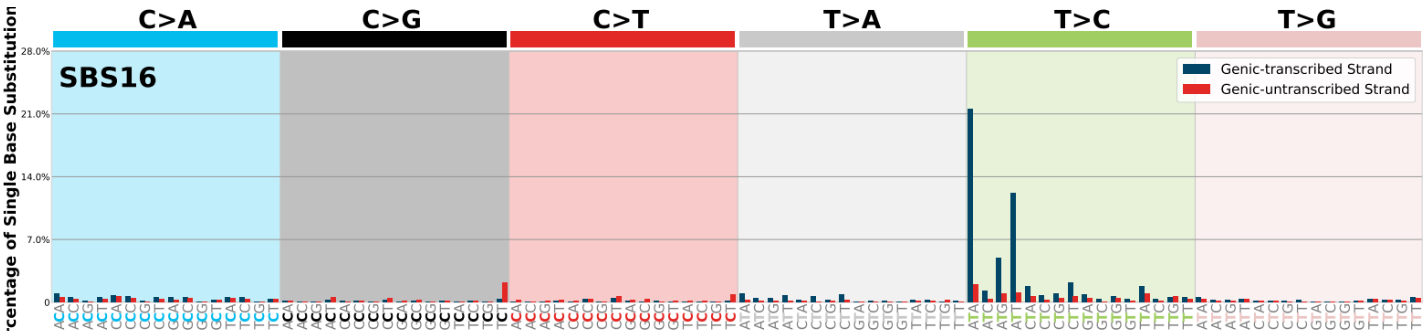

SBS16
C>A
C>G
C>T
T>A
T>C
T>G
Genic-transcribed Strand
Genic-untranscribed Strand
centage of Single Base Substitution
28.0%
21.0%
14.0%
7.0%
0

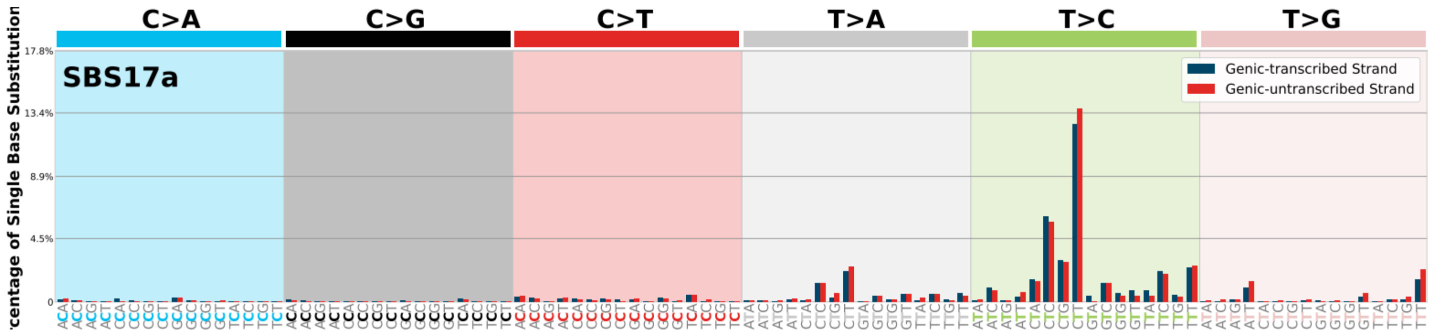
C>A
C>G
C>T
T>A
T>C
T>G
SBS17a
Genic-transcribed Strand
Genic-untranscribed Strand

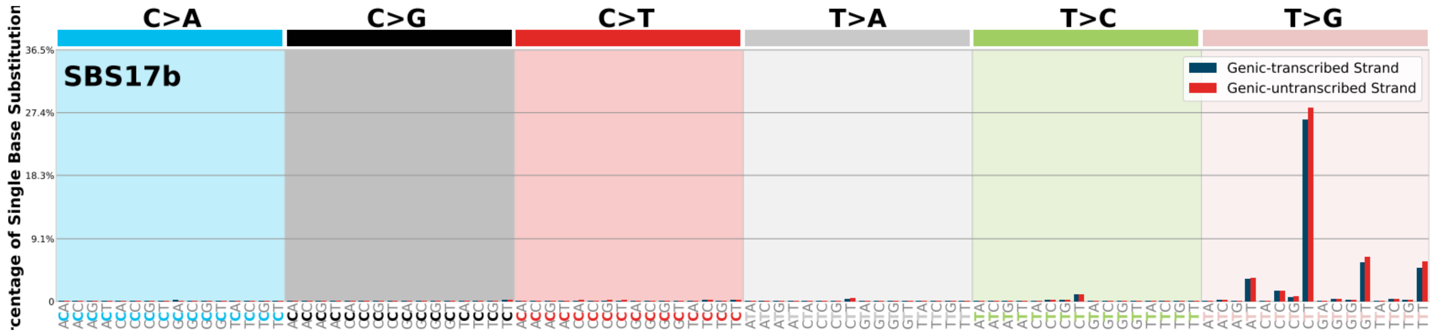
C>A
C>G
C>T
T>A
T>C
T>G
SBS17b
Genic-transcribed Strand
Genic-untranscribed Strand

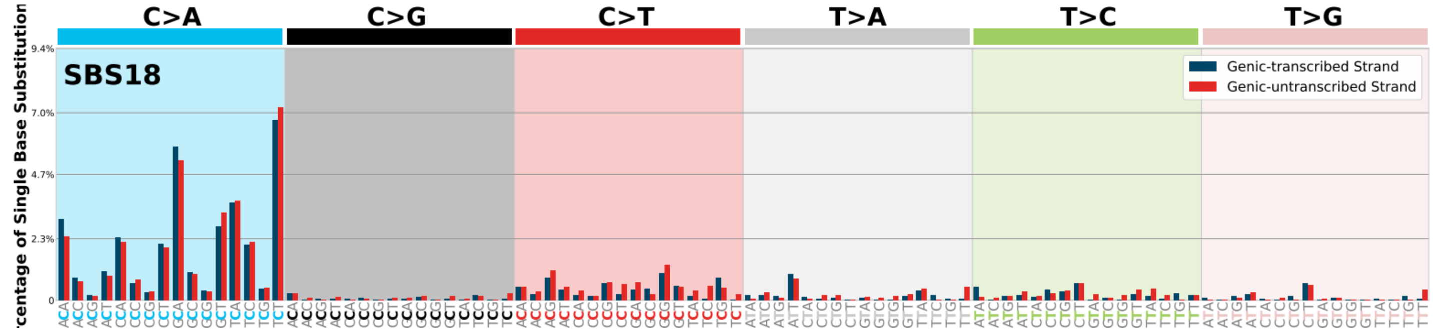
C>A
C>G
C>T
T>A
T>C
T>G
SBS18
Genic-transcribed Strand
Genic-untranscribed Strand

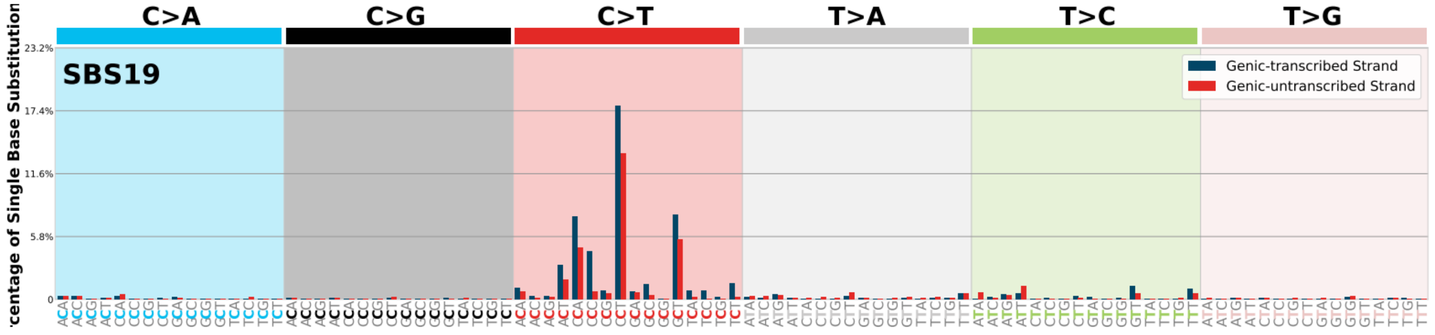
C>A
C>G
C>T
T>A
T>C
T>G
SBS19
Genic-transcribed Strand
Genic-untranscribed Strand

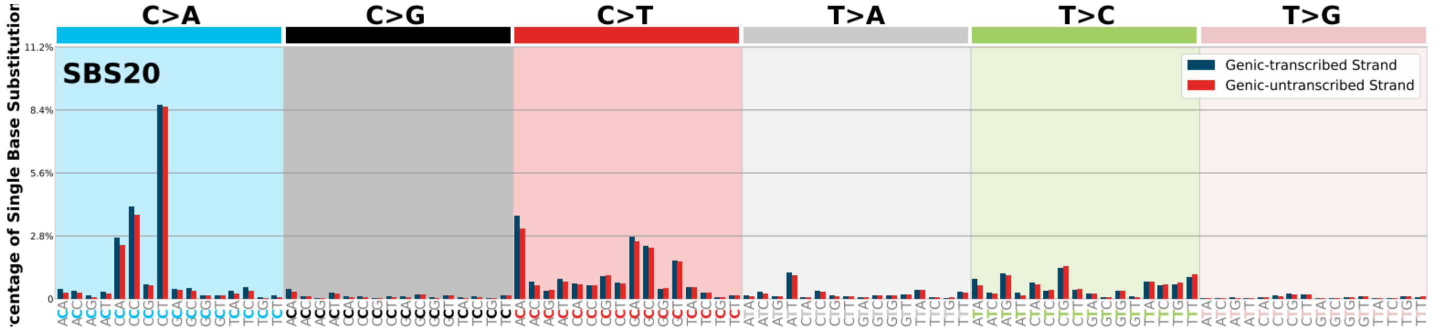
C>A
C>G
C>T
T>A
T>C
T>G
SBS20
Genic-transcribed Strand
Genic-untranscribed Strand

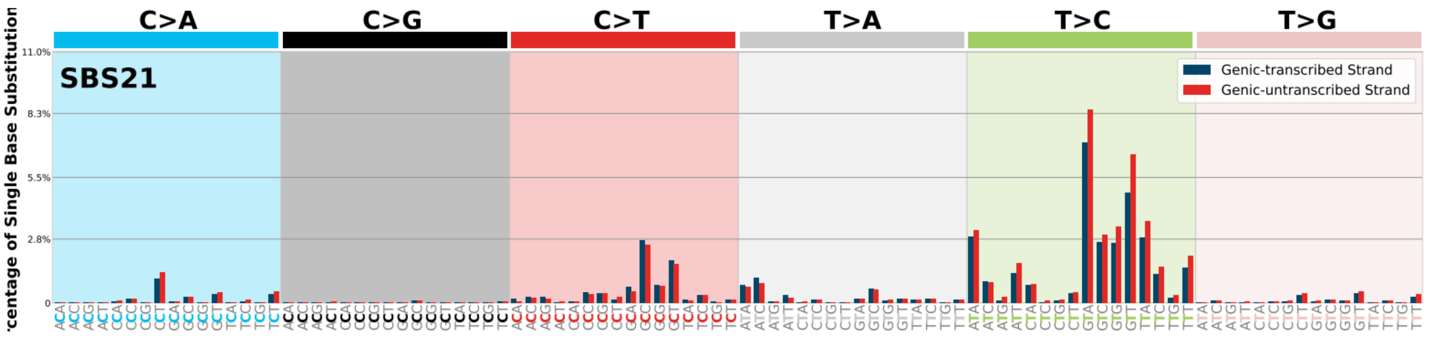
SBS21
C>A
C>G
C>T
T>A
T>C
T>G
Genic-transcribed Strand
Genic-untranscribed Strand
11.0%
8.3%
5.5%
2.8%
0

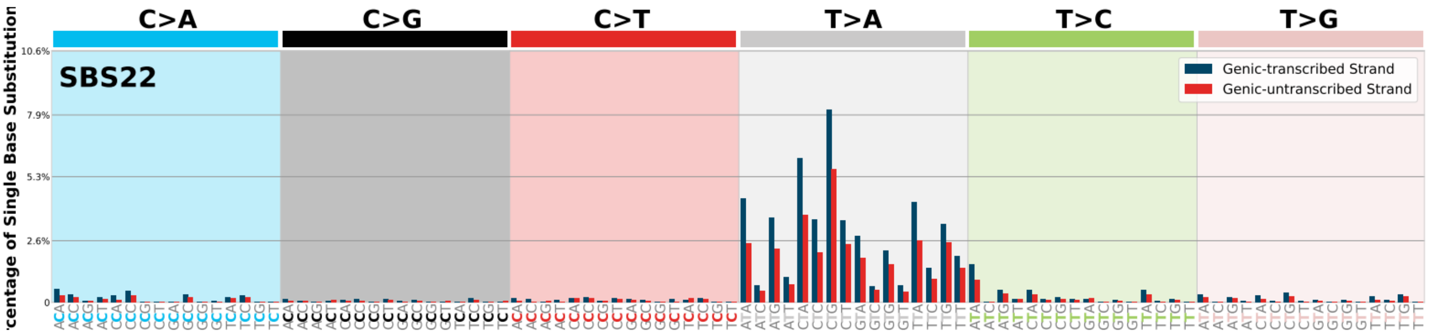
SBS22
C>A
C>G
C>T
T>A
T>C
T>G
Genic-transcribed Strand
Genic-untranscribed Strand
10.6%
7.9%
5.3%
2.6%
0

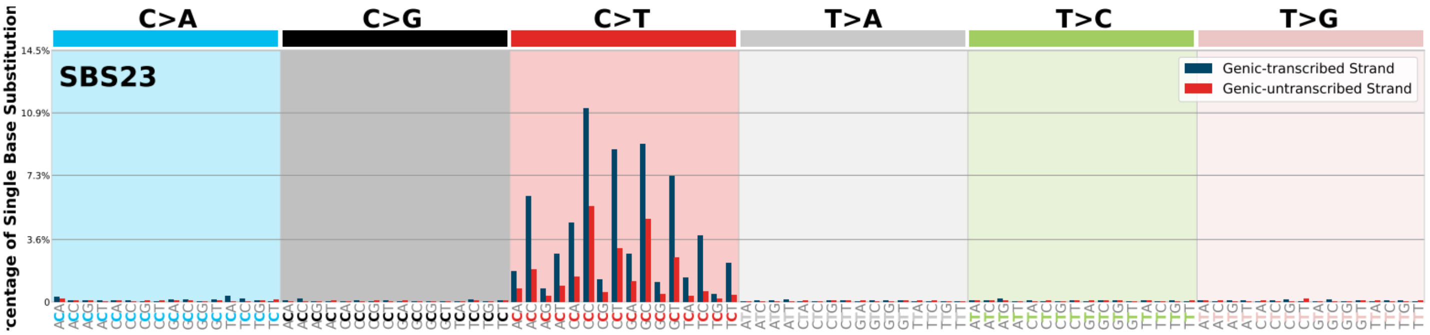
SBS23
C>A
C>G
C>T
T>A
T>C
T>G
Genic-transcribed Strand
Genic-untranscribed Strand
14.5%
10.9%
7.3%
3.6%
0

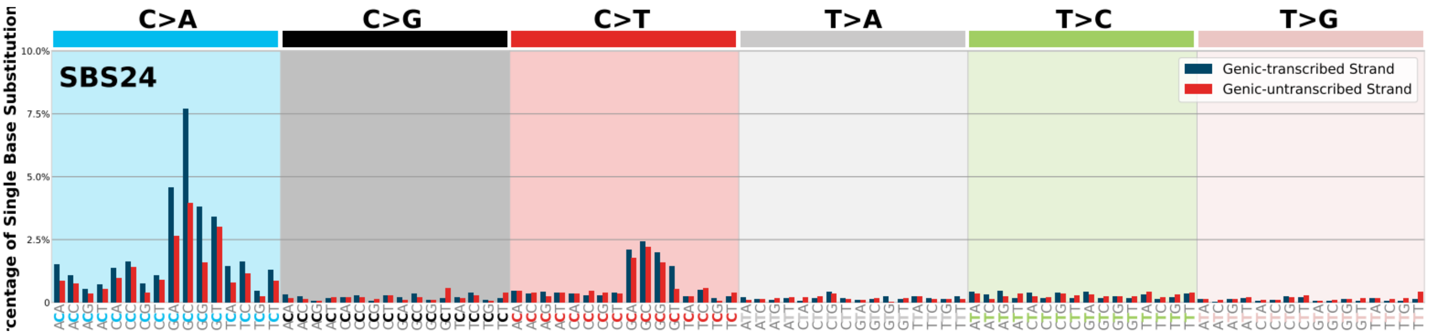
SBS24
C>A
C>G
C>T
T>A
T>C
T>G
Genic-transcribed Strand
Genic-untranscribed Strand
10.0%
7.5%
5.0%
2.5%
0

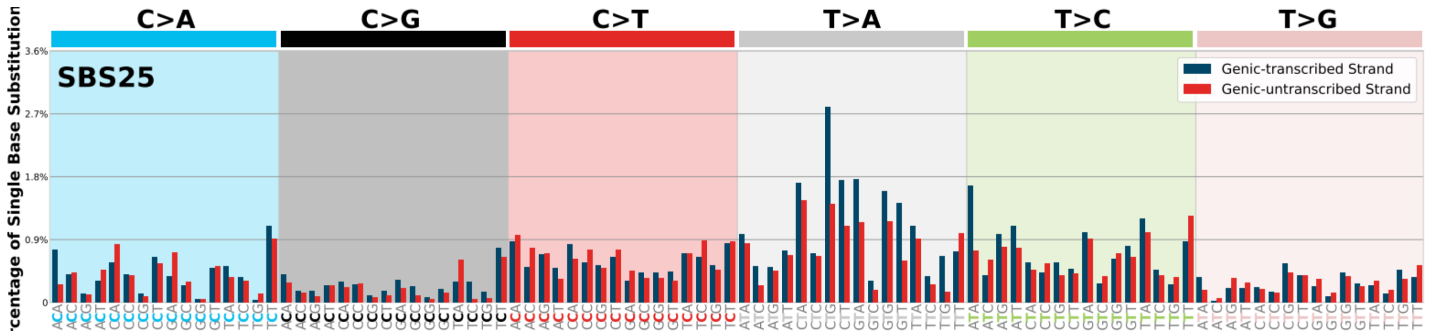
SBS25
C>A
C>G
C>T
T>A
T>C
T>G
Genic-transcribed Strand
Genic-untranscribed Strand
3.6%
2.7%
1.8%
0.9%
0

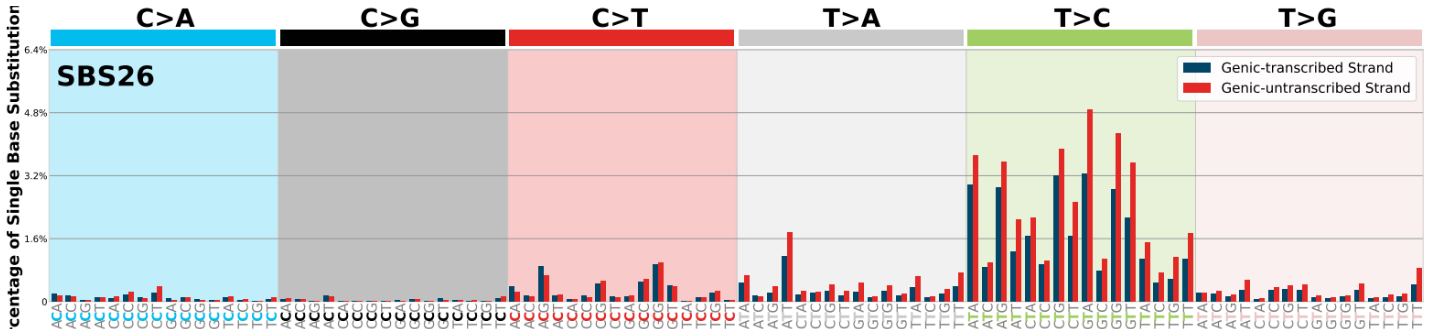
C>A
C>G
C>T
T>A
T>C
T>G
SBS26
Genic-transcribed Strand
Genic-untranscribed Strand
centage of Single Base Substitution
6.4%
4.8%
3.2%
1.6%
0

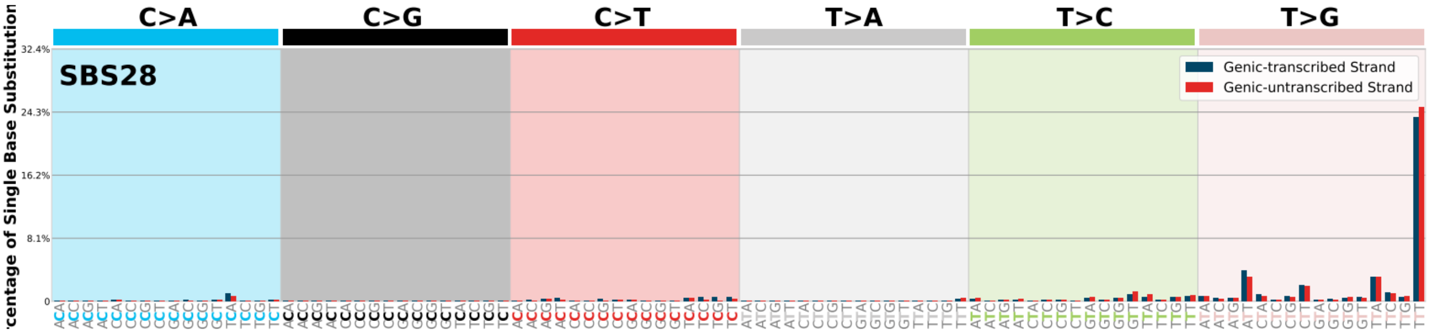
C>A
C>G
C>T
T>A
T>C
T>G
SBS28
Genic-transcribed Strand
Genic-untranscribed Strand
centage of Single Base Substitution
32.4%
24.3%
16.2%
8.1%
0

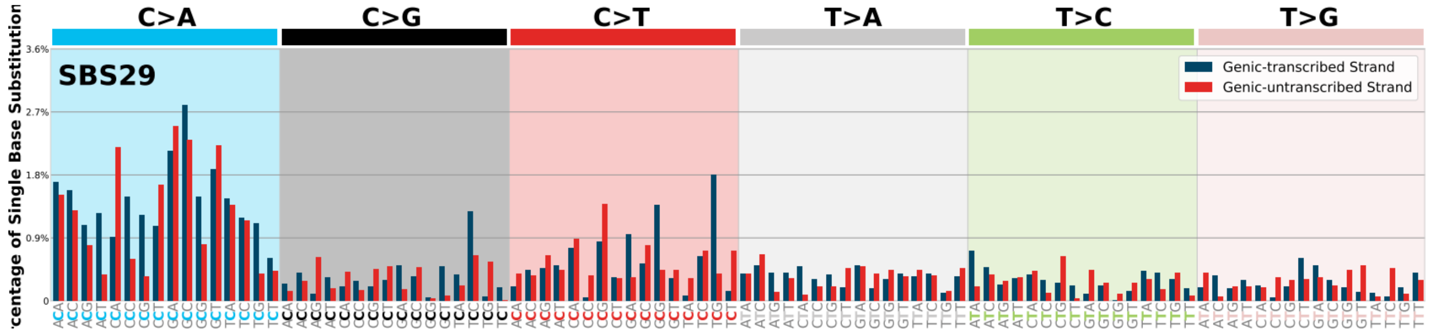
C>A
C>G
C>T
T>A
T>C
T>G
SBS29
Genic-transcribed Strand
Genic-untranscribed Strand
centage of Single Base Substitution
3.6%
2.7%
1.8%
0.9%
0

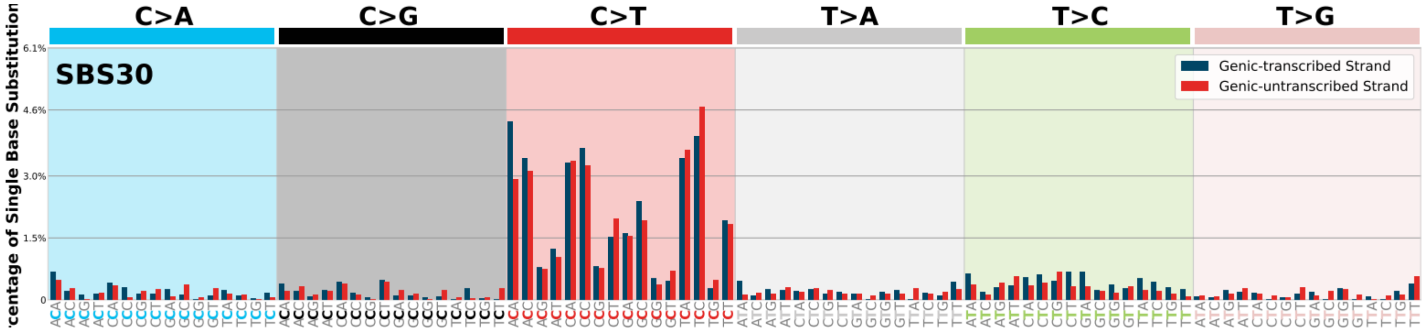
C>A
C>G
C>T
T>A
T>C
T>G
SBS30
Genic-transcribed Strand
Genic-untranscribed Strand
centage of Single Base Substitution
6.1%
4.6%
3.0%
1.5%
0

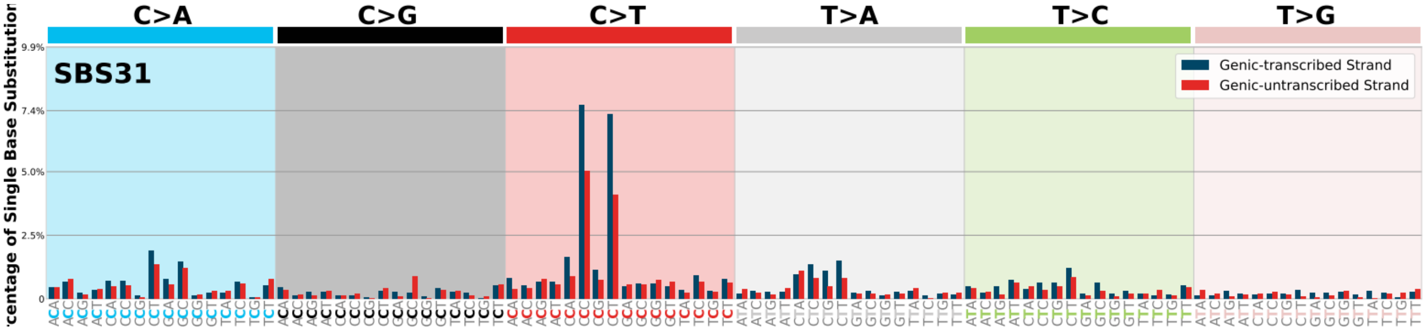
C>A
C>G
C>T
T>A
T>C
T>G
SBS31
Genic-transcribed Strand
Genic-untranscribed Strand
centage of Single Base Substitution
9.9%
7.4%
5.0%
2.5%
0

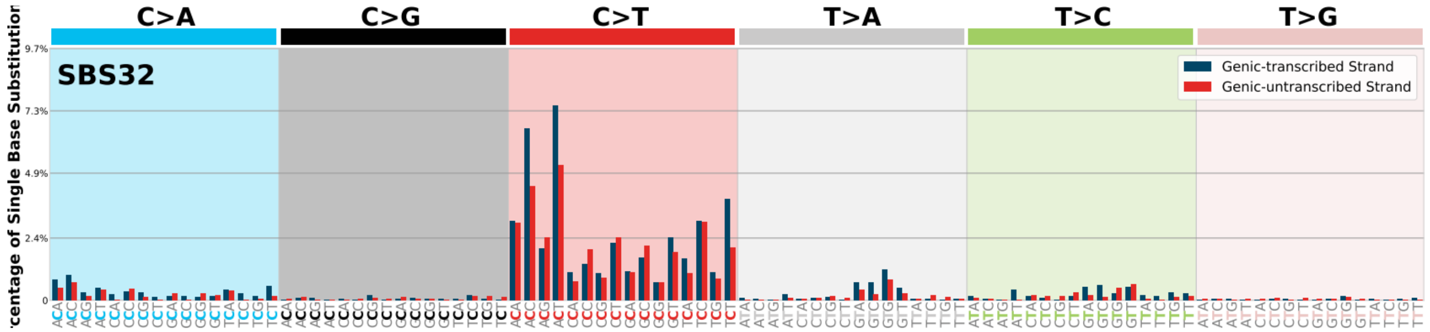
C>A
C>G
C>T
T>A
T>C
T>G
SBS32
Genic-transcribed Strand
Genic-untranscribed Strand
centage of Single Base Substitution
9.7%
7.3%
4.9%
2.4%
0

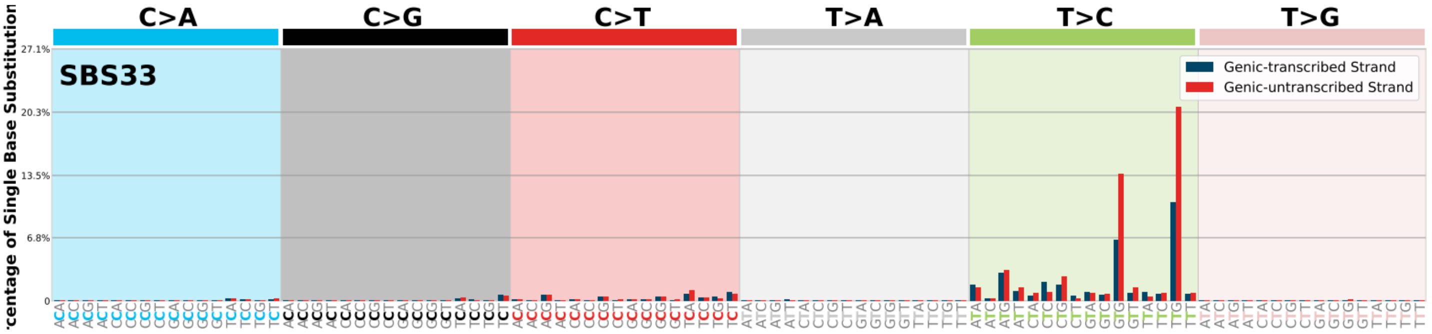
C>A
C>G
C>T
T>A
T>C
T>G
SBS33
Genic-transcribed Strand
Genic-untranscribed Strand
centage of Single Base Substitution
27.1%
20.3%
13.5%
6.8%
0

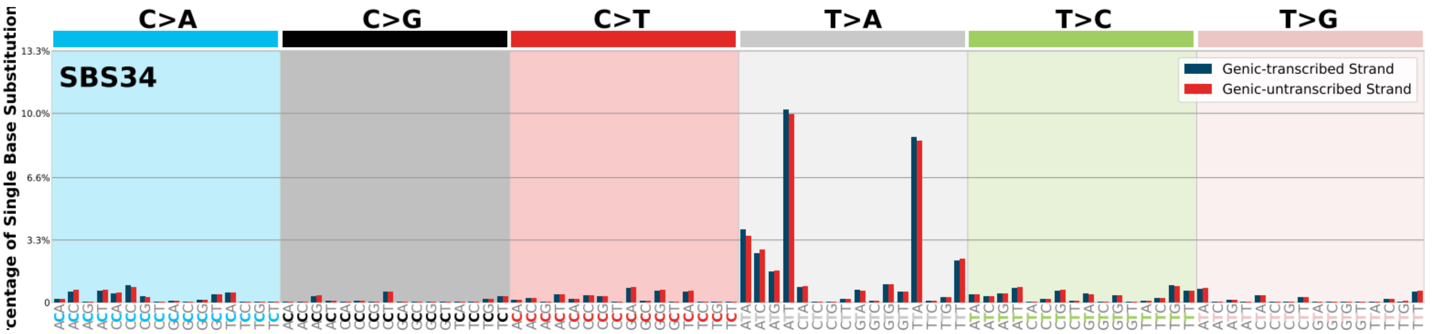
C>A
C>G
C>T
T>A
T>C
T>G
SBS34
Genic-transcribed Strand
Genic-untranscribed Strand
centage of Single Base Substitution
13.3%
10.0%
6.6%
3.3%
0

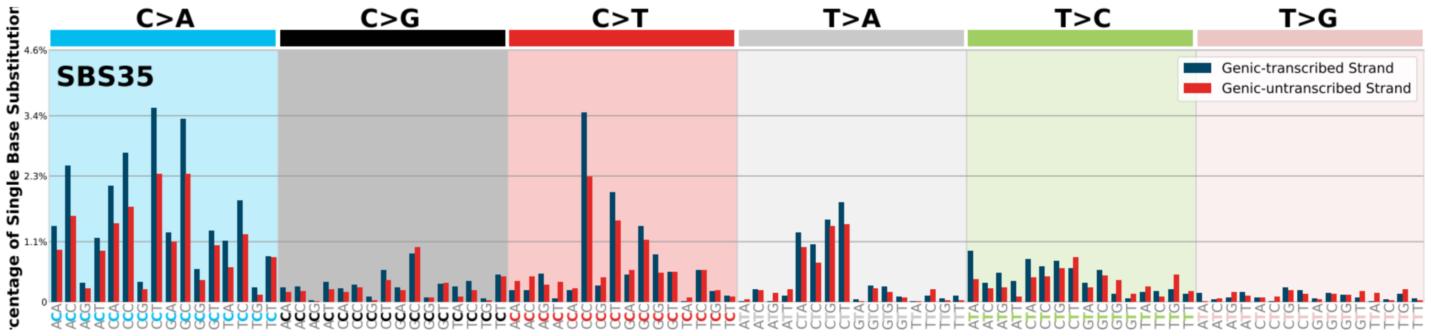
C>A
C>G
C>T
T>A
T>C
T>G
SBS35
Genic-transcribed Strand
Genic-untranscribed Strand
centage of Single Base Substitution
4.6%
3.4%
2.3%
1.1%
0

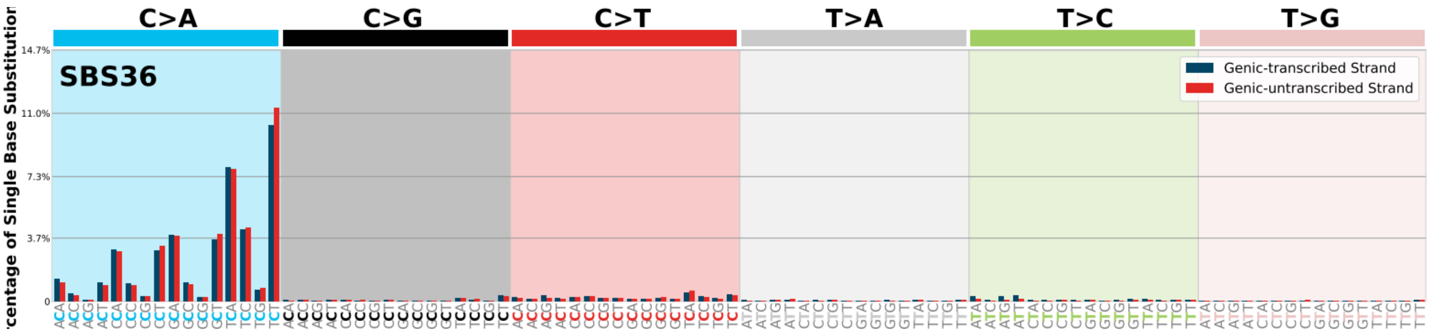
C>A
C>G
C>T
T>A
T>C
T>G
SBS36
Genic-transcribed Strand
Genic-untranscribed Strand
centage of Single Base Substitution
14.7%
11.0%
7.3%
3.7%
0

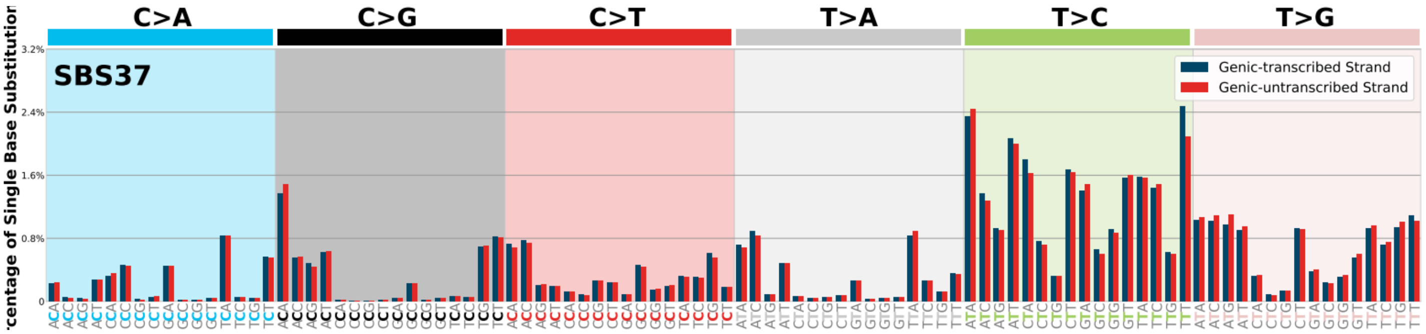
C>A
C>G
C>T
T>A
T>C
T>G
SBS37
Genic-transcribed Strand
Genic-untranscribed Strand
centage of Single Base Substitution
3.2%
2.4%
1.6%
0.8%
0

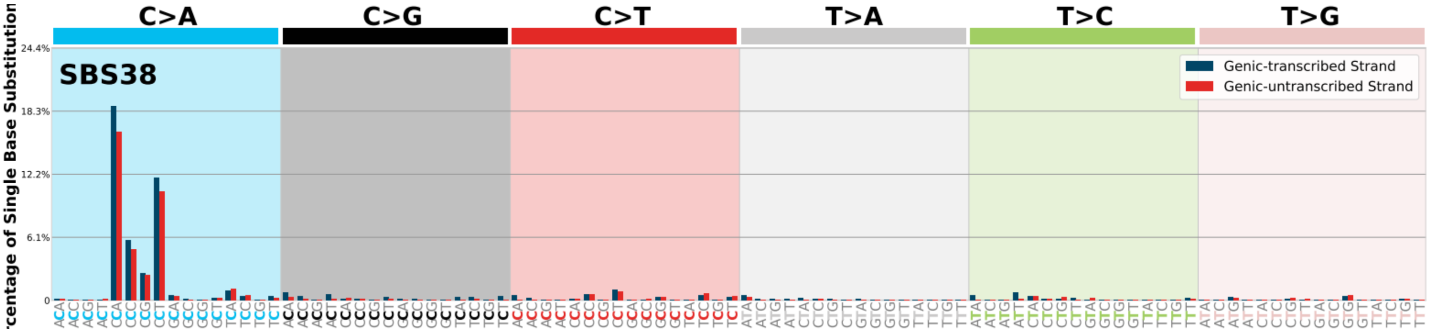
C>A
C>G
C>T
T>A
T>C
T>G
SBS38
Genic-transcribed Strand
Genic-untranscribed Strand
centage of Single Base Substitution
24.4%
18.3%
12.2%
6.1%
0

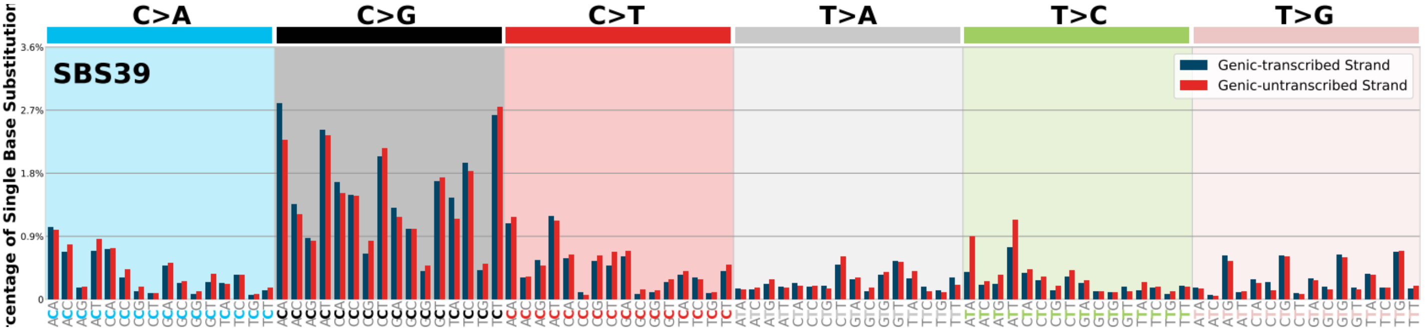
C>A
C>G
C>T
T>A
T>C
T>G
SBS39
Genic-transcribed Strand
Genic-untranscribed Strand
centage of Single Base Substitution
3.6%
2.7%
1.8%
0.9%
0

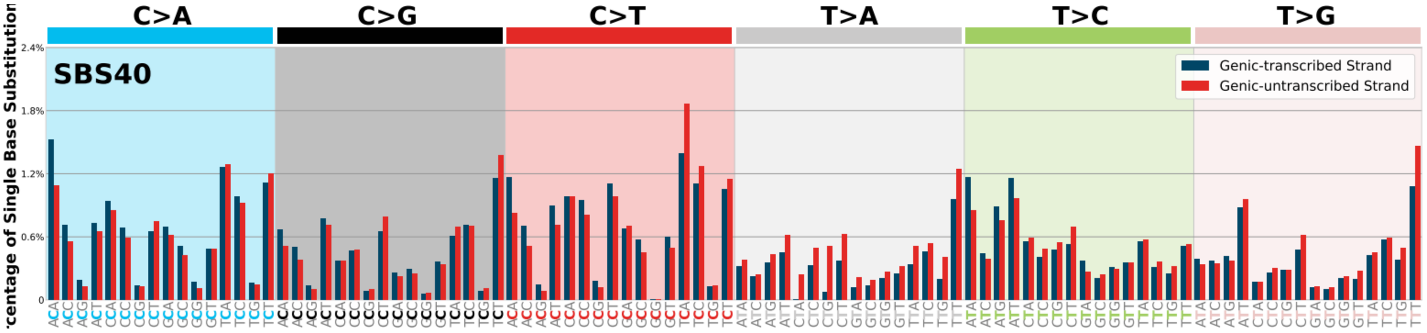
C>A
C>G
C>T
T>A
T>C
T>G
SBS40
Genic-transcribed Strand
Genic-untranscribed Strand
centage of Single Base Substitution
2.4%
1.8%
1.2%
0.6%
0

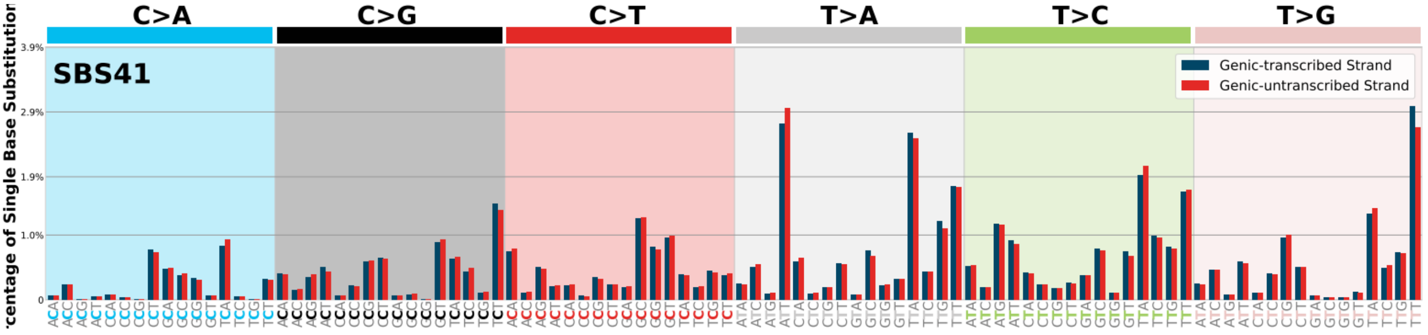
C>A
C>G
C>T
T>A
T>C
T>G
SBS41
Genic-transcribed Strand
Genic-untranscribed Strand
centage of Single Base Substitution
3.9%
2.9%
1.9%
1.0%
0

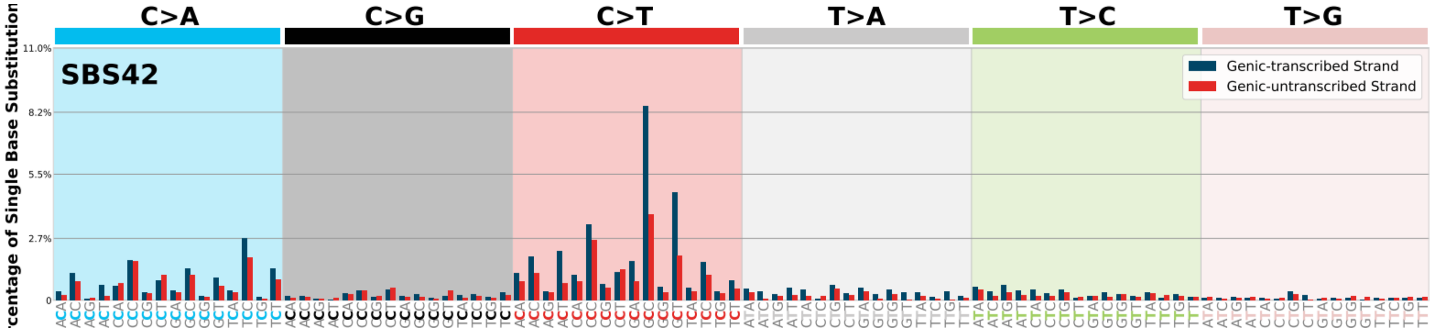
C>A
C>G
C>T
T>A
T>C
T>G
SBS42
Genic-transcribed Strand
Genic-untranscribed Strand

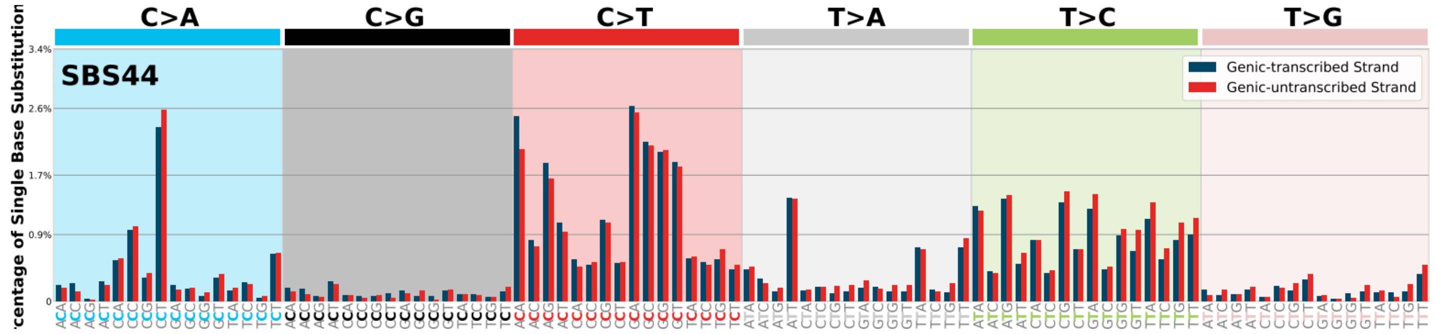
C>A
C>G
C>T
T>A
T>C
T>G
SBS44
Genic-transcribed Strand
Genic-untranscribed Strand

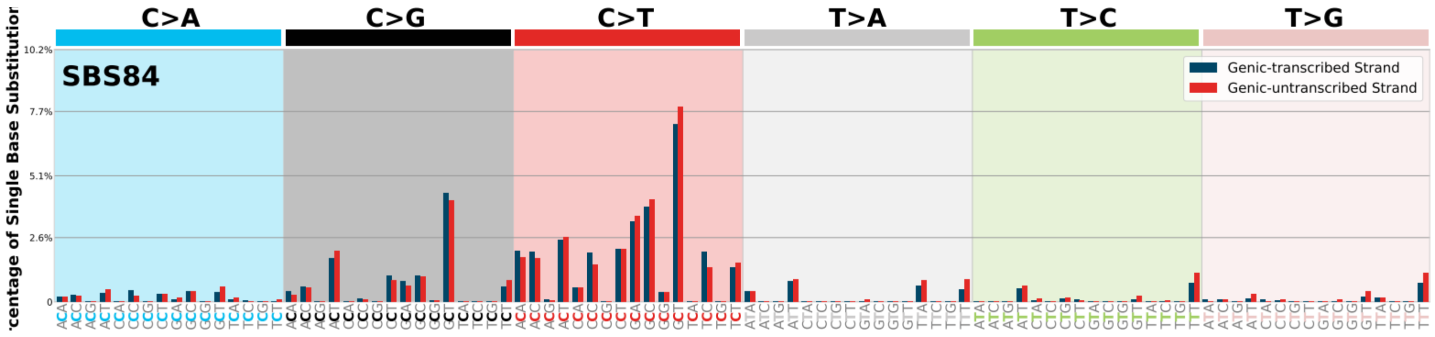
C>A
C>G
C>T
T>A
T>C
T>G
SBS84
Genic-transcribed Strand
Genic-untranscribed Strand

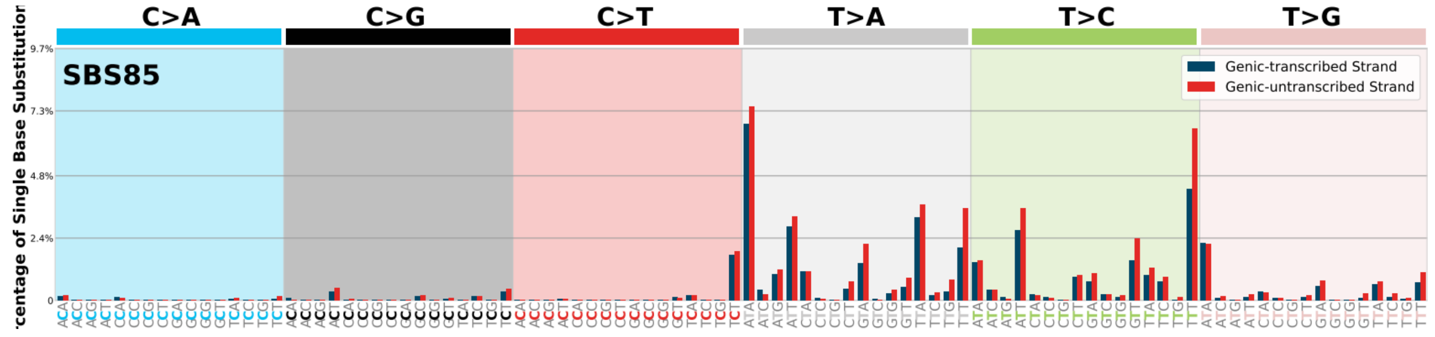
C>A
C>G
C>T
T>A
T>C
T>G
SBS85
Genic-transcribed Strand
Genic-untranscribed Strand

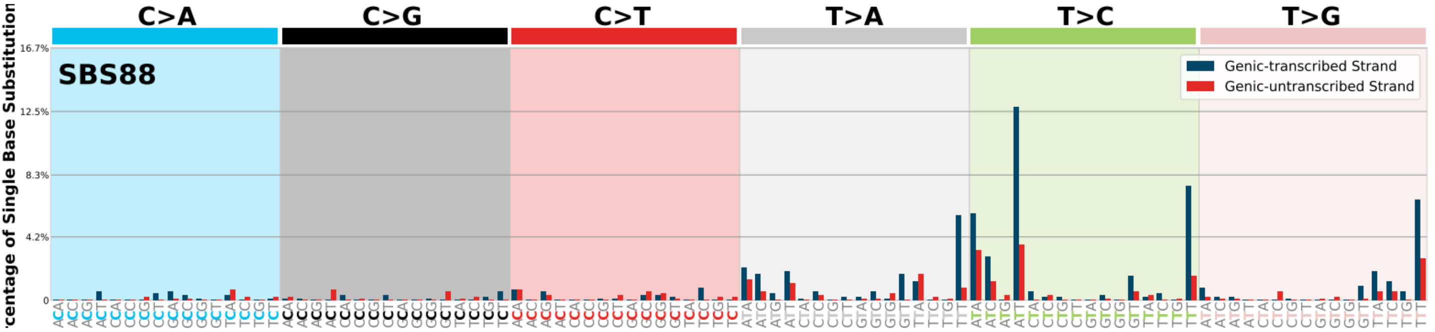
C>A
C>G
C>T
T>A
T>C
T>G
SBS88
Genic-transcribed Strand
Genic-untranscribed Strand

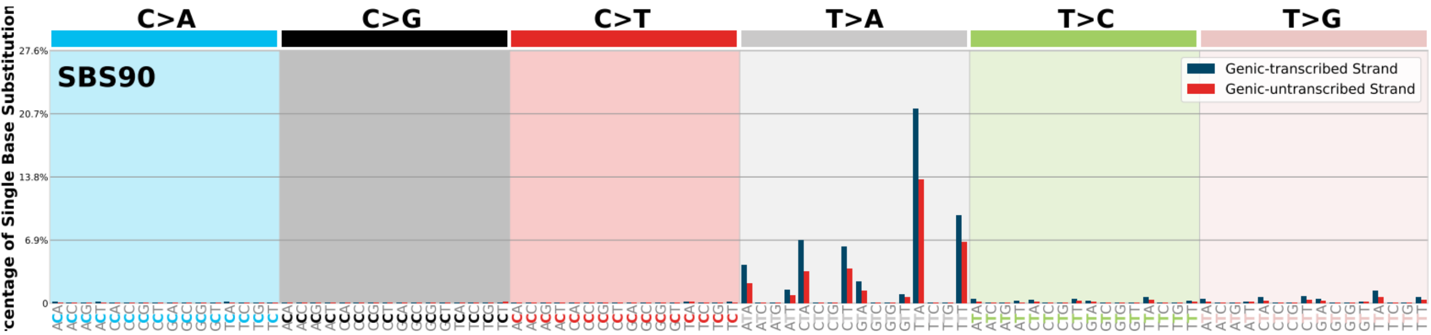
C>A
C>G
C>T
T>A
T>C
T>G
SBS90
Genic-transcribed Strand
Genic-untranscribed Strand
27.6%
20.7%
13.8%
6.9%
0
centage of Single Base Substitution

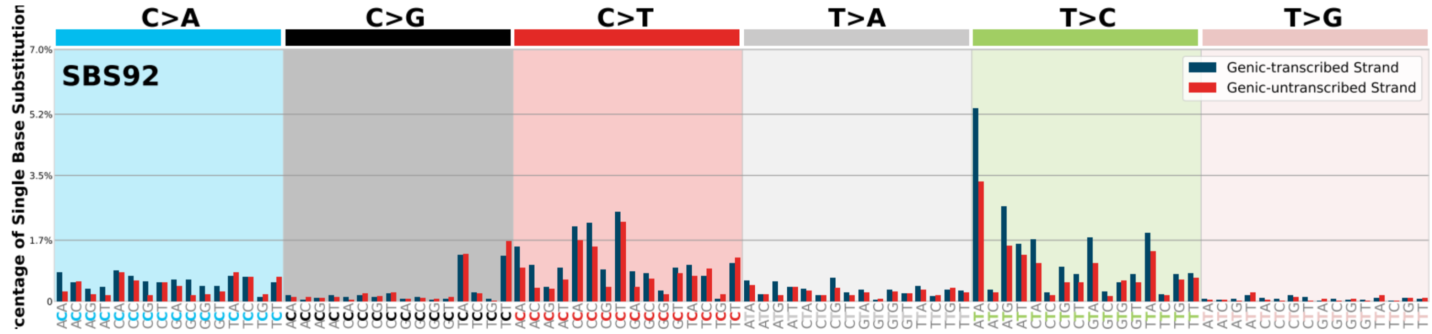
C>A
C>G
C>T
T>A
T>C
T>G
SBS92
Genic-transcribed Strand
Genic-untranscribed Strand
7.0%
5.2%
3.5%
1.7%
0
centage of Single Base Substitution

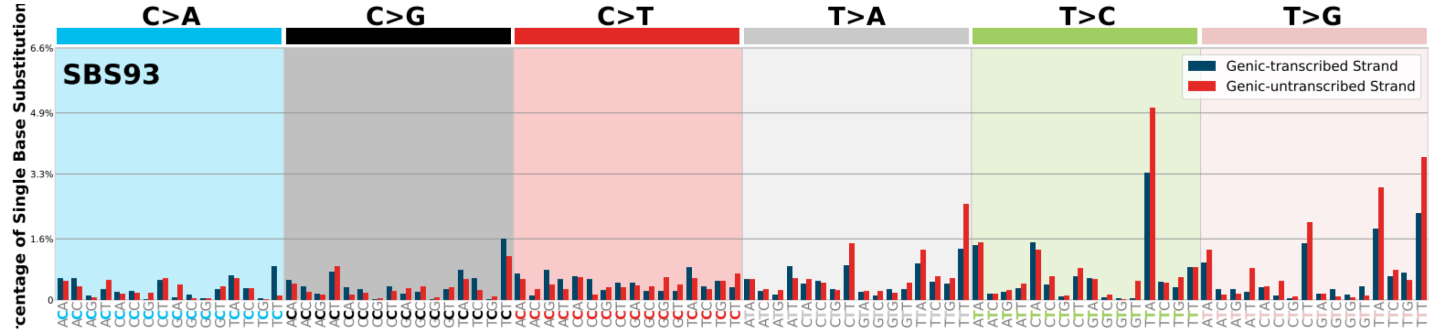
C>A
C>G
C>T
T>A
T>C
T>G
SBS93
Genic-transcribed Strand
Genic-untranscribed Strand
6.6%
4.9%
3.3%
1.6%
0
centage of Single Base Substitution

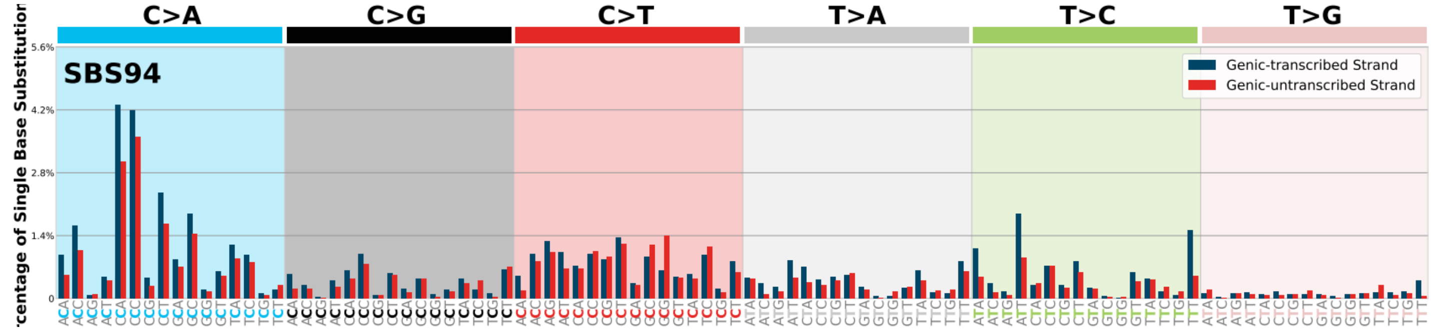
C>A
C>G
C>T
T>A
T>C
T>G
SBS94
Genic-transcribed Strand
Genic-untranscribed Strand
5.6%
4.2%
2.8%
1.4%
0
centage of Single Base Substitution
